# Targeting the Pregnane X Receptor Using Microbial Metabolite Mimicry

**DOI:** 10.1101/792671

**Authors:** Zdeněk Dvořák, Felix Kopp, Cait M. Costello, Jazmin S. Kemp, Hao Li, Aneta Vrzalová, Martina Štěpánková, Iveta Bartoňková, Eva Jiskrová, Karolína Poulíková, Barbora Vyhlídalová, Lars U. Nordstroem, Chamini Karunaratne, Harmit Ranhotra, Kyu Shik Mun, Anjaparavanda P. Naren, Iain Murray, Gary H. Perdew, Julius Brtko, Lucia Toporova, Arne Schon, William G. Wallace, William G. Walton, Matthew R. Redinbo, Katherine Sun, Amanda Beck, Sandhya Kortagere, Michelle C. Neary, Aneesh Chandran, Saraswathi Vishveshwara, Maria M. Cavalluzzi, Giovanni Lentini, Julia Yue Cui, Haiwei Gu, John C. March, Shirshendu Chaterjee, Adam Matson, Dennis Wright, Kyle L. Flannigan, Simon A. Hirota, R. Balfour Sartor, Sridhar Mani

## Abstract

The human pregnane X receptor (PXR), a master regulator of drug metabolism, has important roles in intestinal homeostasis and abrogating inflammation. Existing PXR ligands have substantial off-target toxicity. Based on prior work that established microbial (indole) metabolites as PXR ligands, we proposed microbial metabolite mimicry as a novel strategy for drug discovery that allows to exploit previously unexplored parts of chemical space. Here we report functionalized indole-derivatives as first-in-class non-cytotoxic PXR agonists, as a proof-of-concept for microbial metabolite mimicry. The lead compound, FKK6, binds directly to PXR protein in solution, induces PXR specific target gene expression in, cells, human organoids, and mice. FKK6 significantly represses pro-inflammatory cytokine production cells and abrogates inflammation in mice expressing the human PXR gene. The development of FKK6 demonstrates for the first time that microbial metabolite mimicry is a viable strategy for drug discovery and opens the door to mine underexploited regions of chemical space.

## Introduction

Microbial metabolite mimicry has been proposed as a means to probe areas of chemical space that have been previously underexploited for drug or probe discovery (RanhotraSaha et al., 2016). However, to-date, there has been no critical proof of this concept with regards to new drug discovery for treatment of human disease.

Our laboratory had previously demonstrated that bacterial metabolism of L-tryptophan results in the formation of indoles and indole-derived metabolites, such as indole 3-propionic acid (IPA), which are effective ligands for both mouse and human (h) pregnane X receptors (PXR), and can regulate host small intestinal immunity *via* the Toll-Like Receptor 4 (TLR4) pathway (Venkatesh, Mukherjee et al., 2014). The use of PXR as a therapeutic target for inflammatory bowel disease (IBD) is a compelling concept in both rodents and humans (Cheng, Shah et al., 2012b). While a variety of PXR-activating xenobiotics exist, they all have significant off-target effects in vivo (e.g., drug interactions) which limits their use as clinical drugs for IBD (Cheng, Krausz et al., 2012a).

Using a model developed in our laboratory (Venkatesh et al., 2014), we have shown that the co-occupancy of indole and indole 3-propionic acid (IPA) at the human PXR ligand-binding pocket and/or domain (LBD) is feasible. There are several opportunities to improve ligand binding *via* alteration of the pharmacophore H-bonding and pi-pi interactions (Venkatesh et al., 2014). This led us to propose that endogenous metabolite mimicry could yield more potent PXR ligands (agonists) but with limited toxicity since they would be largely derived from non-toxic metabolite-like subunits (Brave, Lukin et al., 2015, Mani, 2017). Further support for this concept comes from noting that basal indole levels in feces range in the low millimolar concentrations and these levels are relatively non-toxic to mammalian cells [LD_50_ oral in rats ∼ 1 g/kg, ThermoFisher Scientific Safety Sheet data] (Chappell, Darkoh et al., 2016). Similarly, indole metabolites like IPA, in feces, are likely to be present in low to high micromolar concentrations. Indeed, while these concepts have been proposed (Saha et al., 2016), there are very few, if any, clear examples of the utility of these concepts in drug or probe discovery.

The present work was aimed at providing the requisite proof that by developing mimics of microbial indoles using limited chemical analog development, one can significantly improve binding of the indole derivatives to the receptors involved in abrogating intestinal inflammation. Several indole analogs were evaluated as specific PXR agonists and their corresponding off-target liabilities were evaluated. The safety profile of the most promising candidates was assessed in mice and human intestinal organoids in culture. Finally, the action of the PXR-specific lead compounds on intestinal inflammation in mice and human organoids was evaluated.

## Results

### Characterization of FKK compounds as hPXR and/or AhR agonists

Based on our novel findings that indole and IPA can synergistically activate hPXR and can produce functional effects in vivo *(Ranhotra, Flannigan et al., 2016, Venkatesh et al., 2014)*, we hypothesized that the development of a small molecule, representing the interactions of both indole an IPA, is an innovative and potentially promising strategy for generating therapeutics for targeting PXR for diseases such as IBD. To design such molecules, we used our platform technology called the hybrid structure based (HSB) method. The HSB method utilized the interactions of both IPA and indole in the ligand binding domain (LBD) of PXR and the resulting pharmacophore was screened, and the molecules ranked by their individual docking score. Two commercial molecules FKK999 and BAS00641451(BAS451) (Appendix Figure S1a), which had docking scores of 65.89 and 52.66, respectively, were chosen as the starting point for further optimization. Docking studies demonstrated that although both FKK999 and BAS451 orient in the binding pocket of PXR to maximize their interactions with the residues from LBD, FKK999 had a better interaction profile that resembled the interactions of indole and IPA in the same site (Appendix Figure S1b). Docking of BAS451 shows several shared interactions with those of FKK999 but does not include the key ring stacking interaction with Trp299 and electrostatic interactions that contribute to the binding efficacy since BAS451 has only two indole rings and the additional phenyl ring does not compensate for the lost interactions. In PXR cell based transactivation screens, FKK999 activated PXR. BAS451 did not. FKK999 was used for further design of improved bis-indole analogs (FKK 1-10), and all analogs were tested as PXR ligands using in silico approaches (including sites outside the LBD, courtesy AC and SV). As demonstrated by the interactions of FKK5 in the LBD (Appendix Figure S1c) the ligand has arene-H interactions with Ser247 and Met250, electrostatic interactions with Met250 and Cys301 and other favorable interactions with Gln285, His407, Cys284, Met246 and Leu411.

The complete syntheses of indole metabolite mimics FKK1–FKK10 are reported in detail in the Supplementary Methods and Results. Crystal structures were obtained for lead compounds FKK5 (Appendix Figure S1d) and FKK6 (Appendix Figure S1e). The ligand efficiency metrics (LEM) (Cavalluzzi, Mangiatordi et al., 2017) analysis (Abad-Zapatero & Blasi, 2011) for the FKK compounds predicted – FKK5 and FKK6, as the best candidates for further studies. The same LEM analysis also indicated FKK4 as an efficient agonist; however, this congener could show metabolic instability with its *N*-protecting group possibly generating toxic aldehydes by oxidative cleavage in vivo.

All compounds synthesized were tested for their potential to activate PXR and/or AhR via luciferase assays as previously described (Huang, Wang et al., 2007),(Goodwin, Hodgson et al., 1999),(Novotna, Pavek et al., 2011). All FKK compounds demonstrate a concentration-dependent effect on PXR activation (Fig. 1a). By contrast, only FKK2 and FKK9 at 10 μM, respectively, demonstrated significant (>100 fold) AhR activation comparable with dioxin (TCDD) control ligand. To a much lesser extent, variable degrees of dose dependent AhR activation profiles were observed for FKK3,4,7,8,10 and FKK999 (Fig. 1b). FKK compounds did not activate glucocorticoid receptor (GR) (Appendix Figure S2a), vitamin D receptor (VDR) (Appendix Figure S2b), thyroid receptor (TR) (Appendix Figure S2c) and androgen receptor (AR) (Appendix Figure S2d) using cell-based luciferase assays previously described (Bartonkova, Grycova et al., 2016, Bartonkova, Novotna et al., 2015). Similarly, FKK5 or 6 did not induce constitutive androstane receptor (CAR) activity (Appendix Figure S2e). Since RXR is an obligatory protein partner in the active PXR transcription complex, we also assessed the potential for FKK compounds to serve as ligands for RXR (Xie, Barwick et al., 2000). Employing a recently developed radio-ligand binding assay for RXR (Toporova, Macejova et al., 2016), using nuclear extracts from rat liver, we only observed significant displacement of [^3^H]-9-*cis*-retinoic acid by FKK1, and to a lesser extent by FKK8 (Appendix Figure S2f). Since FXR and PPARγ are nuclear receptors known to abrogate colitis(Ning, Lou et al., 2019), we employed a TR-FRET based ligand displacement assay (FXR) (Appendix Figure S2g) and a cell line reporter assay (PPARγ) (Appendix Figure S2h), and did not observe any significant ligand agonist activity for both FKK5 and FKK6. While FKK6 activates human PXR, it does not significantly activate mouse PXR (Appendix Figure S2i). Together, molecular docking, LEM analysis, PXR and AhR transactivation assays show that among the FKK compounds, FKK5 and FKK6 are the best chemical PXR specific ligand leads.

**Fig. 1:**
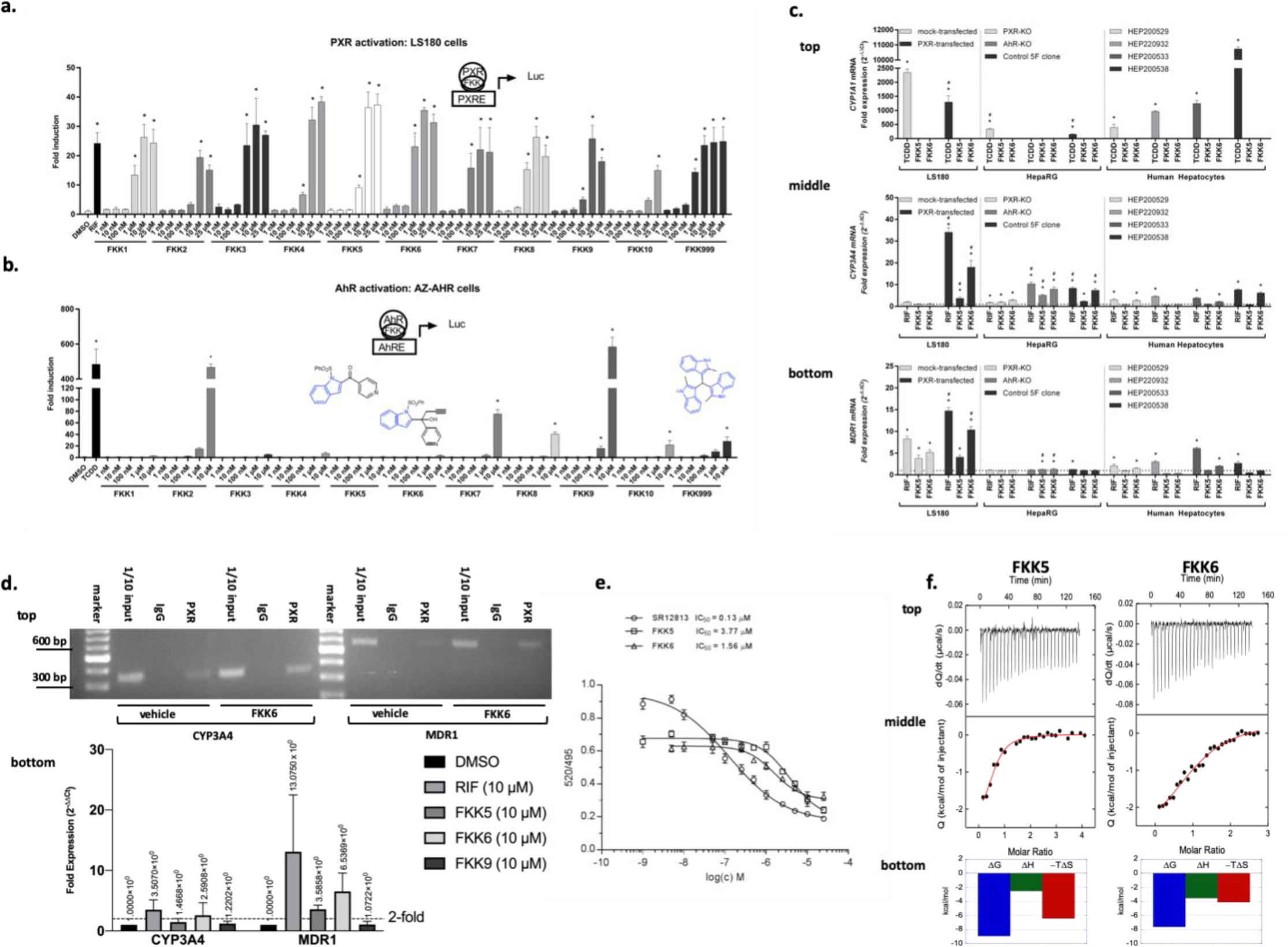
FKK5 and FKK6 activate PXR. **a**, Histogram (mean, 95 %CI) of fold induction of PXR activity reporter assay (luc, luciferase) in LS180 cells transiently transfected with wild-type PXR and p3A4-luc reporter plasmids. **b**, same as **a**, in HepG2 cells stably expressing pCYP1A1 luciferase plasmid (AhR reporter). Chemical structures of FKK5, FKK6 and FKK999 compounds are overlaid. Indole is colored blue. **a, b**, The bar graph(s) depicts one representative experiment of a series of experiments (*n* > 3) performed in four consecutive passages of cells. \**p* < 0.05, one-way ANOVA with Dunnett’s post hoc test. *significant over vehicle (DMSO) control. Histogram (mean, 95 %CI) of fold mRNA expression, of **c**, CYP1A1 (top panel), CYP3A4 (middle panel) and MDR1 (bottom panel) in LS180 cells with or without (mock) transfected PXR plasmid, HepaRG hepatic progenitor cells (PXR-knockout, PXR-KO; AhR-knockout, AhR-KO; parental control 5F clone) and primary human hepatocytes (HEP) from four donors is shown. The bar graph represents one experiment of a series of experiments (*n* > 3) performed in four consecutive passages of LS180 cells; *n* = 2 independent experiments with one well/compound and RT-PCR performed in triplicate for each HepaRG genotype; for each donor hepatocyte, *n* = 1 well/compound and RT-PCR performed in triplicate. ^*, #^*p* < 0.05, two-way ANOVA with Tukey’s post hoc test. *significant over vehicle control. ^#^significant over the same treatment in corresponding mock transfected or knock-out cells. **d**, Chromatin Immunoprecipitation (ChIP) assay in LS174T cells. Top panel, PCR products cells exposed to vehicle or FKK6 and run on a 2% agarose gel. DNA 100 base pair marker; Vehicle, 10% DMSO; FKK6 (10 mM); 1/10 Input – 0.2 million cells before IP; IgG – IP with polyclonal rabbit IgG; PXR – IP with PXR antibody. Bottom panel, quantitative PCR from the ChIP assay for compounds tested with the gene specific PCR amplicon normalized to GAPDH (fold expression). Dash line, 2-fold expression. The data is one representative experiment of two independent experiments (each *n* = 3 biological replicates, *n* = 4 technical replicates). Mean fold expression is expressed above each histogram (mean ± SD). **e**, PXR TR-FRET assay. TR-FRET ratio 520/495 nm are plotted against concentration of compound(s). Half-maximal inhibitory concentrations IC_50_ were obtained from interpolated standard curves (sigmoidal, 4PL, variable slope); error bars show standard deviation of *n* = 2 independent experiments each with four technical replicates. **f**, Microcalorimetric titrations of PXR ligand binding domain (LBD) with FKK5 and FKK6. A single-site binding model was used for fitting the data. The upper panel shows the output signal, dQ/dt, as a function of time. The middle panel shows the integrated heats as a function of the ligand/PXR molar ratio in the cell. The solid line represents the best non-linear least-squares fit of the data. The lower panel shows respective thermodynamic signatures of binding to the PXR ligand binding domain.

### Characterization of FKK Compound(s) Gene Expression Assay Profile in Cells

PXR agonists transcriptionally induce (> 2-fold) canonical target genes encoding drug metabolism enzymes/transporter, *CYP3A4* and *MDR1*, in both liver (hepatocytes) (Kandel, Thomas et al., 2016) and intestinal cells (LS180) (Gupta, Mugundu et al., 2008). HepaRG cells simulate hepatocytes in that PXR ligands can also induce target genes in similar but not identical manner (Andersson, 2010, Aninat, Piton et al., 2006, Antherieu, Chesne et al., 2012). AhR agonists transcriptionally induce target genes, *CYP1A1* and *CYP1A2*, in both hepatocytes (Pastorkova, Vrzalova et al., 2017) and intestinal cells (LS180) (Kubesova, Doricakova et al., 2016). As shown in Fig. 1c (top panel), in LS180 cells, FKK5 and FKK6, respectively, did not induce CYP1A1 mRNA. In PXR-transfected LS180 cells, FKK6 (and to a lesser extent FKK5), induce CYP3A4 (Fig. 1c middle panel) and MDR1 mRNA (Fig. 1c, bottom panel). In HepaRG cells harboring loss of PXR or AhR, target gene induction should be markedly diminished when compared to the wild type control cell line (Brauze, Zawierucha et al., 2017, Williamson, Lorbeer et al., 2016). TCDD, a known AhR ligand, induces CYP1A1 mRNA in HepaRG^TM^ control 5F clone and PXR-KO cells. FKK5 and FKK6 did not induce CYP1A1 mRNA (Fig. 1c, top panel). Rifampicin, a canonical PXR ligand, did not induce CYP3A4 mRNA in HepaRG^TM^ PXR-KO cells as compared with HepaRG^TM^ control 5F clone or HepaRG^TM^ AhR-KO cells. FKK6 > FKK5, respectively, induce CYP3A4 mRNA only in HepaRG^TM^ control 5F clone or HepaRG^TM^ AhR-KO cells (Fig. 1c, middle panel). By contrast, and unlike in LS180 cells, none of the compounds, including rifampicin, induced MDR1 mRNA (Fig. 1c, bottom panel). In primary human hepatocytes (plated, *n* = 4 independent samples), TCDD induced CYP1A1 mRNA in all hepatocytes. FKK5 and FKK6 did not induce CYP1A1 mRNA (Fig. 1c, top panel). Rifampicin and FKK6, unlike FKK5, showed >2fold induction in one of four hepatocyte samples (Fig. 1c, middle panel). Similarly, FKK6, but not FKK5, showed MDR1 mRNA induction in one of four hepatocyte samples (Fig. 1c, bottom panel).

In comparing the PXR target gene expression profile between cell-based models (Fig. 1c), there is a greater induction of CYP3A4 and MDR1 mRNA in LS180 cells when compared with HepaRG cells or primary human hepatocytes.

### Characterization of FKK Compound(s) Cell and tissue Cytotoxicity Potential

LS180 were used to assess cell cytotoxicity using the MTT assay as previously described (Bartonkova et al., 2016, Bartonkova et al., 2015). The results show that the entire series of FKK compounds (up to 10 μM) do not significantly alter LS180 cell survival (Appendix Figure S3a). Since in vitro cytotoxicity studies do not reflect in vivo effects of metabolites as would be generated from FKK compounds in vivo (animals), an acute toxicity study was conducted in C57BL/6 mice. FKK6 was selected as a lead compound for assessment, given that it had the more favorable PXR-selective ligand activity in cell lines tested. A dose of 500 μM in 10% DMSO (Castro, Hogan et al., 1995, Caujolle, Caujolle et al., 1967b) per day was chosen since this concentration represented the maximal solubility of FKK6 in an aqueous solution that could be safely administered to mice. The mice were gavaged for ten (10) consecutive days and necropsied on day 12 with collection of blood (serum). FKK6 does not alter serum chemistry profiles assessed (Appendix Figure S3b). FKK6 does not significantly impact tissue pathology when compared to control (vehicle) exposed mice (Table EV3). Similarly, a 30-day toxicity test was conducted in mice using FKK6 at an oral dose of 200 μM in 0.8% DMSO per day. This dose was chosen to stay within the safe DMSO dose administered to mice over a 30-day period. On day 30, mice were necropsied with collection of blood (serum). FKK6 does not significantly alter serum chemistry profiles (Appendix Figure S3c and d) or impact tissue pathology when compared to control (vehicle) exposed mice (Table EV4).

### Characterization of FKK-induced PXR – DNA interactions via Chromatin Immunoprecipitation (ChIP)

To determine if FKK5 and 6 are able to enhance occupancy of specific PXR DNA-binding elements (PXRE) in cells, we performed ChIP assays using LS174T cells transiently transfected with a PXR expressing plasmid. Extensively passaged LS174T cells endogenously express PXR but at low levels insufficient for good PXR protein pull-down and reporter experiments (Delfosse, Dendele et al., 2015). Thus, PXR transfection allows us to confidently isolate PXR occupancy effects in response to ligands. We verified a semi-quantitative ChIP using PXR-transfected LS174T cells exposed to FKK6 or its vehicle (DMSO). FKK6 augments PXR occupancy of the proximal CYP3A4 and MDR1 promoter (at their respective, PXR-binding elements) (Fig. 1d).

### Characterization of FKK 5 and 6 as direct ligands of PXR in solution

PXR activation is repressed via kinase-dependent phosphorylation (e.g., cdk2) (Smutny, Mani et al., 2013). FKK5 and FKK6 (DiscoverX *scan*MAX^SM^ assay panel of 468 kinases) were tested as potential inhibitors of kinases in vitro and neither of these compounds inhibits the kinases tested (Appendix Figure S4a & Table EV5). Together, these results support a direct interaction of FKK compounds with PXR. To demonstrate this interaction, we performed a cell-free competitive hPXR TR-FRET binding assay. In this assay, FKK5 demonstrated an IC_50_ of 3.77 μM which was similar to that observed with FKK6 ∼ 1.56 μM (Fig. 1e). To obtain direct proof of interaction with hPXR LBD, we performed isothermal titration calorimetry (ITC) experiments using His-tagged hPXR LBD protein (PXR.1) in solution. ITC measures the affinity, *K_a_*, and Gibbs energy (*ΔG = −RTlnK_a_*) and the changes in enthalpy, *ΔH*, and entropy, *ΔS*, associated with the binding of the FKK compounds (*ΔG* = *−RTlnK_a_* = *ΔH − TΔS*). Enthalpic and entropic contributions to binding affinity define the nature of the forces that drive the binding reaction (Ruben, Kiso et al., 2006, Velazquez-Campoy, Todd et al., 2000). This information is conveyed in a thermodynamic signature (Freire, 2008, Velazquez-Campoy, Kiso et al., 2001) and guides drug design and optimization (Garbett & Chaires, 2012). In agreement with previously demonstrated work that IPA alone has weak hPXR LBD interactions as determined using cell based assays (Venkatesh et al., 2014), in this assay, unlike rifampicin, a canonical hPXR agonist, IPA does not induce a thermal signature with respect to its interaction with the hPXR LBD in solution (Appendix Figure S4b). FKK5 binds to a single site in LBD, albeit with better affinity than IPA (*K*_d_ = 0.30 µM) due to a more favorable enthalpy of binding (*Δ*H = −2.5 kcal/mol, −T*Δ*S = −6.4 kcal/mol) (Fig. 1f). Compared to FKK5, FKK6 binds with improved enthalpy (ΔH = -−3.5 kcal/mol) but with an even larger loss in favorable entropy (−T*Δ*S = −4.1kcal/mol), which translates to a loss in binding affinity for FKK6 (*K*_d_ = 2.7 µM) (Fig. 1f).

In addition, we have previously shown that unlike rifampicin, indole in combination with IPA, is unable to activate the PXR mutant (C285I/C301A) (Venkatesh et al., 2014). Similarly, to test the effect FKK6 (10 μM) on this mutant, 293T cells were transfected with wild-type PXR.1 or mutant PXR.1 (C285I/C301A) plasmids along with its cognate luciferase reporter. The cells were exposed to DMSO, rifampicin (10 μM) or FKK6 (10 μM) and reporter activity assessed. While rifampicin significantly induces PXR reporter activity in both wild-type and mutant PXR transfected cells, FKK6 is unable to activate mutant PXR (Fig. 2a). Together, these data show that FKK5 and FKK6 are *bonafide* agonist ligands of PXR.1 in solution and cells.

**Fig. 2:**
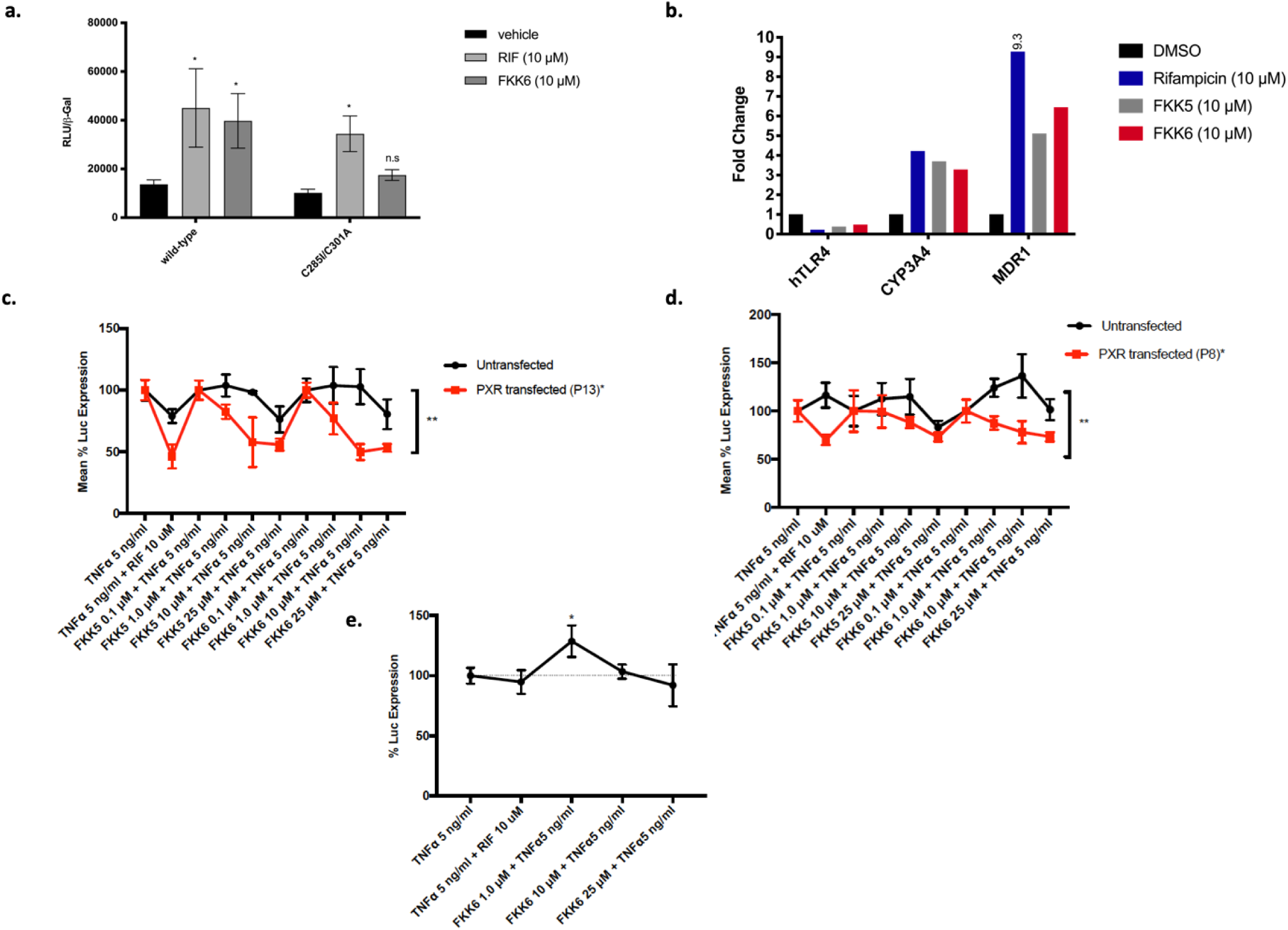
FKK5 and FKK6 inhibit NF-κB activation in a PXR-dependent manner. **a**, PXR (luciferase) reporter assay in HEK293T cells transiently transfected with PXR plasmids (wild-type and ligand binding domain mutant C285I/C301A). RLU, relative light units are shown normalized to beta-galactosidase (β-Gal) expression. The histogram represents one experimental data (of *n* > 2 independent experiments each performed in quadruplicate) mean (95 %CI); n.s. not significant; * *p* < 0.05, two-way ANOVA. **b**, Histogram represents fold change in mRNA expression, normalized to GAPDH, by RT-qPCR from Caco-2 cells exposed to compounds. DMSO, 0.1% DMSO vehicle; 9.3 is 9.3-fold expression. The data is one representative experiment of two independent experiments (each *n* = 3 biologic replicates, *n* = 4 technical replicates). Points and error represent mean (95 %CI) values for control (TNFα 5 ng/ml) normalized NF-κB (luciferase) reporter activity in LS180 cells after **c**, Passage 13 (P13) and **d**, Passage 8 (P8) and in **e**, LS174T PXR-KO (knockout) cells transiently transfected with pNL3.2.NF-κB-RE vector. **c**, **d**, Line graph depicts one representative experiment of a series of experiments (*n* > 4) performed in four consecutive passages of cells. Data was expressed as mean with 95% CI of four technical replicates. **e**, mean (± SD). *, ** *p* < 0.05, two way ANOVA with Tukey’s multiple comparisons test.

### Characterization of FKK5 and FKK6 on PXR-TLR4 and NF-κB signaling in human colon cancer cells

We have previously shown that PXR activation down-regulates TLR4 mRNA expression leading to inhibition of NF-κB signaling (Mani, 2016a, Ranhotra et al., 2016, Venkatesh et al., 2014). We have also shown that FKK5 and FKK6 interact directly with PXR LBD and result in its activation. Thus, to determine the effect of FKK5 and FKK6 on the PXR-TLR4-NF-κB signaling pathway, we used PXR-transfected Caco-2 cells and exposed these cells to FKK5 and 6 since undifferentiated Caco-2 cells inherently have low PXR expression (not detected) but relatively higher TLR4 expression and are excellent in vitro models for the study of enterocyte function(Delie & Rubas, 1997, Hung & Suzuki, 2018, Mani, 2016a, Ranhotra et al., 2016). FKK5 and FKK6 down-regulates TLR4 (0.39-fold and 0.49-fold, respectively) but up-regulates CYP3A4 (3.7-fold and 3.29-fold, respectively) and MDR1 (5.12-fold and 6.45-fold, respectively) mRNA expression (Fig. 2b). Together, these studies provide a rationale for the study of FKK compounds on modulating the PXR-TLR4-NF-κB pathway.

To determine whether FKK5 and 6 also inhibit NF-κB signaling in intestinal cells, we used PXR and NF-κB reporter co-transfected as well as NF-κB reporter transfected LS180 intestinal cell line in early and late passage. In this assay, the enhanced expression of PXR protein was verified by immunoblotting across several repeat experiments (Appendix Figure S4c). FKK5 and FKK6, in a dose-dependent manner, significantly reduced TNFα-induced NF-κB reporter activity in PXR-un-transfected and transfected LS180 cells (Fig. 2c). The effects of FKK compounds on inhibition of TNFα-induced NF-κB reporter activity are significantly greater in PXR-transfected cells as compared with un-transfected LS180 cells (Fig. 2c). Notably, the trend for effects of FKK5 and FKK6 is similar across the passages of LS180 cell lines used (Fig. 2c & d). Notably, PXR-transfected LS180 cells induce PXR target genes (CYP3A4, MDR1) well above un-transfected cells, and FKK5 and FKK6 induce PXR target gene expression in the transfected LS180 cells (Appendix Figure S4d).

To reproduce these findings in a more robust model, we used LS174T cells, in which PXR protein is absent, via a CRISPR-Cas9 knockout of the human PXR locus (see methods). Since the LS174T PXR knockout cells (*PXR*-KO or *NR1I2* KO) were pooled transfectants, the editing efficiency being ∼ 83%, we see that these cells can gain some functional PXR activity with multiple passages (Appendix Figure S5a) even though the PXR protein expression remains very low to undetectable (Appendix Figure S5b). FKK 5 and FKK6 do not induce PXR target genes in *PXR*-KO cells (Appendix Figure S5c). In early passage *PXR*-KO cells, we used PXR and NF-κB reporter co-transfected as well as NF-κB reporter-transfected *PXR*-KO cell line. However, we were unable to see sustained PXR protein in the PXR transiently transfected *PXR*-KO cells; thus, rather than perform the complementation experiment, we proceeded with establishing the effect of rifampicin and FKK6 in the TNF-exposed *PXR*-KO cell line transfected with the NF-κB reporter. Unlike the wild-type LS174T cells, neither rifampicin nor FKK6 repressed NF-κB reporter activity (Fig. 2e). Together, these data confirm that FKK5 and FKK6 are agonist ligands of PXR and that their activity in varied intestinal cells on repressing the TLR4 – TNF-α – NF-κB is directly via PXR.

### Characterization of FKK5 and FKK6 in human colon cancer cells and intestinal organoids exposed to inflammatory and infectious stimuli

To further characterize the anti-inflammatory effects of FKK5 and 6 on human intestinal cells and tissue, we first determined their effects on Caco-2 cell monolayers. These cells were well-established models to study intestinal cell differentiation and physiology, specifically under conditions of inflammatory stimuli (Hung & Suzuki, 2018). Differentiated Caco-2 cells via long term culture have heightened PXR expression (Mani, 2016b). In Caco-2 cells, interleukin-8 (IL8) rather than IL6 is consistently a more robust pro-inflammatory cytokine. In these cells, TNF-α significantly induced IL8 mRNA; however, only FKK6 (25 μM) significantly reduced cytokine-induced IL8 (Fig. 3a). Interestingly, cytokines (when compared to untreated cells) also significantly increased salmonella invasion in Caco-2 cells (Fig. 3b). FKK5 and 6, at all concentrations tested (10 and 25 μM), when compared to cytokines only, significantly decreased cytokine-induced salmonella invasion (Fig. 3b). There is no significant effect of FKK5 or 6 alone on salmonella invasion (Fig. 3b). Similarly, in Caco-2 cells, NF-κB is a seminal regulator of both IL8 and other pro-inflammatory cytokines. In Caco-2 cells, at 2 or 12 h, cytokines (CK only) translocate most of the NF-κB signal by immunofluorescence from the cytoplasm to the nucleus (when compared to No CK) (Fig. 3c). In the presence of FKK5 or 6 (25 μM, respectively), however, there is increased cytoplasmic NF-κB signal, suggesting more retention of NF-κB outside the nucleus (Fig. 3c).

**Fig. 3:**
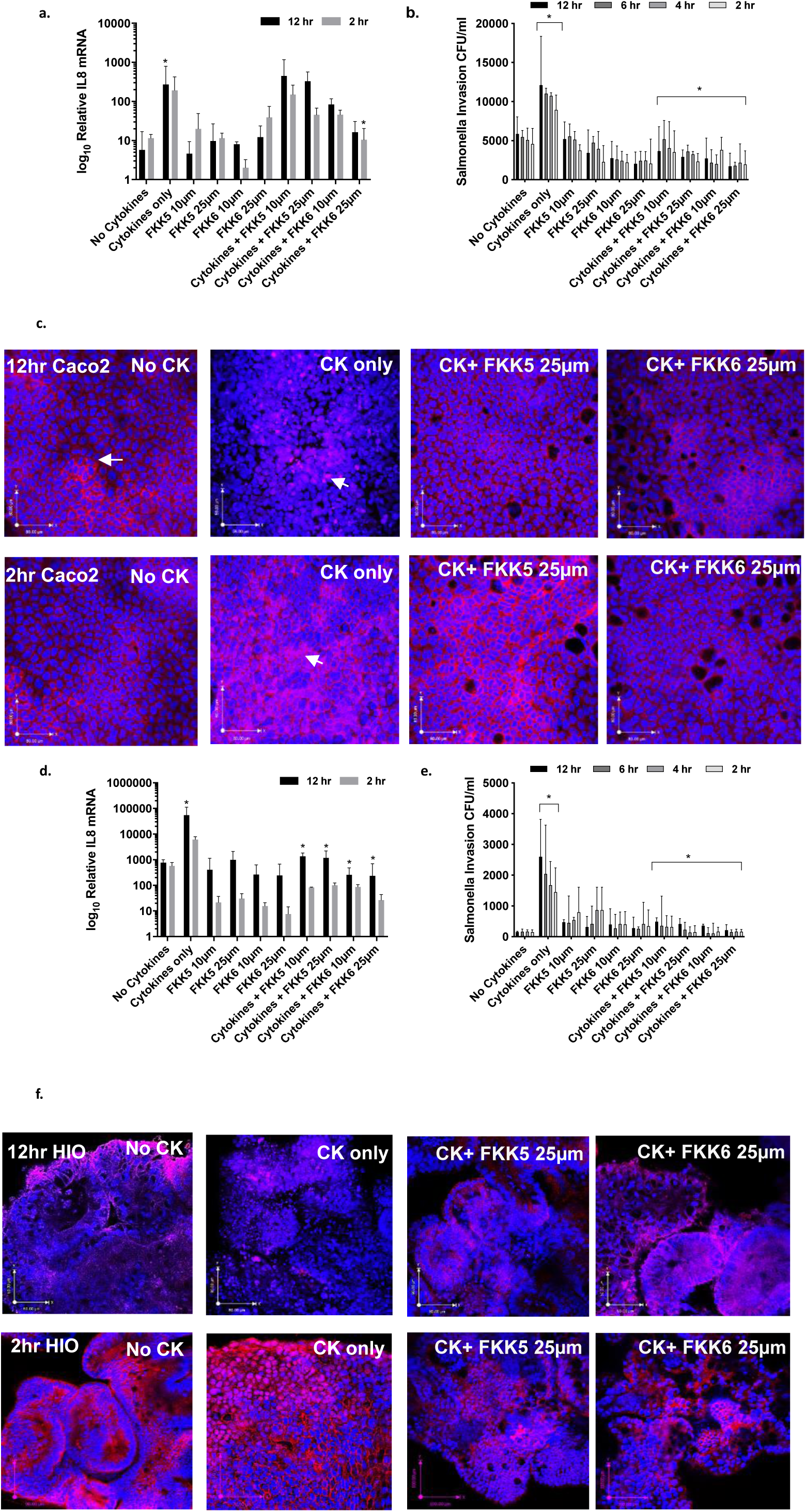
FKK5 and FKK6 inhibit cytokine and salmonella infection induced NF-κB nuclear translocation. In Caco-2 monolayer cells **a**, Log_10_ relative expression of IL8 mRNA with or without cytokines (50 ng/ml IL-1, 10 ng/ml IFNγ, 10 ng/ml TNF α), and cytokine cocktail plus FKK drugs (10 and 25 μM) with 2 h or 12 h incubations. **b**, Salmonella invasion in colony forming units per ml (CFU/ml) at the different time periods of incubation. **c**, confocal images of NFĸB (red) and the nucleus (blue). In human intestinal organoids (HIO), **d-f** are identical to **a-b**. The histogram, mean with 95% CI, depicts combined data from biological replicates (n = 3) with three consecutive passages of cells. Each biological replicate has three technical replicates. Results were expressed as. * *p* < 0.05, two-way ANOVA with Tukey post hoc test. * for cytokines versus no cytokines, compares either the 12 hr, 6hr, 4hr or 2 hr equivalent time-point(s) as indicated; * for cytokines versus cytokines + FKK compound, compares either the 12hr, 6hr, 4hr or 2hr equivalent time-point(s) as indicated. **c**, Arrows represent localization of NFĸB (red, cytoplasmic; purple, nuclear). Confocal images shown are representative images from (n = 3) biological replicates.

While Caco-2 cultures are important models to study intestinal barrier function, they have limitations in that culture conditions can widely change the experimental results (Delie & Rubas, 1997). Thus, for a more accurate representation of the effects of compounds on PXR activity, especially with regards to inflammation, we investigated the effects of our FKK analogs on primary human intestinal organoids (HIOs). In iPSC derived HIOs, cytokines induce a significant increase in IL8 mRNA levels at 12 h (Figure 3d; *p* < 0.05 Two Way ANOVA) but not at 2 h after exposure. FKK5 and FKK6 (each at 10 and 25μM, respectively), has no significant effect on basal IL8 expression (Fig. 3d); however, when compared to cytokines alone, in the presence of cytokines both FKK5 and 6 significantly reduced IL8 mRNA expression after 12 h of exposure (Fig. 3d). Similarly, cytokines significantly increased salmonella invasion in HIOs (Fig. 3e). FKK5 and FKK6 (both at 10 and 25 μM, respectively), had no significant effect on basal levels of salmonella invasion (Fig. 3e); however, when compared with cytokines alone, FKK5 and FKK6, both significantly reduced salmonella invasion (Fig. 3e). Interestingly, salmonella invasion is significantly reduced at all time-points studied (2, 4, 6 and 12 h post exposure to cytokines). In HIOs, at 2 or 12 h, by visual inspection of the immunofluorescence images, cytokines (CK only) translocated most of the NF-κB signal from the cytoplasm to the nucleus (CK+FKK compared to No CK) (Fig. 3f). In the presence of FKK 5 or 6 (25 μM, respectively), however, there was increased cytoplasmic NF-κB signal, suggesting more retention of NF-κB outside the nucleus (Fig. 3f).

In another HIO system, we confirmed that our analogs, FKK5 and 6, significantly induce PXR target genes (Appendix Figure S6a). Interestingly, CYP3A4, a target gene that is only modestly (as opposed to MDR1) elevated in our intestinal cell lines exposed to FKK5 (Fig. 1c), was also only modestly increased in the HIO experiments (Supplemental Fig. 6a: borderline significance for CYP3A4 *p* = 0.056). Since HIOs can be used to study effects of FKK compounds on PXR target genes, to determine if this would correlate with their ability to inhibit TNFα mediated induction of pro-inflammatory cytokine (IL8), HIOs from different individuals were exposed to TNFα in the presence or absence of FKK5 (*n* = 9) or FKK6 (*n* = 3) or FKK9 (*n* = 5). In pair-wise comparison of HIOs from the same individual exposed to DMSO or FKK compounds, respectively, FKK5 (10 μM) significantly lowered IL8 mRNA levels (Fig. 4a); similarly, FKK6 (10 μM) reduced IL8 mRNA levels in organoids but did not reach statistical significance (*p* = 0.07) (Fig. 4b). FKK9, which is a strong AhR agonist but also has PXR agonist activity, did not show significant effects on IL-8 mRNA levels in this assay (Fig. 4c).

**Fig. 4:**
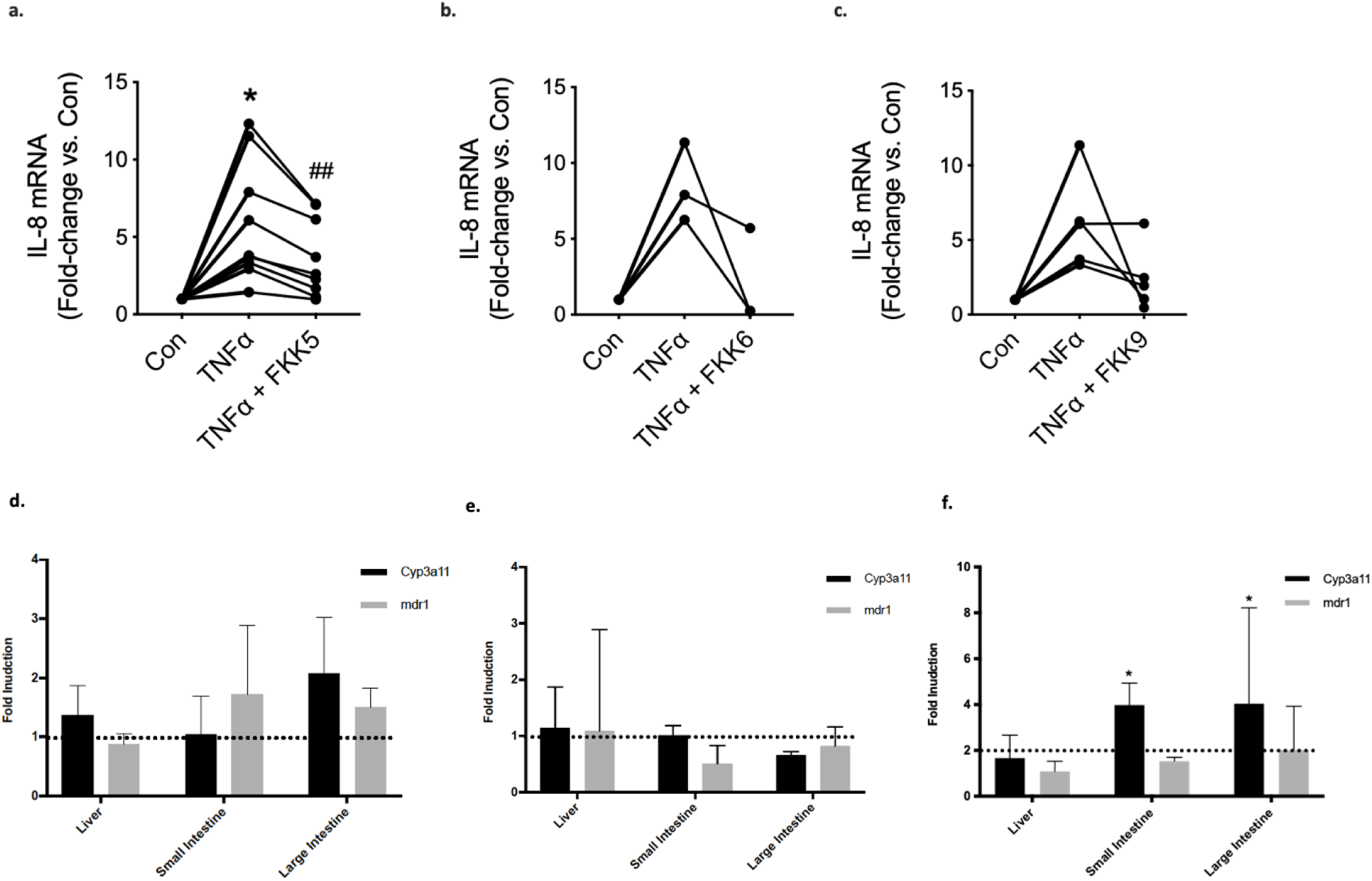
FKK6 inhibits cytokine mediated IL8 induction in human colonic organoids and induces PXR target genes in mice. Quantitative gene expression by RT-qPCR of IL8 mRNA from human colonic enteroids (over 40 enteroids per individual sample per well) exposed to control (vehicle) or TNF α. with or without **a**, FKK5 (*n* = 9 paired samples). **b**, FKK6 (*n* = 3 paired samples; one removed due to poor inflammatory response). **c**, FKK9 (*n* = 5; 3 removed due to poor response and tissue integrity). Data shown as mean fold change relative to vehicle control (con) for each paired sample (*n* = 3 replicates). Each paired sample data set represents biopsies from an individual patient. ^*, ##^ *p* < 0.05, Pair-wise one way ANOVA with Tukey’s post hoc test. Fold induction (mRNA) of genes in **d**, C57BL/6 mice, **e**, *pxr^-/-^* mice, and **f**, *hPXR* mice (mice expressing the human PXR gene) gavaged with vehicle (10% DMSO; *n* = 3) or FKK6 (500 μM in 10% DMSO; *n* = 3) every 12h for 3 total doses. The entire experiment was repeated two independent times and one representative experiment is shown. Each mouse (each organ) was studied in quadruplicate assays and normalized to internal control, GAPDH. The histograms show mean (95% CI) values for gene expression. * *p* < 0.05, two-way ANOVA with Tukey’s post hoc test.

Together, these data in Caco-2 cells, HIOs derived from multiple sources and centers, suggest that the lead FKK compounds, FKK5 and 6, are capable of inducing PXR target gene expression in all intestinal cell types. This correlates with reduction of TNFα induced pro-inflammatory cytokine (IL8) expression, reduced invasion from intestinal pathobiont, *S. typhimurium* as well as reduced accumulation of nuclear NF-κB.

### PXR target gene expression in mice

The FKK compounds presented here are first generation indole/IPA analogs that have not been optimized for pharmaceutical discovery. However, to determine the effects in vivo, specifically as it relates to the ability to induce PXR target genes in different mice tissues after oral gavage, we performed studies using FKK6, our representative best lead, in C57BL/6 mice. FKK6 was gavaged at concentration ∼ 500 μM in 10% DMSO and administered to C57BL/6 mice over 36 or 60 h (3 or 5 doses, respectively). There was no significant effect (< 2-fold induction with large variability) in PXR target gene (cyp3a11, mdr1a, mdr1b or mdr1) expression across organs (liver, small and large intestine) in mice (Fig. 4d: 36 h; Appendix Figure S6b: 60 h). There is a borderline induction of cyp3a11 (2.07-fold) in the large intestine of wild-type mice (Appendix Figure S6b). In *pxr^-/-^* mice exposed to FKK6, the expression levels for all target genes (cyp3a11, mdr1) are ≤ 1.5fold to that of DMSO exposed mice. Indeed, there is no biologically significant effect on PXR target gene (mdr1) expression across organs (liver, small and large intestine) in *pxr^-/-^* mice (Fig. 4e: 36 h; Appendix Figure S6c: 60 h). However, cyp3a11 is significantly suppressed in the small (0.25x DMSO) and large intestines (0.36x DMSO) in the 60 h exposure study (Appendix Figure S6c). These data suggest that PXR might be important in these organs as basal regulators of these genes. By contrast, there is a biologically significant induction of cyp3a11 (but not mdr1) expression in small and large intestines as compared to DMSO and/or liver cyp3a11 expression in *hPXR* mice (mice expressing the human PXR gene) when exposed to 36 h of FKK6 (three doses) (Fig. 4f). Interestingly, when the FKK6 dosing is prolonged (five doses or 60 h exposure), there is a distinct increase in mdr1 expression in the liver (as compared to the intestines) and no significant effects on cyp3a11 expression in *hPXR* mice (mice expressing the human PXR gene) (Appendix Figure S6d). Together, these results demonstrate that FKK compounds induce PXR target genes in a PXR-dependent manner in mice.

### Effect of FKK6 in mouse model of acute colitis in vitro and in vivo

To characterize the effects of lead compound, FKK6, on murine colitis, adult C57Bl/6 female *hPXR* mice (mice expressing the human PXR gene) were exposed to DSS acutely. Mice either received once daily vehicle gavage plus intrarectal bolus or FKK6 (200 μM) oral gavage plus intrarectal bolus (200 μM) for 10 days until necropsy. FKK6 treated mice showed significantly less weight loss and significantly greater colon length on day 10 (Fig. 5a, b). The inflammation score was significantly less for FKK6 treated mice (Fig. 5c); however, the fecal lipocalin 2 and FITC-dextran measurements show a decreased trend only (Fig. 5d, e). These findings are replicated when we use the clustering method (weight loss, colon length), wherein it correctly identifies the two groups of mice that received two different treatments. By including, fecal lipocalin 2 as a cluster feature in the analysis then the clustering method identifies the two groups of mice correctly except for one of the samples. The intestinal tissue cytokines, IL10 and TNFα (and TNFR2) showed a significant increase and decrease, respectively, in FKK6 treated mice as compared with controls (Fig. 5f). IL17f, a IL17a heterodimer partner, is a prime mediator of colitis (Tang, Kakuta et al., 2018), and this cytokine is significantly reduced in FKK6 treated mice as compared with controls (Fig 5f). By contrast, FKK6 did not alter weight loss, colon length, inflammation score, lipocalin 2, FITC-dextran levels in *pxr^-/-^* mice (Fig. 6a–f). Together, these results show that a representative FKK lead compound, FKK6, significantly reduces dextran sodium sulfate (DSS) induced colitis in mice in a PXR-dependent manner.

**Fig. 5:**
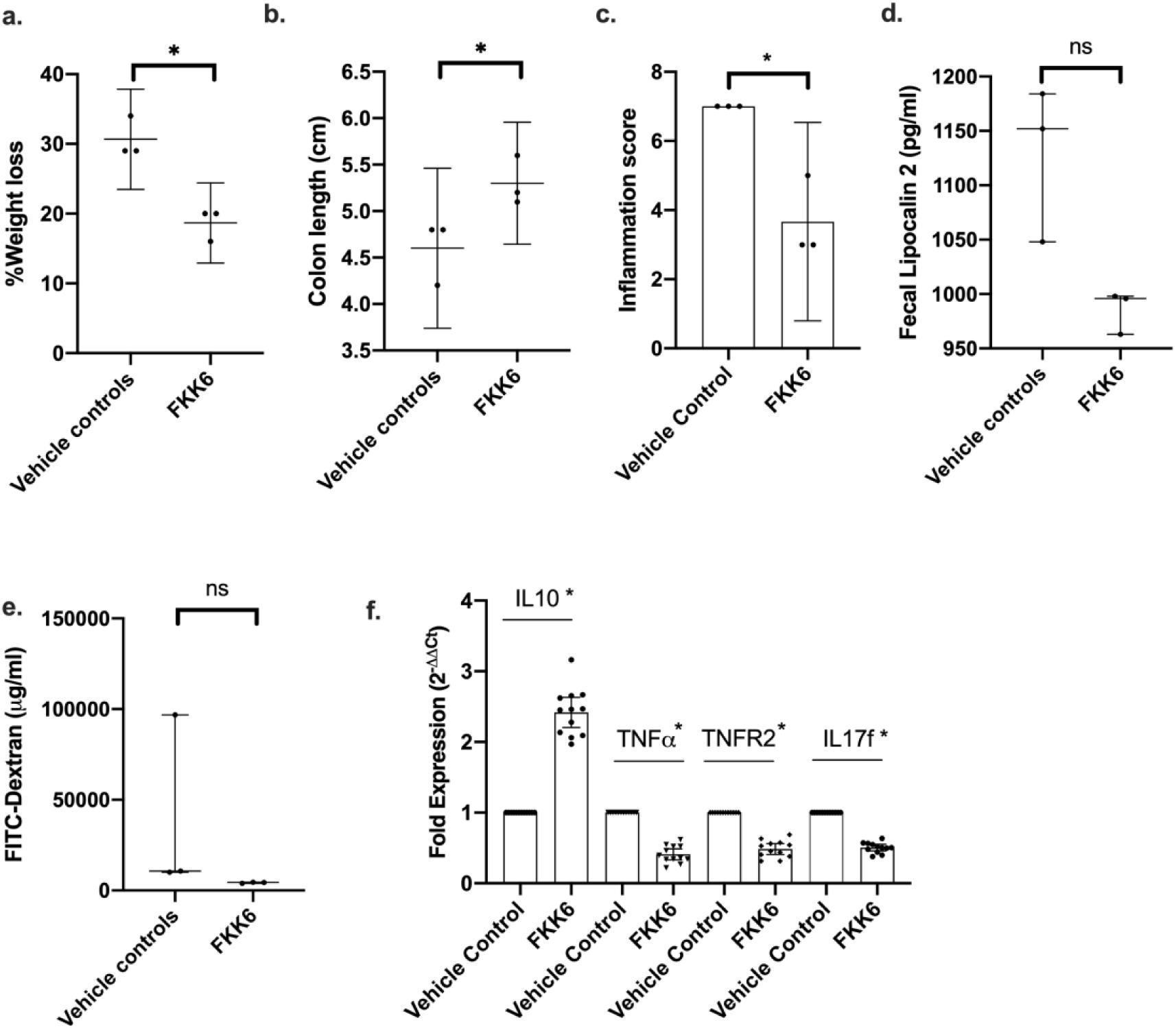
FKK6 abrogates Dextran Sulfate Sodium induced colitis in mice expressing the human PXR gene. After coin-toss randomization, *hPXR* mice (mice expressing the human PXR gene) were allocated to treatment with vehicle (0.8% DMSO) (n = 3/genotype) or FKK6 (200 micromolar) (n = 3/genotype), by simultaneous oral gavage and intra-rectal delivery starting day 1 – 10 of DSS administration. **a**, % weight loss from baseline (day 1 *vs* day 10) **b**, colon length (cm) **c**, Inflammation score (see methods) **d**, Fecal lipocalin 2 (pg/ml) **e**, serum FITC-dextran (μg/ml). **a – c**, mean (95 %CI). **d**,**e**, median (interquartile range). **f**, fold expression of mRNA as illustrated in *hPXR* mouse colon tissue exposed to DSS (n = 3/group; each PCR performed in quadruplicate). (**a-e**), The entire experiment was repeated three independent times and one representative experiment is shown. *(**a**, **b**, **c**) *p* < 0.05, Welch’s t-test; *(**d**, **e**) p < 0.05, Mann-Whitney test; ns, not significant; (**f**), *compares FKK6 versus vehicle control for each gene, *p* < 0.05, Two-way ANOVA

**Fig. 6:**
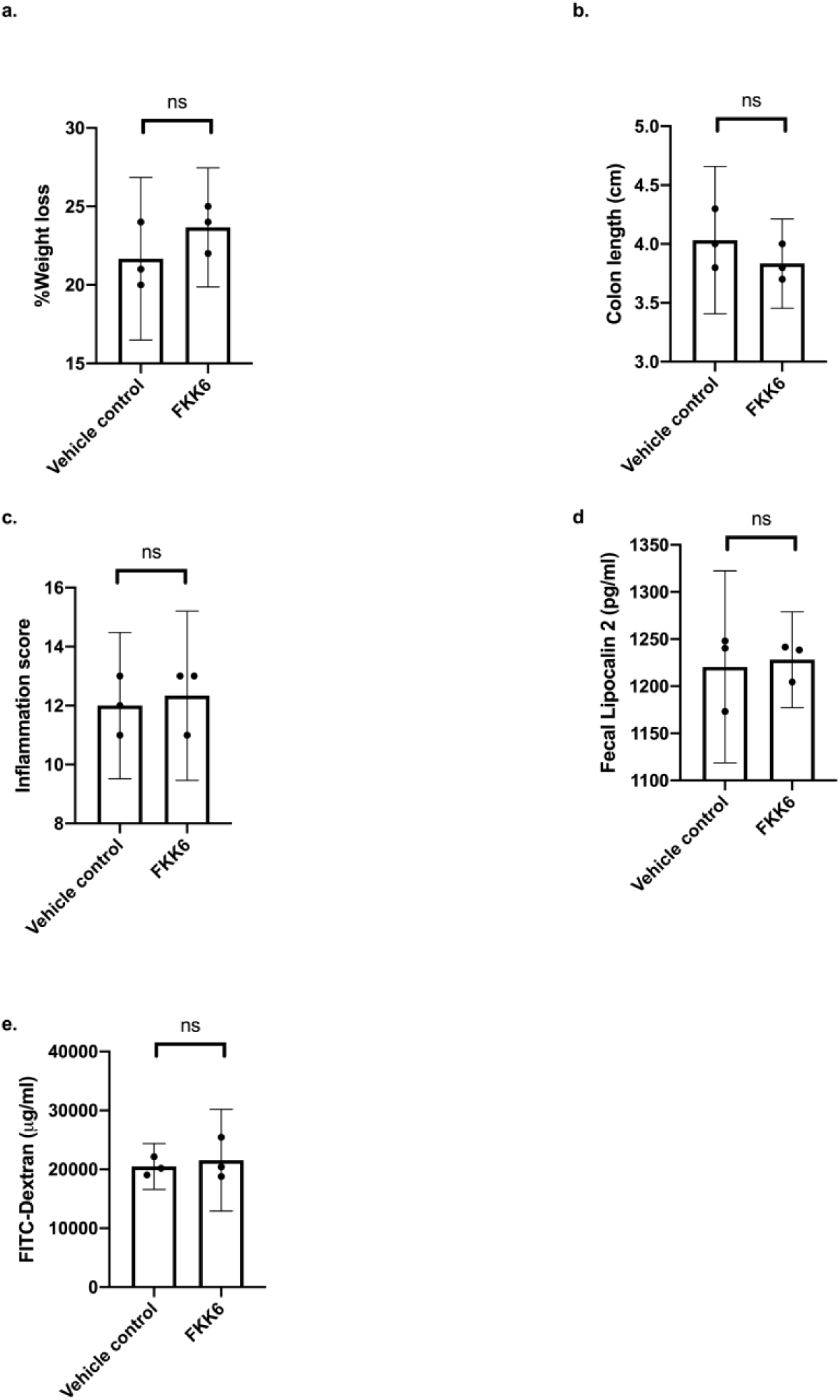
FKK6 does not abrogate Dextran Sulfate Sodium induced colitis in *pxr ^-/-^* mice. As in Fig 5, p*xr ^-/-^* mice underwent the identical experimental procedure and analysis. The entire experiment was repeated three independent times and one representative experiment is shown. *(**a – e**) mean (95 %CI); Welch’s t-test; ns, not significant.

## Discussion

A longstanding problem in drug design and discovery is the very limited regions of chemical space that are exploited by the molecules conventionally available to small and large pharmaceuticals and academia. While it has been proposed that natural metabolite derivatives are likely to serve as more potent and well tolerated drugs, proof of this concept is lacking (Saha et al., 2016). Herein, we show that indole compounds of the FKK series mimic the docking of the natural indole and indole propionic acid derivatives for PXR, and have produced two potent lead compounds, FKK5 and FKK6. While not optimized for therapeutic delivery, these leads showed significant activity in suppressing intestinal inflammation in vitro and, for FKK6, in mice and human tissues simulating colitis.

There are several caveats to bear in mind when evaluating the data presented. While we show direct binding of FKK6 to PXR protein, our studies have not resolved the direct binding residues that interact with FKK6. Ongoing research to obtain a crystal structure of co-crystallized PXR and FKK is currently underway. The in vivo disposition of FKK compounds and their metabolites is currently unknown. The importance of pharmacokinetic profiling is that it will allow us to partially explain the serendipitous results that our FKK compounds lack PXR activation properties in liver cells (primary hepatocytes and HepaRG), as compared with intestinal cells (LS180, LS174T). The metabolites generated in mice may also have importance, in that they could inhibit parent compound interactions with PXR, and these will be studied further. Recent research reveals that intestinal PXR activators can induce hypertriglyceridemia and metabolic perturbations (Meng, Gwag et al., 2019). We did not observe any significant changes in lipid profiles in mice exposed to either a 10-day or 30-day gavage of FKK6. Indeed, it is likely that one explanation for this might be that PXR activation can have opposing effects on lipid metabolism depending on the nature of the xenobiotic itself (i.e. non-receptor mediated action of the compound itself like anti-oxidant function) (Hwang, Yoo et al., 2009). These effects may cancel out on certain pathways like lipid metabolism (Hwang et al., 2009). FKK6 has been studied with respect to a limited number of cellular targets that relate to PXR. While we find that FKK effects in cells and human intestinal organoids (HIOs) correlates with changes in PXR target gene and NF-κB signaling, it is possible that some of its protective effects could come via other targets hitherto uncharacterized (e.g., tubulin) (La Regina, Bai et al., 2015). Loss of PXR studies in HIOs would be required to confirm these correlations; these studies are in progress using both LS174T cells and iPSC cells in which PXR is knocked out using CRISPR-Cas9. We have shown that FKK6 has utility in the DSS-induced model of colitis. The use of mouse models beyond DSS induced colitis (e.g., *IL10^-/-^*) could broaden the translatability of these compounds during preclinical drug development. Future strategies to improve solubility and targeted delivery could include formulations used in existing compounds for colon targeted delivery (Guo, Zong et al., 2018). Despite these caveats, our studies provide and validate the initial proof-of-concept of the utility of microbial metabolite mimicry as a potential drug discovery modality.

The prototypical pharmacologic ligands of human PXR include rifaximin, rifampicin, hyperforin and SR12813 (Orans, Teotico et al., 2005). Rifaximin is human PXR agonist ligand, but as an antibiotic, changes intestinal microbiota that has resulted in emergence of drug resistant with chronic administration (Farrell, 2013). Furthermore, despite the claim that rifaximin is non-absorbable via the intestinal tract, chronic administration of rifaximin to mice has resulted in hepatic steatosis (Cheng et al., 2012a). Indeed, while controversial(Abdel-Razik, Mousa et al., 2018), rifaximin can potentially induce insulin resistance in humans and is not beneficial in the treatment of non-alcoholic steatohepatitis (Cobbold, Atkinson et al., 2018). Finally, for the rifamycin family (rifaximin and rifampicin), it is possible that targets other than nuclear receptors, such as p300 or other enhancer activation, may be more salient in determining PXR and non-PXR target gene interactions (Smith, Eckalbar et al., 2014, Yan, Wang et al., 2017). Upon chronic dosing, rifampicin accumulation is not targeted towards the intestines but towards brain and lungs (Shan, 2004). Similarly, hyperforin has multiple cellular targets (e.g., reactive oxygen species, elastases, Protein kinase C) (Galeotti, Vivoli et al., 2010),(Feisst & Werz, 2004, McCue & Phang, 2008). Hyperforin pharmacokinetics suggest rapid absorption without accumulation in tissues and thus with each dose causing clinically relevant adverse drug-drug interactions (Biber, Fischer et al., 1998, Soleymani, Bahramsoltani et al., 2017). Hyperforin can induce reversible liver injury in mice (Negres, Scutari et al., 2016). SR12813 induces degradation of 3-hydroxy-3-methylglutaryl-coenzyme A (HMG-CoA) reductase independent of effects on PXR (Berkhout, Simon et al., 1996). It is to be noted that for unknown reasons, SR12813 was never clinically developed as an anti-cholesterol drug. Our PXR ligands, in particular FKK6, has the tendency to activate intestinal PXR and target genes much more robustly than in the liver. In acute and 30 day toxicology studies in mice, we did not find changes in lipid metabolism or associated pathology. In the kinase assay screen, there was very little effect of either FKK5 or FKK6 to inactivate kinases or interfere with their activation function (e.g., p300). While only a host of enzymes and receptors have been evaluated, it is of interest that these indole analogs have limited cross-receptor activation as opposed to their parent indole metabolites from bacteria (Venkatesh et al., 2014). Thus, with this rationale, in the future, we will perform detailed head-to-head comparisons of our compounds against known PXR ligands for bench marking efficacy and chronic toxicity.

In summary, the FKK lead compounds are suitable starting points for further optimization of potency and/or for improving drug-like qualities and similar properties for preclinical drug discovery. Possible clinical applications would be in fields in which increased intestinal permeability potentiates disease manifestations (e.g., inflammatory bowel disease) (Bischoff, Barbara et al., 2014). Additionally, since these are relatively well tolerated agents, applications in chemoprevention applications are also possible. Our work describes the first fully realized drug discovery using microbial metabolite mimicry. These strategies could be expanded widely to other receptors (and diseases) that derive ligands from the host microbiome.

## EXPERIMENTAL SECTION (Online Methods)

### Chemistry, Compound Crystal structure, LEM analysis and In silico experiments

See *Supplementary Methods*

### Biology

Rifampicin, 2,3,7,8 – tetrachlorodibenzo-*p*-dioxin (TCDD), dexamethasone, 1α,25 Dihydroxyvitamin D_3_, Triiodo-L-Thyronine (T3), Dihydrotestosterone (DHT), SR12813 (S4194) were purchased from Sigma. LS174T cells were originally purchased from ATCC and prior to use cell line authenticity and was validated as previously described (Wang, Venkatesh et al., 2011). Caco-2 cells were purchased from ATCC and was validated by multiplexed STR DNA profiling performed by Genetica. Human Caucasian colon adenocarcinoma cells LS180 (ECACC No. 87021202) were purchased from European Collection of Cell Cultures (ECACC). Stably transfected gene reporter cell lines AZ-AHR, AZ-GR, IZ-VDRE, PZ-TR and AIZ-AR were as described elsewhere (Bartonkova et al., 2016, Bartonkova et al., 2015, Illes, Brtko et al., 2015, Novotna et al., 2011, Novotna, Pavek et al., 2012). Cells were cultured in Dulbecco’s modified Eagle’s medium (DMEM) or RPMI-1640 medium (cultivation of AIZ-AR cells) supplemented with 10% of fetal bovine serum, 100 U/mL streptomycin, 100 mg/mL penicillin, 4 mM L-glutamine, 1% non-essential amino acids, and 1 mM sodium pyruvate. Cells were maintained at 37 °C and 5% CO2 in a humidified incubator. HepaRG^TM^ is a human hepatoma cell line isolated from a liver tumor of a female patient suffering from hepatocarcinoma and hepatitis C infection (Gripon, Rumin et al., 2002). The cells possess a pseudodiploid karyotype and have been characterized as an oval duct bi-potent hepatic cell line as they have the ability to differentiate into both biliary and hepatocyte lineages in the presence of DMSO(Parent, Marion et al., 2004). Three HepaRG^TM^ derived cell lines, including 5F Clone control cells, and cells with targeted functional PXR and AhR gene knockouts, PXR-KO and AhR-KO respectively, were purchased from Sigma (CZ). Cells were handled according to the manufactureŕs protocol. Human Hepatocytes were purchased from Biopredic International (France) and Triangle Research Labs, LLC-Lonza (US). Cultures were maintained in serum-free medium at 37°C and 5% CO_2_ in a humidified incubator.

### Generation of PXR knockout LS174T cell line

LS174T cells (ATCC: CL-188; 7 000 3535) (Synthego Corporation, Redwood City, CA) were the parental cells used to generate pooled clones of PXR knockout. CRISPR Cas9 mediated knockout cell (pool/clone) of (gene name) in (cell line) cells were generated by Synthego Corporation (Redwood City, CA, USA). Briefly, cells were first tested negative for mycoplasma. Guide RNA was selected for high activity, specificity and activity to create premature stop codons through frameshift mutations in the coding region via insertions and or deletions (Indels)within exon 2 of the gene encoding human *PXR* (*NR1I2* transcript ID ENST00000337940). To generate specific guide RNAs, based on off-target analysis, the following modified RNAs were selected – *NR1I2-119,807,249 5’ AAGAGGCCCAGAAGCAAACC-3’* [*TGG*]*-PAM*. To generate these cells, ribonucleoproteins (RNPs) containing the Cas9 protein and synthetic chemically modified sgRNA (Synthego) were electroporated into the cells using Synthego’s optimized protocol. Editing efficiency is assessed upon recovery, 48 h post electroporation. Genomic DNA is extracted from a portion of the cells, PCR amplified and sequenced using Sanger sequencing. The resulting chromatograms are processed using Synthego Inference of CRISPR edits software (ice.synthego.com). The pooled PXR KO clones were validated for PXR protein (loss of) expression and target gene induction activity through serial passages (*Appendix Figure S5a & b*).

### Cell Culture and Transfection Assays

Cells were seeded in 96 well plates, stabilized for 24 h, and then incubated for 24 h with tested compounds; vehicle was DMSO (0.1 % v/v). Incubations were performed in four technical replicates. LS180 cells transiently transfected by lipofection (FuGENE HD Transfection Reagent) with pSG5-PXR plasmid along with a luciferase reporter plasmid p3A4-luc(Goodwin et al., 1999, Huang et al., 2007), were used for assessment of PXR transcriptional activity. Transcriptional activity of AHR, GR, VDR, TR and AR were studied in stably transfected gene reporter cell lines AZ-AHR, AZ-GR, IZ-VDRE, PZ-TR and AIZ-AR, respectively (Bartonkova et al., 2016, Bartonkova et al., 2015, Illes et al., 2015, Novotna et al., 2011, Novotna et al., 2012). Cells were seeded in 96-well plates, stabilized for 24 h, and then incubated for 24 h with tested compound and/or vehicle DMSO (0.1% v/v) in the presence (antagonist mode) or absence (agonist mode) of rifampicin (RIF; 10 μM), 2,3,7,8-tetrachlorodibenzo-*p*-dioxin (TCDD; 5 nM), dexamethasone (DEX; 100 nM), calcitriol (1α,25-VD3; 50 nM), 3,5,3’-triiodothyronine (T3; 10 nM) or dihydrotestosterone (DHT; 100 nM) respectively. After the treatments, cells were lysed and luciferase activity was measured on Tecan Infinite M200 Pro plate reader (Schoeller Instruments, Prague, Czech Republic). The data are expressed as fold induction ± SD of luciferase activity over the control cells (agonistic mode — in the absence of a model ligand) or as a percentage of maximal activation ± SD (antagonistic mode — in the presence of a model ligand). Differences were tested using one-way ANOVA with Dunnett’s post hoc test, *p* < 0.05, was considered significant (*). For data presented in Supplementary Figure 8, transfections and analysis of AhR activation was performed as previously published (Murray, Flaveny et al., 2010). For all the transfection experiments, specifically mutiplexed transfections, the control wells were mock transfected (PXR vector). However, to exclude that PXR ligand (FKK) effect is not due to the non-specific binding of PXR (or other factors) upstream of the luciferase gene within non-PXR binding elements, we performed pilot experiments in different cell passages (P11, P14, P17) of LS174T cells using a mock luciferase plasmid (luciferase not containing an upstream PXR binding element of the CYP3A4 promoter). The PXR ligands tested were rifampicin (10 μM) and FKK6 (1−25 μM). There was no effect of these ligands at any concentration on luciferase reporter activity (data not shown).

Reverse transfection assays were performed in Caco-2 cells as previously published with the following modifications (Kublbeck, Anttila et al., 2015). The assays were performed in a 48-well plate format using 3.0 × 10_4_ cells per well using pSG5-PXR plasmid (300 ng/well); β-Gal expression plasmid (600 ng/well); CYP3A4-luciferase reporter (500 ng/well) as previously published (Huang et al., 2007). Transfection using lipofectamine_®_ LTX (1 μl/mg of plasmid) proceeded for 24 h followed by an additional 24 h exposure to FKK5. Cell lysates were prepared for β-Gal and luciferase assays as previously published (Huang et al., 2007). The data are expressed as mean ± SD RLU (relative light units from luciferase assay) normalized to β-Gal activity in the same well. Differences were tested using 2-way ANOVA with Tukey’s multiple comparison test, *p* < 0.05 was considered significant (*). Standard transient transfection assays (PXR, CAR) in HEK293T cells were performed as previously published (Huang et al., 2007).

NF-κB reporter assays were performed using LS180 cells. Briefly, LS180 cells (Passage 8 and 13 were used since NF-κB activity can be influenced by cell passage) (Mastropietro, Tiscornia et al., 2015) were transiently transfected using Fugene HD transfection reagent with a reporter plasmid pNL3.2-NF-κB-RE[NlucP/NF-κB-RE/hygro] from Promega (Hercules, CA, USA); with or without co-transfection of the wt-PXR expression vector. The transfected cells were seeded in 96-well plates at density 2.5×10_4_ cells per well. Following 16 h of stabilization, cells were treated with a vehicle (DMSO; 0.1% v/v), rifampicin (RIF; 10 µM) or tested compounds FKK5 and/or FKK6 in a concentration range from 0.1 to 25 µM for 24 h. For the last 4 h of the treatment, a combination of tested compounds or rifampicin with tumor necrosis factor α (TNFα; 5 ng/mL), a model inducer of NF-κB receptor transcriptional activity, was applied. Following the incubation time, cells were lysed, and Nano luciferase activity was measured in 96-well format using a Tecan Infinite M200 Pro plate reader (Schoeller Instruments, Czech Republic).

### Quantitative Real-Time PCR

Cells were seeded in 6-well plates, stabilized for 24 h, and then incubated for 24 h with test compounds; vehicle was DMSO (0.1% v/v). Primary human hepatocytes from four different donors (HEP200529, HEP220932, HEP200533, HEP200538) were incubated for 24 h with test compounds; vehicle was DMSO (0.1% v/v). Total RNA was isolated using TRI Reagent® (Molecular Research Center, Ohio, USA). cDNA was synthesized from 1000 ng of total RNA using M-MuLV Reverse Transcriptase (New England Biolabs, Ipswich, Massachusetts, USA) at 42 °C for 60 min in the presence of random hexamers (New England Biolabs). Quantitative reverse transcriptase-polymerase chain reaction (qRT-PCR) was performed using LightCycler® 480 Probes Master on a LightCycler® 480 II apparatus (Roche Diagnostic Corporation). The levels of *CYP1A1, CYP1A2*, *CYP3A4, MDR1* and *GAPDH* mRNAs were determined using Universal Probes Library (UPL; Roche Diagnostic Corporation) probes and primers listed below:

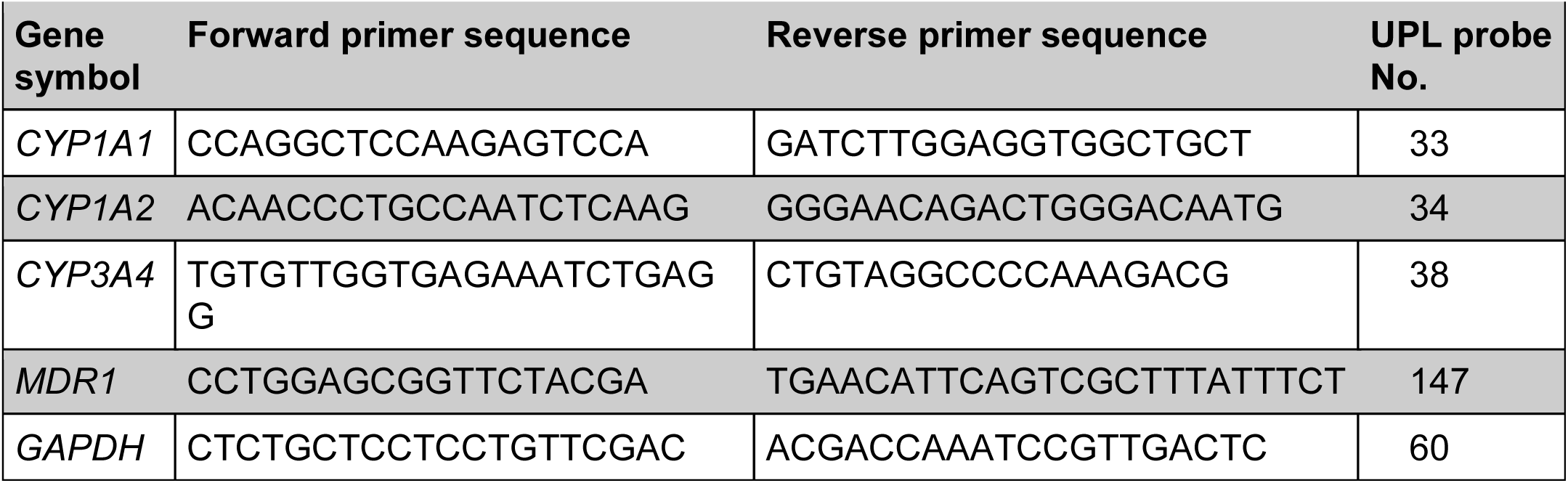

The following protocol was used: an activation step at 95 °C for 10 min was followed by 45 cycles of PCR (denaturation at 95 °C for 10 s; annealing with elongation at 60 °C for 30 s). The measurements were performed in triplicate. Gene expression was normalized *per GAPDH* as a housekeeping gene. The data were processed according to the comparative C_T_ method (Schmittgen & Livak, 2008). Data are expressed as fold induction ± SD over the vehicle-treated cells. The bar graph depicts one representative experiment of a series of experiments performed in three consecutive passages of cells. Differences were tested using one-way ANOVA with Dunnett’s post hoc test, *p* < 0.05 was considered significant.

In a separate set of experiments in Caco-2 cells, 1 × 10_5_ cells were transfected with pSG5-PXR in 6 well plates for 12 h, then treated with vehicle (0.1% DMSO), Rifampcin, FKK5 and FKK6 each at a concentration of 10 µM, respectively, in triplicate for 24 h. Cells were harvested for preparation of total RNA using TRIzol Reagent (ambion, #15596026, Carsbad, CA 92008) and reverse transcribed to cDNA using the High Capacity cDNA Reverse Transcription Kit (Thermo Fisher Scientific, #4368814, LT 02241). RT-qPCR was performed using Thermo Fisher PowerUp SYBR Green Master Mix (# A25742) and Thermo Fisher qPCR 7900HT. Each sample RT-qPCR was repeated in quadruplicate and normalized to internal control, GAPDH. The entire experiment was repeated at least 2 separate times. The primer sequences are noted below.

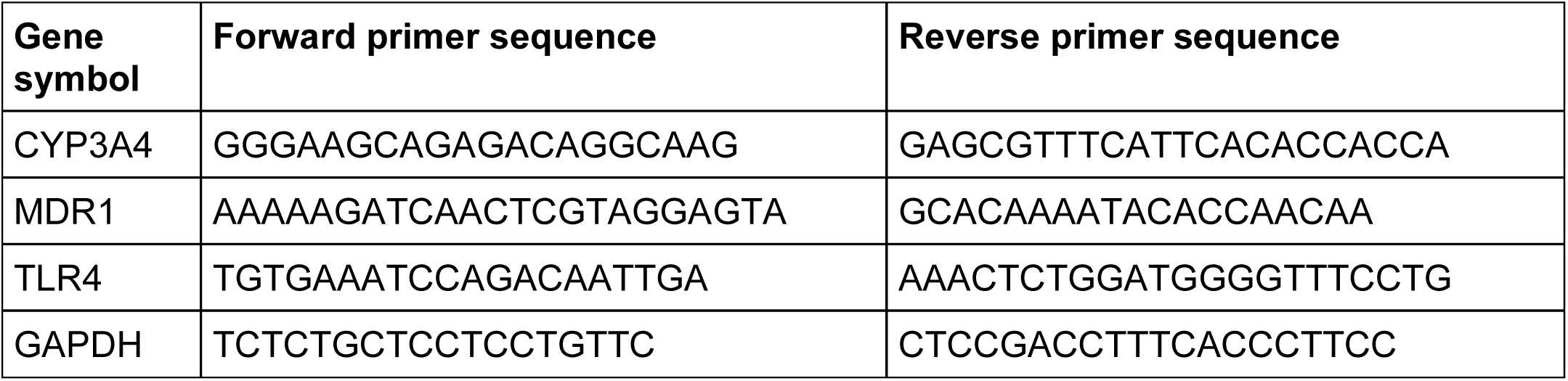

### Cytotoxicity Assays

For LS180 cells, the MTT colorimetric assay was performed as previously published (Bartonkova et al., 2015, Schiller, Kainz et al., 1992) and using established proliferation assay kits (Vybrant MTT Cell Proliferation Assay kit, Ref # V13154, lot # 1774057, Life Technologies).

### Chromatin Immunoprecipitation (ChIP) Assays

LS174T cells were plated in 150-mm cell culture dishes followed by transfection with pSG5-PXR (2 μg/plate/1 × 10_7_ cells). After 12 h, cells were exposed to vehicle (DMSO 0.1% v/v) or 10 μM FKK5, FKK6 or FKK9, respectively, for 24 h. Chromatin Immunoprecipitation (ChIP) was performed using previously published protocols(Nelson, Denisenko et al., 2006, Wang et al., 2011). Briefly, cells were chemically cross-linked with formaldehyde in culture media for 15 min at room temperature. Glycine was added to a final concentration of 0.125 M to stop cross-linking for 5 min at room temperature. Cells were rinsed twice with cold PBS (pH 7.5) and scraped in PBS supplemented with protease inhibitor cocktail (PIC).

Cell pellets were collected and immediately resuspended in immunoprecipitation (IP) buffer (150 mM NaCl, 50 mM Tris-HCl at pH 7.5, 5 mM EDTA, 0.5% NP-40, 1% Triton X-100). The nuclei were centrifuged at 12000 × *g* for 1 min and resuspended in fresh IP buffer. Chromatin was sheared by sonication to average size of 0.2-0.9 kb. Lysates were cleared by centrifugation at 12,000 × *g* for 10 min. Chromatin from 2 × 10_6_ cells (about 200 μl) was incubated overnight at 4 °C with the following 2 μg rabbit anti-PXR [sc-25381 (H-160), Santa Cruz Biotechnology, CA], or normal rabbit IgG (sc-2027, Santa Cruz Biotechnology, CA). Immune Complexes were captured with 20 µl of packed Protein A-Agarose (sc-2001, Santa Cruz Biotechnology) per IP for 1h at 4 °C and washed three times with IP buffer. Ten percent (10%) (wt/vol) Chelex 100 was added to the beads followed by boiling of the suspension and centrifugation at 12000 × *g* for 1 min to obtain chromatin.

Purified chromatin DNA was used to perform RT-qPCR. We used PowerUp SYBR Green Master Mix (A25742, Thermo Fisher Scientific, TX) in a 10 μl reaction, 0.5 μl DNA template, 0.5 μl primer pairs (10 μM each), 5 μl Master Mix and 3.5 μl H_2_O in 384-well plates on an ABI 7900, default three-step method, 40 cycles. Data was analyzed using SDS 2.2.1 program (ABI Biotechnology). We used the following primers for ChIP:

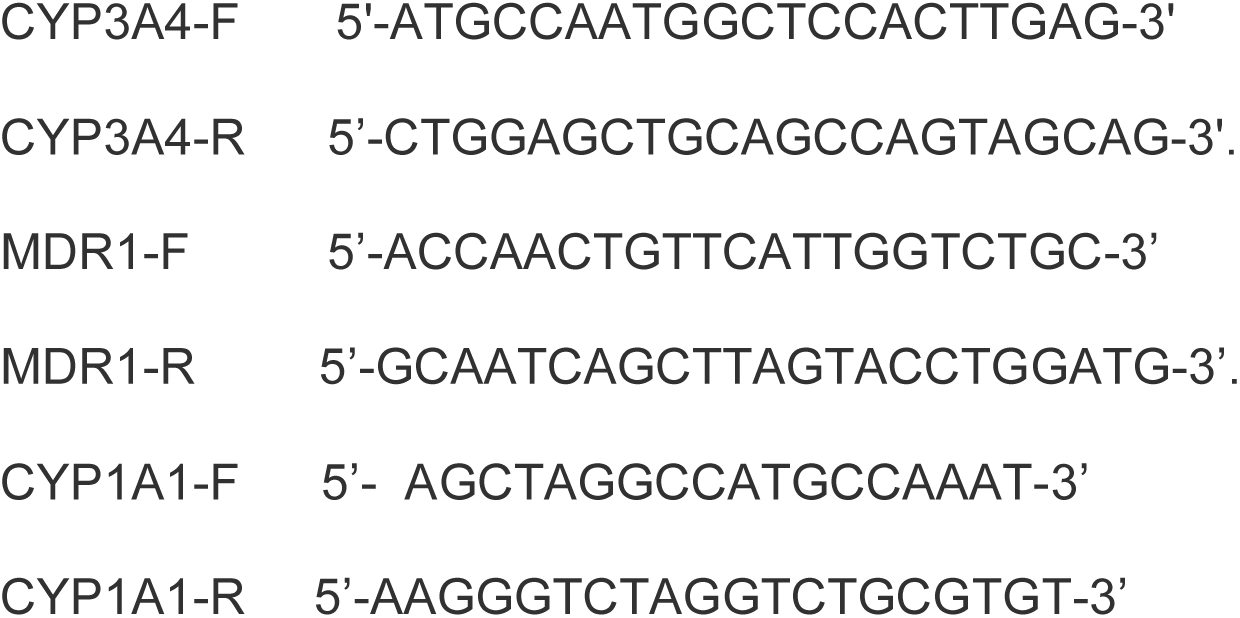

### Expression and purification of His-tagged PXR Ligand Binding Domain (LBD) protein

The cloning, expression and purification of His-tagged PXR LBD was performed as previously published (Wallace, Betts et al., 2013) with the following modifications in the stated protocol as follows. For protein expression, Luria-Bertani (LB) media was inoculated with a saturated culture of BL21-Gold cells transformed with HIS-LIC plasmid containing the PXR LBD construct. The mixture was allowed to shake at 37 *°*C until the cells reached an OD_600_ ∼ 0.6, and then the temperature was reduced to 18 °C, at which time IPTG was added (final concentration of 0.1 mM) to induce protein expression. For purification, the His-tag was not removed, and the un-cleaved protein was loaded onto the gel filtration column with buffer containing HEPES (25 mM, pH 7.5) and NaCl (150 mM).

### Isothermal titration calorimetry (ITC)

Isothermal titration calorimetry (ITC) was carried out using a VP – ITC microcalorimeter from MicroCal⁄Malvern Instruments (Northampton, MA, USA). The protein and its ligands were prepared in 25 mM Hepes, pH 7.5, with 150 mM NaCl and DMSO at a concentration of 6% for the experiment with FKK5 and FKK6 and 2% in the experiment with rifampicin and 3-IPA. In all the experiments the ligand solution was injected in 10-µL aliquots into the calorimetric cell containing PXR-LBD (ligand binding domain) at a concentration of 3 – 6 µM. The respective concentrations of FKK5, FKK6, rifampicin, and 3-IPA in the syringe were 60, 80, 330, and 400 µM. The experiments were carried out at 37 °C. The heat evolved upon each injection of the ligands was obtained from the integral of the calorimetric signal. The heat associated with binding to PXR-LBD in the cell was obtained by subtracting the heat of dilution from the heat of reaction. The individual heats were plotted against the molar ratio, and the enthalpy change (*ΔH*), association constant (*K_a_* = 1⁄*K_d_*), and the stoichiometry were obtained by nonlinear regression of the data. In the case of rifampicin, a binding model was chosen that took into account two sets of sites with different binding affinities.

### hPXR TR-FRET

The LanthaScreen TR-FRET PXR (SXR) Competitive Binding Assay Kit (PV4839; Invitrogen, USA). The PXR ligand binding assay of FKK5, FKK6, and FKK9 was performed using LanthaScreen TR-FRET PXR (SXR) Competitive Binding Assay Kit according to the manufacturer’s instructions. The assays were done in a volume of 20 μl in 384-well black plates containing different concentrations of tested compounds in the range of 1 nM to 25 μM. DMSO and 100μM SR12813 were used as a negative and positive control, respectively. The reaction mixture was incubated at room temperature for 1 hour in a dark, and then fluorescent signals were measured at 495 nm and 520 nm, with the excitation filter 340 nm, on Infinite F200 microplate reader (Tecan Group Ltd, Switzerland). Finally, the TR-FRET ratio was calculated by dividing the emission signal of 520 nm by that at 495 nm. All PXR binding assays were performed as two independent experiments, each with a minimum of four replicates. Final IC_50_ were obtained by processing the data with GraphPad Prism 6 using standard curve interpolation (sigmoidal, 4PL, variable slope). Since rifampicin, at higher concentrations, interferes with FRET signals in this assay (Shukla, Nguyen et al., 2009, Shukla, Sakamuru et al., 2011), SR12813 serves as a positive control PXR agonist ligand and it demonstrates an IC_50_ of 0.127 μM, which is similar to previously published results (Shukla et al., 2009).

### Human studies

Human tissues (duodenum), was collected according to standard research protocols approved by the Institutional Review Board and Department of Pathology at Cincinnati Children’s Hospital (IRB: 2014-6279; renewed 11/27/2017).

Additional samples were collected from consenting healthy patients undergoing colonoscopy at the University of Calgary endoscopy unit for colon cancer screening or to investigate gastrointestinal symptoms with normal colonoscopic appearance and normal histology (Study ID: REB18-0631_REN1). Biopsies via endoscope were taken from the colonic mucosa and immediately placed in Intesticult^TM^ Organoid Growth Media supplemented with antibiotic/antimycotic (Stemcell Technologies) and transferred from the unit to the lab.

### Isolation of crypts and culture of human enteroids from patient-derived duodenum (and RT-PCR assays)

Discarded duodenum tissue after surgery was obtained and used to isolate crypts. Embedded crypts in Matrigel formed three-dimentional (3D) structure referred to as ‘enteroids’. The isolation protocol has been described as previously (Mahe, Sundaram et al., 2015). Enteroids were incubated with 10 µM compounds (FKK5, FKK6, and Rifampicin) at day 4 from isolation. DMSO was added to enteroids as negative control. After incubation for 24 h, enteroids were collected in RNA-free tube followed by breaking down Matrigel by pipetting vigorously (using 1 mL PBS). The enteroids were pelleted at 16000 g for 3 min (4_o_C). RNA was extracted using miRNA isolation kit (Invitrogen; #AM1561) using the protocol provided by Invitrogen. The cDNA was synthesized from the extracted RNA (final amount of RNA: 1 µg) using SuperScript III (Invitrogen, #18080-051) through two-step procedure provided by Invitrogen. PowerUp SYBR Green (Applied Biosystem, #A25742) was used for RT-PCR and performed 40 cycles using Quant Studio3 (Invitrogen, #A28132). Primers (Invitrogen, #A15612)

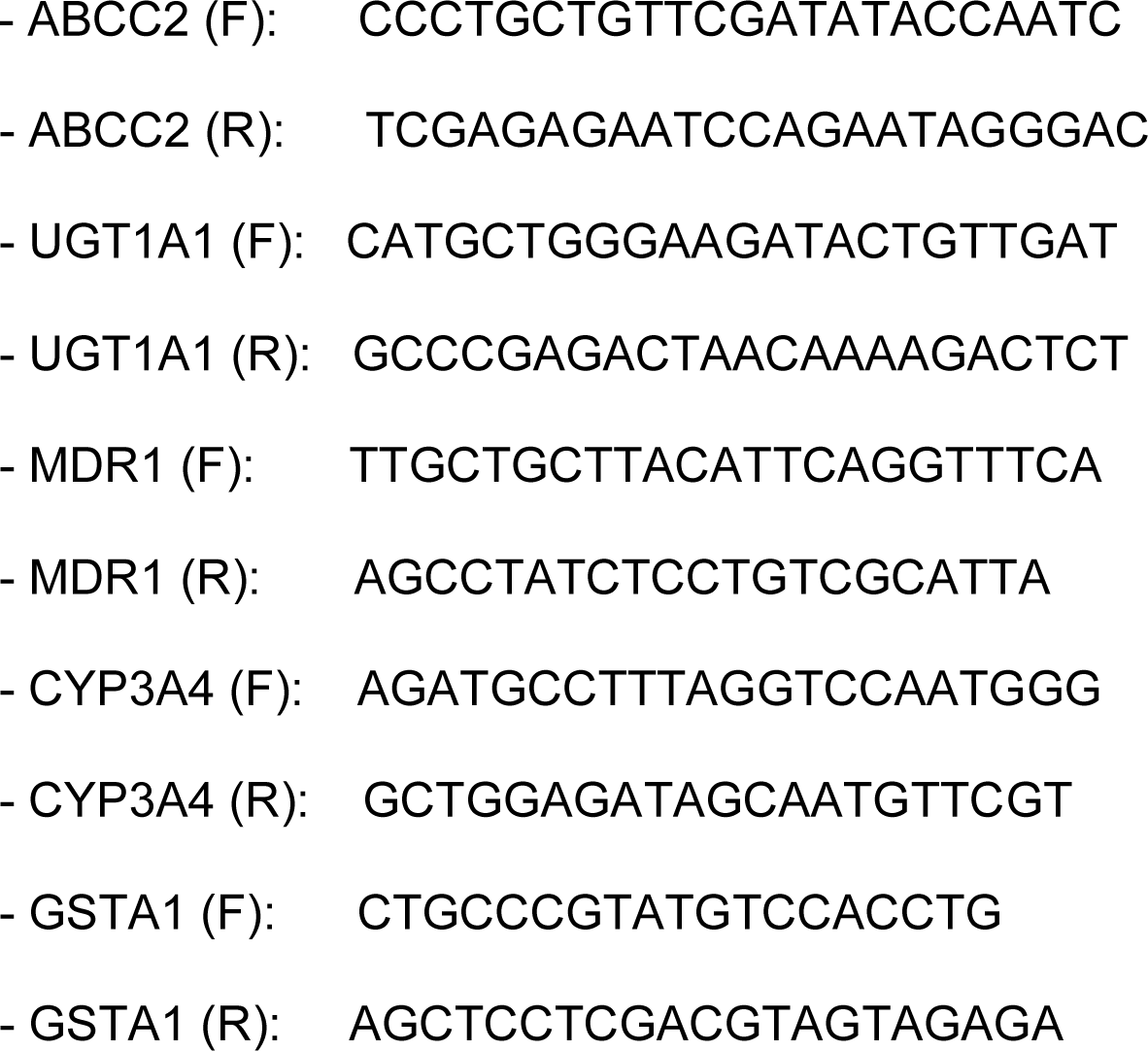

### Culture of human-derived colonic enteroids and FKK treatment assays (Calgary Protocol)

Samples were immediately processed to yield isolated colonic crypts as previously described (Fernando, Dicay et al., 2017). Isolated colonic crypts were embedded in Matrigel® (Corning) and cultured in Intesticult^TM^ Organoid Growth Media. Human organoids were pretreated for 6 h with either FKK5, FKK6, FKK9 or vehicle (DMSO) in Intesticult^TM^ Organoid Growth Media. Organoids were then stimulated with human recombinant TNFalpha in the presence of either treatment for an additional 12 h. After the 12 h treatment organoids were washed in PBS, disrupted in Tri Reagent® (Sigma, Oakville, ON, CAN) and frozen at −80°C. After thawing, chloroform was added, samples were centrifuged for 15 min and the resulting aqueous phase was then added to equal volumes of 70% ethanol and further processed using the RNeasy mini kit (Qiagen). cDNA was synthesized using the Quantitect RT kit (Qiagen) according to the manufacturers’ protocol. Resulting cDNA was used as a template for quantitative real-time PCR using Perfecta SYBR Green FastMix with ROX (QuantiBio). PCR and analysis were performed using a StepOne PCR system (Applied Biosystems). Gene expression was calculated relative to β-Actin expression and expressed as fold-change of control. The primers used were as follows: Human ®-Actin (NM_001101), Human CXCL8 (NM_000584)(data not reported), Human CYP3A4 (NM_001202855) and Human ABCB1 (NM_000927).

### Caco2 cell cultures and iPSC derived human intestinal organoids (and RT-PCR Assays)

Caco-2 cells (ATCC, Manassas, VA) passage 25–35 were maintained and expanded in culture flasks in Dulbecco’s Modified Eagle Medium (DMEM) with 10 % FBS and 1 × antibiotic-antimycotic. The cells were kept in a humidified 37 °C incubator with 5% CO_2_. Media was changed every 2-3 days and with regular passage 1-2 times a week. For all experiments, Caco-2 cells (5 × 10_4_ cells/mL) were seeded into 12 well Transwell inserts or coverslips and cultured for 3 weeks. HIOs were generated from iPSCs as described previously(Spence, Mayhew et al., 2011), with some modifications. The iPSC cell line IISH1i-BM1 (WiCell., Madison, WI) was cultured on Matrigel plates to 80% confluence in mTeSR™1 medium (Stem Cell Technologies., Seattle, WA) and differentiated to endoderm using Definitive Endoderm Kit (Stem Cell Technologies., Seattle, WA) for 4 days with daily media changes. Endoderm was then differentiated to hindgut for 5 days with the addition of FGF-4 (500 ng/mL; Peprotech., Rocky Hill, NJ) and Chir99021 (3 µM; Cayman., Ann Arbor, MI) in RPMI media supplemented with 2% FBS. Floating hindgut spheroids were collected on days 3−5 of hindgut differentiation and suspended in Matrigel beads for HIO differentiation. HIOs were maintained in WENRAS media (Fujii, Matano et al., 2015) which maintains proliferation in the culture. Stock HIOs were passaged weekly by removing Matrigel with Cell Recovery Solution (Corning, NY) and mechanically breaking up organoids with pipetting. HIOs were seeded into new Matrigel beads at a 1:3 split, using WENRAS with added Rock Inhibitor Y-27632 (10 µM) and Chir99021 (3 µM; Cayman., Ann Arbor, MI) for the first two days of culture.

HIOs and Caco-2 were exposed to a pro-inflammatory cytokine cocktail (10 ng/mL IFNγ, 10 ng/mL TNFα, 50 ng/mL IL-1β) for varying time points (2 − 24 h) in combination with DMSO control or FKK5 and FKK6 compounds (10 and 25 µm). Relative mRNA levels of IL-8 and IL-6 were determined using quantitative real time PCR, with normalization to GAPDH. RNA was isolated using an RNeasy Mini Kit (Qiagen) as per the manufacturer’s instructions. On column DNAse (Qiagen) was used to remove any contaminating DNA. Next, cDNA was formed using iScript cDNA Synthesis Kit (Bio-rad) and 100 ng RNA. Quantitative real-time PCR was performed with SsoAdvanced Universal SYBR Green kit (Bio-rad). The following primers were used: IL-8 Forward: ATACTCCAAACCTTTCCACCC; IL-8 Reverse: TCTGCACCCAGTTTTCCTTG; IL-6 Forward: CCACTCACCTCTTCAGAACG; IL-6 Reverse: CATCTTTGGAAGGTTCAGGTTG; GAPDH Forward: ACATCGCTCAGACACCAT; GAPDH Reverse: TGTAGTTGAGGTCAATGAAGGG. For immuno-fluorescent staining of NF-ĸB translocation to the cell nucleus, HIOs and Caco-2 cell monolayers were fixed in 4% formaldehyde (in 1 × PBS) overnight at 4 °C. The formaldehyde was removed by washing with 1 × PBS, and an antigen retrieval step was performed by incubating in 10 mM sodium-citrate buffer in a vegetable steamer for 20 min. The samples were then blocked with normal donkey serum (10 % in PBS with 0.3% Triton x-100) for one hour. Samples were incubated overnight with primary antibodies: rabbit anti-NF-κB p65 (D14E12 # 8242, 1:400 dilution, Cell Signaling Technology, Danvers MA). Samples were then immersed in Alexa Fluor® 555 donkey anti-rabbit secondary antibody (A-31572, Life Technologies) at a 1:500 dilution. Nuclei were stained with Hoechst 33342 (Life Technologies). The samples were then imaged using a Zeiss LSM880 Confocal/Multiphoton Upright Microscope, with 3-D image rendering using Volocity. Note, for controls, normal rabbit IgG (#2729S, Life Technologies) was used.

FKK compounds were tested for their ability to prevent *Salmonella* invasion in vitro in non-inflamed and inflamed cell cultures. Overnight cultures of *Salmonella typhimurium* 14038 were diluted to OD_600_ 0.01 and injected into with HIOs or immersed over Caco-2 monolayers for 1 hour at 37_o_C. The invasive ability of *Salmonella* was assessed using the gentamicin protection assay (Gagnon, Zihler Berner et al., 2013). Bacterial adhesion scenarios to the Caco-2 surface were set up as described above. Non-adhered bacteria were removed by washing twice in PBS, followed by incubation with 1 mL gentamicin (150 µg/mL _–1_ in DMEM) for 1 hour at 37_o_C to kill the adhered extracellular bacteria. Dead bacteria were removed by washing twice in PBS, followed by an incubation with 500 µl 0.1 % Triton X-100 for 15 min at 37_o_C to lyse the HIOs and Caco-2, and release the intracellular (invaded) bacteria. Serial fold dilutions and plating were then employed to determine CFU/mL. We normalized to total protein using Bradford Assay.

### Animal Studies

FKK6 was solubilized to saturation concentrations (∼ 500 μM) at room temperature in 10% DMSO. This concentration of DMSO is safe in mice, especially if administered in short course (maximum tolerated dose in mice is ∼ 2.5 gm/kg/day for 35 consecutive days) (Brayton, 1986, Caujolle, Caujolle et al., 1967a). The dose of 500 μM (∼1 mg/kg/day) was chosen based on the knowledge that despite poor solubility, some similar derivatives could have 100% bioavailability(Zhu, Sun et al., 2019); others in the same class, have a lower bioavailability (∼ 1%)(Li, Jin et al., 2019). Since we do not know the bioavailability of FKK6 in mice, a conservative guide is that 10-90% of the administered dose for any given mouse is absorbed. This would ensure the most mice would exhibit concentrations exceeding 50 μM in the feces, which would be higher than 10 μM concentrations used in cell culture.

Male and female C57BL/6J mice (Jackson Labs Stock 000664)(6-10 weeks), *pxr^-/-^* (6 −10 weeks) (Staudinger, Goodwin et al., 2001), and male and female mice expressing the human PXR gene (*hPXR*) [Taconic, C57BL/6-*Nr1i2_tm1(NR1I2)Arte_* (9104, females; 9104, males), 6-10 weeks] (Scheer, Ross et al., 2010) were administered either 10% DMSO (*n* = 3) or FKK6 (*n* = 3) in either 3 doses or 5 doses, with each dose interval of 12 h. Each gavage dose volume was 100 μL. At 2−4h after the end of the scheduled dosing period, mice were sacrificed, and tissues harvested for further analysis. The mouse studies were performed three independent times over an 18-month period. All purchased mice were acclimatized in the Einstein vivarium for ∼ 2 weeks before experiments commenced. All mice (n ≤ 5 per stainless steel cage and separated by gender) were maintained on standard non-irradiated chow (LabDiet 5058, Lab Supply) and sterile water in the barrier facility observing a 12 h night/day cycle. The room temperature was maintained at 25 ± 2°C with 55 ± 5% humidity. All studies were approved by the Institute of Animal Studies at the Albert Einstein College of Medicine, INC (IACUC # 20160706, 20160701 and preceding protocols) and specific animal protocols were also approved by additional protocols (IACUC# 20170406, 20170504) and Animal Care and Use Review Office (ACURO) of the US Army Medical Research and Materiel Command (PR160167, W81XWH-17-1-0479).

### Dextran Sulfate Sodium (DSS)-induced acute colitis studies

For these studies 7-8 week old female (DeVoss & Diehl, 2014) *hPXR* (mice expressing the human PXR gene) and *pxr^-/-^* mice (C57Bl/6 background) were bred and each genotype was separated into two treatment groups using a random allocation method via coin toss – (1) control (vehicle treated consisting 0.8% DMSO in 100 μL drinking water) administered once per day via oral gavage and via intrarectal gavage for a total of 10 consecutive days (*n* = 3) and (2) FKK6 in 0.8% DMSO at 200 μM (0.4 mg/kg/day) in 100 μL drinking water administered once per day via oral gavage and via intrarectal gavage for a total of 10 consecutive days (*n* = 3). The rationale for choosing this dose is based on the extrapolation of total dose delivered over 10 days to mice (∼ 4 mg/kg), which is equivalent to that delivered to mice for determining target gene expression over 3-5 doses (3-5 mg/kg). The sample size of n = 3 was chosen arbitrarily as means to explore if at least one mouse responded to FKK6. Three independent biological repeat experiments were performed to confirm our observations. Intrarectal delivery, at a 45° angle, was achieved using a 1 mL syringe attached to a 19G needle that had a polyethylene tubing (#427516, 0.58 mm O.D, Becton Dickinson, Sparks, MD) inserted to 3 cm into anal opening under very light anesthesia (1.5% isofluorane with nose cone) till no visible liquid was seen outside the anus. Fecal pellets were collected prior to initiation of the treatment protocol. All mice were first given their treatment allocation and 4 h later were all treated with 3% DSS (MPBio LLC, Salon, OH; Cat# 160110) in drinking water. All mice were starved overnight (12 h) to allow for smooth intrarectal delivery; however, the mice were allowed to feed *ad libitum* during the day. This protocol was continued for 10 consecutive days after which mice were sacrificed and organs/tissues/feces harvested for further study. On day 10, prior to necropsy, mice were administered FITC-dextran and mouse serum was collected for the FITC-dextran assay as previously described (Vetizou, Pitt et al., 2015). The experimenter was not blinded to treatment allocation; however, the pathologist (KS) evaluating histologic scores is blinded to treatment allocation. All studies were approved by the Institute of Animal Studies at the Albert Einstein College of Medicine, INC (IACUC # 20160706 and preceding protocols). Colitis scoring was performed as previously published and post-examination masking was conducted (Kim, Shajib et al., 2012, Meyerholz & Beck, 2018).

For RT-qPCR of colon tissue for cytokine expression data, new experiments were conducted with 3% DSS and the identical FKK6 dose (n = 3 mice/group) and schedule as above. RT-qPCR was performed using PowerUp^TM^ SYBR® Green Master Mix (A25741, ThermoFisher Scientific) and primers specific for IL10, TNFα, TNFR2, and IL7f. Each target gene was analyzed in quadruplicate. Normalizing controls included GAPDH (used to illustrate data) and TBP.

Primer sequences used:

TBP-Primers:

F: ACCGTGAATCTTGGCTGTAAAC

R: GCAGCAAATCGCTTGGGATTA

GAPDH-Primers:

F: AACTTTGGCATTGTGGAAGG

R: GGATGCAGGGATGATGTTCT

IL10-Primers:

F: TGAGGCGCTGTCGTCATCGATTTCTCCC

R: ACCTGCTCCACTGCCTTGCT

TNFR2-Primers:

F: CAGGTTGTCTTGACACCCTAC

R: GCACAGCACATCTGAGCCT

TNFα-Primers:

F: ATGAGAAGTTCCCAAATGGC

R: AGCTGCTCCTCCACTTGGTGG

IL17f-Primers:

F: TGCTACTGTTGATGTTGGGAC

R: AATGCCCTGGTTTTGGTTGAA

### Statistical Analysis

All data sets were initially visually inspected and then assessed for normality and lognormality, descriptive and outlier analysis using tests specified in GraphPad Prism 8.2.0 (272) (except as noted). Student *t*-test, one-way analysis of variance (1-Way ANOVA), two-way analysis of variance (2-Way ANOVA) followed by Dunnett test, as well as values of EC_50_ and IC_50_, was calculated using GraphPad Prism version 8.2.0 (272) for Windows (GraphPad Software, La Jolla, California, USA) are indicated as appropriate in each figure legend. The statistical analysis of FKK6 treatment end-points (% weight loss, colon length, lipocalin 2, and FITC-dextran) in *pxr^+/+^* and *pxr^-/-^* was performed using student *t*-test and non-parametric tests for non-normal data as well as by clustering clustering method on the six sample data vector to split the mice into two groups. The “kmeans” function of the R software was used to obtain the clusters.

## DATA AVAILABILITY

The FKK5-PXR docking model has been submitted to BioModels with identifier: MODEL1912090005.

The immunofluorescence images have been submitted to BioImage Archives with identifier: S-BSST310

FKK5 and FKK6 structures have been deposited in the Cambridge Structural Database under deposition numbers 1948848 and 1948849, respectively.

## AUTHOR INFORMATION

### Corresponding Author(s)

** Phone : (718) 430-2871 Fax: (718) 430-8550

** Phone: (215) 991-8135 Fax: (215) 848-2271

** Phone: + 420 585634903 Fax: n/a

### Author Contributions

The manuscript was written through contributions of all authors. All authors have given approval to the final version of the manuscript. The following are the specific contributions made:

Zdeněk Dvořák^1^‡* (conceptual and experimental design, data interpretation of [TR-FRET, gene expression in cell lines, HepaRG and human hepatocytes, reporter gene assays (PXR, AHR, VDR, TR, GR, AR, NFκB), provided materials, helped in paper writing.]

Felix Kopp^2^‡ (originated and synthesized FKK1-10 chemicals).

Cait M. Costello^17^ (Caco-2 and HIO experiments, salmonella invasion studies)

Jazmin S. Kemp^17^ (Caco-2 and HIO experiments, salmonella invasion studies)

Hao Li^3^‡ (all mouse studies, select transfection studies, ChIP assay)

Aneta Vrzalová^1^‡ [experimental assays, data acquisition, data evaluation of reporter gene assays (PXR, AHR), gene expression in intestinal cell lines (transfections, treatments, RT-PCR) and in human hepatocytes cultures Hep200529+Hep220932]

Martina Štěpánková^1^ (experimental, data acquisition, data evaluation of gene expression in HepaRG and human hepatocytes cultures Hep200533+Hep200538 (RT-PCR)]

Iveta Bartoňková^1^ [experimental, data acquisition, data evaluation of reporter gene assays (PXR, AHR, VDR, TR, AR, GR, NFκB); HepaRG (WT, PXR-KO, AHR-KO) of cultivation, differentiation, RT-PXR analyses, western blots for validation of receptors expression; validation of LS174 WT/PXR-KO cells using RT-PCR; human hepatocytes cultures Hep200533+Hep200538 (RT-PCR)]

Eva Jiskrová^1^ (general experimental, data acquisition, data evaluation of TR-FRET / PXR)

Karolína Poulíková^1^ [experimental, data acquisition, data evaluation of reporter gene assays in intestinal cells (NFκB, co-transfected PXR), mock experiments for PXR and AHR, protein levels by SallySue)]

Barbora Vyhlídalová^1^ [experimental, data acquisition, data evaluation of reporter gene assays in intestinal cells (NFκB, co-transfected PXR), mock experiments for PXR and AHR, protein levels by SallySue]

Lars U. Nordstroem^2^ (performed and developed crystals of FKK5 and FKK6 in solution)

Chamini Karunaratne^2^ (scale up synthesis of some FKK compounds)

Harmit Ranhotra^3, $^ (reverse transfection studies with PXR)

Kyu Shik Mun^5^ (generated organoid data in Cincinnati)

Anjaparavanda P. Naren^5^ (conceptual and experimental design of organoid data generated in Cincinnati)

Iain Murray^6,^ Gary H. Perdew^6^ (externally validated effects of FKK compounds on AhR reporter assays)

Julius Brtko^7^, Lucia Toporova^7^ (experimental design, data acquisition, data evaluation of the radioligand binding assay for RXR)

Arne Schon^8^ (performed and analyzed Isothermic Titration Calorimetry/ITC studies)

Bret Wallace^9^, William G. Walton^9^, Matthew R. Redinbo^9^ (generated soluble PXR protein for ITC)

Katherine Sun^10^, Amanda Beck^4^ (performed blinded pathology readings)

Sandhya Kortagere^11^* (performed PXR docking studies)

Michelle C. Neary^12^ (performed X-ray diffraction analysis of FKK compounds in solid state)

Aneesh Chandran^13^, Saraswathi Vishveshwara^13^ (performed PXR docking studies)

Maria M. Cavalluzzi^14^, Giovanni Lentini^14^ (performed LEM analysis)

Julia Yue Cui^15^, Haiwei Gu^16^ (developed extraction methods for fecal analysis of FKK5 and 6 and MS verification of compound)

John C. March^17^ (Caco-2 and HIO experimental design and setup, salmonella invasion studies)

Shirshendu Chaterjee^18^ (statistical analysis for all studies performed)

Adam Matson^19^, Dennis Wright^20^ (expert advice on animal models pertinent to barrier dysfunction and on FKK compound synthesis)

Kyle L. Flannigan^21^, Simon A. Hirota^21^ (performed organoid studies in Calgary).

Ryan Balfour Sartor^22^ (helped with identification of cytokines for study and interpretation of DSS model)

Sridhar Mani^3,^* (initial conceptual design of microbial metabolite mimicry, planned all the studies presented in the paper, developed the team and troubleshooting of all experiments presented)

‡These authors contributed equally to this work.

### Additional Information (Competing and Conflict of Interest Statement)

The studies presented here are included in a patent submitted by The Albert Einstein College of Medicine in conjunction with Palacký University and The Drexel University College of Medicine to the US Patent and Trademark Office. Funding for these studies are listed under acknowledgement.

## ACKNOWLEDGMENTS

The studies presented here were funded in part by the ICTR Pilot Award (AECOM to S.M & F.K); Grant# 362520 Broad Medical Research Program (BMRP, not Litwin) at CCFA (Crohn’s & Colitis Foundation of America) (to S.M); NIH grants R35 ES028244 (to G.H.P); CA127231; CA 161879 and Department of Defense Partnering PI (W81XWH-17-1-0479; PR160167) (to S.M.), (ES030197) (to S.M.,J.C.,H.G) as well as R43DK105694 (PI: Jay Wrobel), P30DK041296 (PI: Alan Wolkoff) (Pilot Awards, S.M); Diabetes Research Center Grant (P30 DK020541); Cancer Center Grant (P30CA013330 PI: David Goldman); 1S10OD019961-01 NIH Instrument Award (PI: John Condeelis); LTQ Orbitrap Velos Mass Spectrometer System (1S10RR029398); and NIH CTSA (1 UL1 TR001073). Additional invaluable assistance was obtained from Vera DesMarais PhD (Light Microscopy and Image Analysis Analytical Imaging Facility (AIF) Albert Einstein College of Medicine, Bronx, NY), Amanda Beck DVM (Histology and Comparative Pathology Core, Albert Einstein College of Medicine, Bronx, NY), Lars Nordstroem PhD and Chamini Karunaratne PhD (Chemical Synthesis and Biology Core, Albert Einstein College of Medicine, Bronx, NY), Yungping Qiu PhD (Stable Isotopes and Metabolomics Core Facility, Albert Einstein College of Medicine, Bronx, NY), and the Proteomics Core Facility, Albert Einstein College of Medicine, Bronx, NY, and The Czech Science Foundation [19-00236S] and the Operational Programme Research, Development and Education – European Regional Development Fund, the Ministry of Education, Youth and Sports of the Czech Republic [CZ.02.1.01/0.0/0.0/16_019/0000754](Z.D.), and Christian Jobin (*E. coli* NC101 strains, University of Florida, Gainesville, FL). Arpan De performed the biofilm assays. MMC and GL are indebted to professor Cele Abad-Zapatero who invented AtlasCBS and disclosed its potential as a tool to facilitate drug design and development endeavors. SV and AC are supported by the National Academy of Sciences, India. The authors thank Ms. Gurmeet Bindra in the inflammatory intestinal tissue bank (IITB) and the staff at the University of Calgary endoscopy unit for assistance with research sample collection and Dr. Marilyn Gordon of the Human Tissue Research Lab at the University of Calgary for sample preparation and human intestinal organoid protocol optimization.

**ABBREVIATIONS USED**
PXR: Pregnane X Receptor
SXR: steroid and xenobiotic receptor
FKK: Felix Kopp Kortagere
DMSO: dimethyl sulfoxide
AhR: aryl hydrocarbon receptor
GR: glucocorticoid receptor
VDR: vitamin D receptor
VDRE: VDR elements (DNA binding)
TR: thyroid receptor
AR: androgen receptor
RT: reverse transcription
qPCR: quantitative polymerase chain reaction
TLR4: toll-like receptor 4
GLP-1: glucagon-like peptide-1
IPA: Indole 3-propionic acid
Å: angstrom
TCDD: 2,3,7,8 – tetrachlorodibenzo-*p*-dioxin
T3: Triiodo-L-Thyronine
DHT: Dihydrotestosterone
KO: knockout
1α,25-VD3: calcitriol
DEX: dexamethasone
ADMET: absorption, distribution, metabolism, excretion, toxicology
STR: short tandem repeats
DNA: deoxyribonucleic acid
rpm: revolutions per minute
NaCl: sodium chloride
HEPES: 4-(2-hydroxyethyl)-1-piperazineethanesulfonic acid
EDTA: ethylenediaminetetra acetic acid
SDS–PAGE: sodium dodecyl sulfate polyacrylamide gel electrophoresis
CZ: Czech Republic
US: United States of America
PBS: Phosphate Buffered Saline
PIC: protease inhibitor cocktail
×: *g* times gravity
MTT: 3-(4,5-dimethylthiazolyl-2)-2,5-diphenyltetrazolium bromide)

## Relevant Links

https://www.nature.com/articles/nature23874

This paper provides direct evidence of a microbial metabolite that mimics a host response via a specific host receptor and forms the basis for host molecule mimicry.

https://www.sciencedirect.com/science/article/pii/S1359644616300356?via%3Dihub

This review paper positions that a new drug discovery class could be developed using microbial metabolite mimicry

https://plu.mx/plum/a/?doi=10.1016/j.drudis.2016.02.009&theme=plum-sciencedirect-theme&hideUsage=true

This twitter feed of the review paper lays some theoretical basis for this to occur

## REFERENCES

Abad-Zapatero C, Blasi D (2011) Ligand Efficiency Indices (LEIs): More than a Simple Efficiency Yardstick. Molecular informatics 30: 122–32

Abdel-Razik A, Mousa N, Shabana W, Refaey M, Elzehery R, Elhelaly R, Zalata K, Abdelsalam M, Eldeeb AA, Awad M, Elgamal A, Attia A, El-Wakeel N, Eldars W (2018) Rifaximin in nonalcoholic fatty liver disease: hit multiple targets with a single shot. Eur J Gastroenterol Hepatol 30: 1237–1246

Andersson TB (2010) The application of HepRG cells in evaluation of cytochrome P450 induction properties of drug compounds. Methods in molecular biology (Clifton, NJ) 640: 375–87

Aninat C, Piton A, Glaise D, Le Charpentier T, Langouet S, Morel F, Guguen-Guillouzo C, Guillouzo A (2006) Expression of cytochromes P450, conjugating enzymes and nuclear receptors in human hepatoma HepaRG cells. Drug metabolism and disposition: the biological fate of chemicals 34: 75–83

Antherieu S, Chesne C, Li R, Guguen-Guillouzo C, Guillouzo A (2012) Optimization of the HepaRG cell model for drug metabolism and toxicity studies. Toxicology in vitro : an international journal published in association with BIBRA 26: 1278–85

Bartonkova I, Grycova A, Dvorak Z (2016) Profiling of Vitamin D Metabolic Intermediates toward VDR Using Novel Stable Gene Reporter Cell Lines IZ-VDRE and IZ-CYP24. Chemical research in toxicology 29: 1211–22

Bartonkova I, Novotna A, Dvorak Z (2015) Novel stably transfected human reporter cell line AIZ-AR as a tool for an assessment of human androgen receptor transcriptional activity. PloS one 10: e0121316

Berkhout TA, Simon HM, Patel DD, Bentzen C, Niesor E, Jackson B, Suckling KE (1996) The novel cholesterol-lowering drug SR-12813 inhibits cholesterol synthesis via an increased degradation of 3-hydroxy-3-methylglutaryl-coenzyme A reductase. The Journal of biological chemistry 271: 14376–82

Biber A, Fischer H, Romer A, Chatterjee SS (1998) Oral bioavailability of hyperforin from hypericum extracts in rats and human volunteers. Pharmacopsychiatry 31 Suppl 1: 36–43

Bischoff SC, Barbara G, Buurman W, Ockhuizen T, Schulzke JD, Serino M, Tilg H, Watson A, Wells JM (2014) Intestinal permeability--a new target for disease prevention and therapy. BMC gastroenterology 14: 189

Brauze D, Zawierucha P, Kiwerska K, Bednarek K, Oleszak M, Rydzanicz M, Jarmuz-Szymczak M (2017) Induction of expression of aryl hydrocarbon receptor-dependent genes in human HepaRG cell line modified by shRNA and treated with beta-naphthoflavone. Molecular and cellular biochemistry 425: 59–75

Brave M, Lukin DJ, Mani S (2015) Microbial control of intestinal innate immunity. Oncotarget 6: 19962–3

Brayton CF (1986) Dimethyl sulfoxide (DMSO): a review. The Cornell veterinarian 76: 61–90

Castro CA, Hogan JB, Benson KA, Shehata CW, Landauer MR (1995) Behavioral effects of vehicles: DMSO, ethanol, Tween-20, Tween-80, and emulphor-620. Pharmacology, biochemistry, and behavior 50: 521–6

Caujolle FM, Caujolle DH, Cros SB, Calvet MM (1967a) Limits of toxic and teratogenic tolerance of dimethyl sulfoxide. Ann N Y Acad Sci 141: 110–26

Caujolle FME, Caujolle DH, Cros SB, Calvet M-MJ (1967b) LIMITS OF TOXIC AND TERATOGENIC TOLERANCE OF DIMETHYL SULFOXIDE. Annals of the New York Academy of Sciences 141: 110–125

Cavalluzzi MM, Mangiatordi GF, Nicolotti O, Lentini G (2017) Ligand efficiency metrics in drug discovery: the pros and cons from a practical perspective. Expert opinion on drug discovery 12: 1087–1104

Chappell CL, Darkoh C, Shimmin L, Farhana N, Kim D-K, Okhuysen PC, Hixson J (2016) Fecal Indole as a Biomarker of Susceptibility to Cryptosporidium Infection. Infection and immunity 84: 2299–2306

Cheng J, Krausz KW, Tanaka N, Gonzalez FJ (2012a) Chronic exposure to rifaximin causes hepatic steatosis in pregnane X receptor-humanized mice. Toxicol Sci 129: 456–68

Cheng J, Shah YM, Gonzalez FJ (2012b) Pregnane X receptor as a target for treatment of inflammatory bowel disorders. Trends in pharmacological sciences 33: 323–30

Cobbold JFL, Atkinson S, Marchesi JR, Smith A, Wai SN, Stove J, Shojaee-Moradie F, Jackson N, Umpleby AM, Fitzpatrick J, Thomas EL, Bell JD, Holmes E, Taylor-Robinson SD, Goldin RD, Yee MS, Anstee QM, Thursz MR (2018) Rifaximin in non-alcoholic steatohepatitis: An open-label pilot study. Hepatol Res 48: 69–77

Delfosse V, Dendele B, Huet T, Grimaldi M, Boulahtouf A, Gerbal-Chaloin S, Beucher B, Roecklin D, Muller C, Rahmani R, Cavailles V, Daujat-Chavanieu M, Vivat V, Pascussi JM, Balaguer P, Bourguet W (2015) Synergistic activation of human pregnane X receptor by binary cocktails of pharmaceutical and environmental compounds. Nature communications 6: 8089

Delie F, Rubas W (1997) A human colonic cell line sharing similarities with enterocytes as a model to examine oral absorption: advantages and limitations of the Caco-2 model. Critical reviews in therapeutic drug carrier systems 14: 221–86

DeVoss J, Diehl L (2014) Murine models of inflammatory bowel disease (IBD): challenges of modeling human disease. Toxicol Pathol 42: 99–110

Farrell DJ (2013) Rifaximin in the treatment of irritable bowel syndrome: is there a high risk for development of antimicrobial resistance? J Clin Gastroenterol 47: 205–11

Feisst C, Werz O (2004) Suppression of receptor-mediated Ca2+ mobilization and functional leukocyte responses by hyperforin. Biochemical pharmacology 67: 1531–9

Fernando EH, Dicay M, Stahl M, Gordon MH, Vegso A, Baggio C, Alston L, Lopes F, Baker K, Hirota S, McKay DM, Vallance B, MacNaughton WK (2017) A simple, cost-effective method for generating murine colonic 3D enteroids and 2D monolayers for studies of primary epithelial cell function. Am J Physiol Gastrointest Liver Physiol 313: G467–g475

Freire E (2008) Do enthalpy and entropy distinguish first in class from best in class? Drug discovery today 13: 869–74

Fujii M, Matano M, Nanki K, Sato T (2015) Efficient genetic engineering of human intestinal organoids using electroporation. Nature protocols 10: 1474–85

Gagnon M, Zihler Berner A, Chervet N, Chassard C, Lacroix C (2013) Comparison of the Caco-2, HT-29 and the mucus-secreting HT29-MTX intestinal cell models to investigate Salmonella adhesion and invasion. Journal of microbiological methods 94: 274–9

Galeotti N, Vivoli E, Bilia AR, Bergonzi MC, Bartolini A, Ghelardini C (2010) A prolonged protein kinase C-mediated, opioid-related antinociceptive effect of st John’s Wort in mice. J Pain 11: 149–59

Garbett NC, Chaires JB (2012) Thermodynamic studies for drug design and screening. Expert opinion on drug discovery 7: 299–314

Goodwin B, Hodgson E, Liddle C (1999) The orphan human pregnane X receptor mediates the transcriptional activation of CYP3A4 by rifampicin through a distal enhancer module. Mol Pharmacol 56: 1329–39

Gripon P, Rumin S, Urban S, Le Seyec J, Glaise D, Cannie I, Guyomard C, Lucas J, Trepo C, Guguen-Guillouzo C (2002) Infection of a human hepatoma cell line by hepatitis B virus. Proceedings of the National Academy of Sciences 99: 15655–15660

Guo Y, Zong S, Pu Y, Xu B, Zhang T, Wang B (2018) Advances in Pharmaceutical Strategies Enhancing the Efficiencies of Oral Colon-Targeted Delivery Systems in Inflammatory Bowel Disease. Molecules 23

Gupta A, Mugundu GM, Desai PB, Thummel KE, Unadkat JD (2008) Intestinal human colon adenocarcinoma cell line LS180 is an excellent model to study pregnane X receptor, but not constitutive androstane receptor, mediated CYP3A4 and multidrug resistance transporter 1 induction: studies with anti-human immunodeficiency virus protease inhibitors. Drug metabolism and disposition: the biological fate of chemicals 36: 1172–80

Huang H, Wang H, Sinz M, Zoeckler M, Staudinger J, Redinbo MR, Teotico DG, Locker J, Kalpana GV, Mani S (2007) Inhibition of drug metabolism by blocking the activation of nuclear receptors by ketoconazole. Oncogene 26: 258–68

Hung TV, Suzuki T (2018) Short-Chain Fatty Acids Suppress Inflammatory Reactions in Caco-2 Cells and Mouse Colons. Journal of Agricultural and Food Chemistry 66: 108–117

Hwang IK, Yoo KY, Li H, Park OK, Lee CH, Choi JH, Jeong YG, Lee YL, Kim YM, Kwon YG, Won MH (2009) Indole-3-propionic acid attenuates neuronal damage and oxidative stress in the ischemic hippocampus. Journal of neuroscience research 87: 2126–37

Illes P, Brtko J, Dvorak Z (2015) Development and Characterization of a Human Reporter Cell Line for the Assessment of Thyroid Receptor Transcriptional Activity: A Case of Organotin Endocrine Disruptors. Journal of agricultural and food chemistry 63: 7074–83

Kandel BA, Thomas M, Winter S, Damm G, Seehofer D, Burk O, Schwab M, Zanger UM (2016) Genomewide comparison of the inducible transcriptomes of nuclear receptors CAR, PXR and PPARalpha in primary human hepatocytes. Biochimica et biophysica acta 1859: 1218–27

Kim JJ, Shajib MS, Manocha MM, Khan WI (2012) Investigating intestinal inflammation in DSS-induced model of IBD. J Vis Exp

Kubesova K, Doricakova A, Travnicek Z, Dvorak Z (2016) Mixed-ligand copper(II) complexes activate aryl hydrocarbon receptor AhR and induce CYP1A genes expression in human hepatocytes and human cell lines. Toxicology letters 255: 24–35

Kublbeck J, Anttila T, Pulkkinen JT, Honkakoski P (2015) Improved assays for xenosensor activation based on reverse transfection. Toxicology in vitro : an international journal published in association with BIBRA 29: 1759–65

La Regina G, Bai R, Coluccia A, Famiglini V, Pelliccia S, Passacantilli S, Mazzoccoli C, Ruggieri V, Verrico A, Miele A, Monti L, Nalli M, Alfonsi R, Di Marcotullio L, Gulino A, Ricci B, Soriani A, Santoni A, Caraglia M, Porto S et al. (2015) New Indole Tubulin Assembly Inhibitors Cause Stable Arrest of Mitotic Progression, Enhanced Stimulation of Natural Killer Cell Cytotoxic Activity, and Repression of Hedgehog-Dependent Cancer. Journal of medicinal chemistry 58: 5789–5807

Li J, Jin Y, Fu H, Huang Y, Wang X, Zhou Y (2019) Pharmacokinetics and bioavailability of gelsenicine in mice by UPLC-MS/MS. Biomed Chromatogr 33: e4418

Mahe MM, Sundaram N, Watson CL, Shroyer NF, Helmrath MA (2015) Establishment of human epithelial enteroids and colonoids from whole tissue and biopsy. J Vis Exp

Mani S (2016a) Chapter 23 – Regulation of Host Chromatin by Bacterial Metabolites. In Chromatin Signaling and Diseases, Binda O, Fernandez-Zapico ME (eds) pp 423–442. Boston: Academic Press

Mani S (2016b) Chapter 23 – Regulation of Host Chromatin by Bacterial Metabolites A2 – Binda, Olivier. In Chromatin Signaling and Diseases, Fernandez-Zapico ME (ed) pp 423–442. Boston: Academic Press

Mani S (2017) Indole microbial metabolites: expanding and translating target(s). Oncotarget 8: 52014–52015

Mastropietro G, Tiscornia I, Perelmuter K, Astrada S, Bollati-Fogolin M (2015) HT-29 and Caco-2 reporter cell lines for functional studies of nuclear factor kappa B activation. Mediators of inflammation 2015: 860534

McCue PP, Phang JM (2008) Identification of human intracellular targets of the medicinal Herb St. John’s Wort by chemical-genetic profiling in yeast. J Agric Food Chem 56: 11011–7

Meng Z, Gwag T, Sui Y, Park SH, Zhou X, Zhou C (2019) The atypical antipsychotic quetiapine induces hyperlipidemia by activating intestinal PXR signaling. JCI insight 4

Meyerholz DK, Beck AP (2018) Principles and approaches for reproducible scoring of tissue stains in research. Laboratory Investigation 98: 844–855

Murray IA, Flaveny CA, DiNatale BC, Chairo CR, Schroeder JC, Kusnadi A, Perdew GH (2010) Antagonism of aryl hydrocarbon receptor signaling by 6,2’,4’-trimethoxyflavone. The Journal of pharmacology and experimental therapeutics 332: 135–44

Negres S, Scutari C, Ionica FE, Gonciar V, Velescu BS, Seremet OC, Zanfirescu A, Zbarcea CE, Stefanescu E, Ciobotaru E, ChiriTa C (2016) Influence of hyperforin on the morphology of internal organs and biochemical parameters, in experimental model in mice. Rom J Morphol Embryol 57: 663–673

Nelson JD, Denisenko O, Bomsztyk K (2006) Protocol for the fast chromatin immunoprecipitation (ChIP) method. Nature protocols 1: 179–85

Ning L, Lou X, Zhang F, Xu G (2019) Nuclear Receptors in the Pathogenesis and Management of Inflammatory Bowel Disease. Mediators of inflammation 2019: 2624941

Novotna A, Pavek P, Dvorak Z (2011) Novel stably transfected gene reporter human hepatoma cell line for assessment of aryl hydrocarbon receptor transcriptional activity: construction and characterization. Environmental science & technology 45: 10133–9

Novotna A, Pavek P, Dvorak Z (2012) Construction and characterization of a reporter gene cell line for assessment of human glucocorticoid receptor activation. European journal of pharmaceutical sciences : official journal of the European Federation for Pharmaceutical Sciences 47: 842–7

Orans J, Teotico DG, Redinbo MR (2005) The nuclear xenobiotic receptor pregnane X receptor: recent insights and new challenges. Mol Endocrinol 19: 2891–900

Parent R, Marion MJ, Furio L, Trepo C, Petit MA (2004) Origin and characterization of a human bipotent liver progenitor cell line. Gastroenterology 126: 1147–56

Pastorkova B, Vrzalova A, Bachleda P, Dvorak Z (2017) Hydroxystilbenes and methoxystilbenes activate human aryl hydrocarbon receptor and induce CYP1A genes in human hepatoma cells and human hepatocytes. Food and chemical toxicology : an international journal published for the British Industrial Biological Research Association 103: 122–132

Ranhotra HS, Flannigan KL, Brave M, Mukherjee S, Lukin DJ, Hirota SA, Mani S (2016) Xenobiotic Receptor-Mediated Regulation of Intestinal Barrier Function and Innate Immunity. Nuclear receptor research 3:1–19

Ruben AJ, Kiso Y, Freire E (2006) Overcoming roadblocks in lead optimization: a thermodynamic perspective. Chemical biology & drug design 67: 2–4

Saha S, Rajpal DK, Brown JR (2016) Human microbial metabolites as a source of new drugs. Drug discovery today 21: 692–8

Scheer N, Ross J, Kapelyukh Y, Rode A, Wolf CR (2010) In vivo responses of the human and murine pregnane X receptor to dexamethasone in mice. Drug metabolism and disposition: the biological fate of chemicals 38: 1046–53

Schiller CD, Kainz A, Mynett K, Gescher A (1992) Assessment of viability of hepatocytes in suspension using the MTT assay. Toxicology in vitro : an international journal published in association with BIBRA 6: 575–8

Schmittgen TD, Livak KJ (2008) Analyzing real-time PCR data by the comparative C(T) method. Nature protocols 3: 1101–8

Shan L (2004) (11)C-Labeled rifampicin. In Molecular Imaging and Contrast Agent Database (MICAD), Bethesda (MD): National Center for Biotechnology Information (US)

Shukla SJ, Nguyen DT, Macarthur R, Simeonov A, Frazee WJ, Hallis TM, Marks BD, Singh U, Eliason HC, Printen J, Austin CP, Inglese J, Auld DS (2009) Identification of pregnane X receptor ligands using time-resolved fluorescence resonance energy transfer and quantitative high-throughput screening. Assay Drug Dev Technol 7: 143–69

Shukla SJ, Sakamuru S, Huang R, Moeller TA, Shinn P, Vanleer D, Auld DS, Austin CP, Xia M (2011) Identification of clinically used drugs that activate pregnane X receptors. Drug metabolism and disposition: the biological fate of chemicals 39: 151–9

Smith RP, Eckalbar WL, Morrissey KM, Luizon MR, Hoffmann TJ, Sun X, Jones SL, Force Aldred S, Ramamoorthy A, Desta Z, Liu Y, Skaar TC, Trinklein ND, Giacomini KM, Ahituv N (2014) Genome-wide discovery of drug-dependent human liver regulatory elements. PLoS Genet 10: e1004648

Smutny T, Mani S, Pavek P (2013) Post-translational and post-transcriptional modifications of pregnane X receptor (PXR) in regulation of the cytochrome P450 superfamily. Current drug metabolism 14: 1059–69

Soleymani S, Bahramsoltani R, Rahimi R, Abdollahi M (2017) Clinical risks of St John’s Wort (Hypericum perforatum) co-administration. Expert opinion on drug metabolism & toxicology 13: 1047–1062

Spence JR, Mayhew CN, Rankin SA, Kuhar MF, Vallance JE, Tolle K, Hoskins EE, Kalinichenko VV, Wells SI, Zorn AM, Shroyer NF, Wells JM (2011) Directed differentiation of human pluripotent stem cells into intestinal tissue in vitro. Nature 470: 105–9

Staudinger JL, Goodwin B, Jones SA, Hawkins-Brown D, MacKenzie KI, LaTour A, Liu Y, Klaassen CD, Brown KK, Reinhard J, Willson TM, Koller BH, Kliewer SA (2001) The nuclear receptor PXR is a lithocholic acid sensor that protects against liver toxicity. Proc Natl Acad Sci U S A 98: 3369–74

Tang C, Kakuta S, Shimizu K, Kadoki M, Kamiya T, Shimazu T, Kubo S, Saijo S, Ishigame H, Nakae S, Iwakura Y (2018) Suppression of IL-17F, but not of IL-17A, provides protection against colitis by inducing Treg cells through modification of the intestinal microbiota. Nat Immunol 19: 755–765

Toporova L, Macejova D, Brtko J (2016) Radioligand binding assay for accurate determination of nuclear retinoid X receptors: A case of triorganotin endocrine disrupting ligands. Toxicology letters 254: 32–6

Velazquez-Campoy A, Kiso Y, Freire E (2001) The binding energetics of first- and second-generation HIV-1 protease inhibitors: implications for drug design. Archives of biochemistry and biophysics 390: 169–75

Velazquez-Campoy A, Todd MJ, Freire E (2000) HIV-1 protease inhibitors: enthalpic versus entropic optimization of the binding affinity. Biochemistry 39: 2201–7

Venkatesh M, Mukherjee S, Wang H, Li H, Sun K, Benechet AP, Qiu Z, Maher L, Redinbo MR, Phillips RS, Fleet JC, Kortagere S, Mukherjee P, Fasano A, Le Ven J, Nicholson JK, Dumas ME, Khanna KM, Mani S (2014) Symbiotic bacterial metabolites regulate gastrointestinal barrier function via the xenobiotic sensor PXR and Toll-like receptor 4. Immunity 41: 296–310

Vetizou M, Pitt JM, Daillere R, Lepage P, Waldschmitt N, Flament C, Rusakiewicz S, Routy B, Roberti MP, Duong CP, Poirier-Colame V, Roux A, Becharef S, Formenti S, Golden E, Cording S, Eberl G, Schlitzer A, Ginhoux F, Mani S et al. (2015) Anticancer immunotherapy by CTLA-4 blockade relies on the gut microbiota. Science 350: 1079–84

Wallace BD, Betts L, Talmage G, Pollet RM, Holman NS, Redinbo MR (2013) Structural and functional analysis of the human nuclear xenobiotic receptor PXR in complex with RXRalpha. Journal of molecular biology 425: 2561–77

Wang H, Venkatesh M, Li H, Goetz R, Mukherjee S, Biswas A, Zhu L, Kaubisch A, Wang L, Pullman J, Whitney K, Kuro-o M, Roig AI, Shay JW, Mohammadi M, Mani S (2011) Pregnane X receptor activation induces FGF19-dependent tumor aggressiveness in humans and mice. The Journal of clinical investigation 121: 3220–32

Williamson B, Lorbeer M, Mitchell MD, Brayman TG, Riley RJ (2016) Evaluation of a novel PXR-knockout in HepaRG cells. Pharmacology research & perspectives 4: e00264

Xie W, Barwick JL, Simon CM, Pierce AM, Safe S, Blumberg B, Guzelian PS, Evans RM (2000) Reciprocal activation of xenobiotic response genes by nuclear receptors SXR/PXR and CAR. Genes Dev 14: 3014–23

Yan L, Wang Y, Liu J, Nie Y, Zhong XB, Kan Q, Zhang L (2017) Alterations of Histone Modifications Contribute to Pregnane X Receptor-Mediated Induction of CYP3A4 by Rifampicin. Molecular pharmacology 92: 113–123

Zhu Y, Sun N, Yu M, Guo H, Xie Q, Wang Y (2019) Discovery of aryl-substituted indole and indoline derivatives as RORgammat agonists. Eur J Med Chem 182: 111589

