## Supplementary Information for "Targeting the Pregnane X Receptor Using Microbial Metabolite Mimicry"

[1] From the Department of Cell Biology and Genetics, Palacký University, Olomouc 78371, Czech Republic; Departments of [2] Biochemistry, [3] Medicine, Genetics and Molecular Pharmacology, and [4] Pathology of The Albert Einstein College of Medicine, Bronx, NY 10461, USA; [5] Cincinnati Children's Hospital Medical Center, Cincinnati, OH 45229; [6] Department of Veterinary and Biomedical Sciences, Penn State College of Agricultural Sciences, University Park, PA 16802, USA; [7] Institute of Experimental Endocrinology, BMC, Slovak Academy of Sciences, Dúbravská cesta 9, 845 05 Bratislava, Slovac Republic; [8] The Department of Biology, Johns Hopkins University, Baltimore, MD 21218, USA; [9] Department of Chemistry, University of North Carolina, Chapel Hill, NC 27599; [10] The Department of Pathology, New York University School of Medicine, New York, NY 10016; [11] The Department of Microbiology and Immunology, Drexel University College of Medicine, Philadelphia, PA 19129 USA; [12] Department of Chemistry, City University of New York-Hunter College, New York NY 10065; [13] Molecular Biophysics Unit, Indian Institute of Science, Bangalore 560012, India; [14] Department of Pharmacy – Pharmaceutical Sciences, University of Bari Aldo Moro, Bari 70125, Italy; [15] Department of Environmental and Occupational Health Sciences, University of Washington, Seattle, WA 98105; [16] Center for Metabolic and Vascular Biology, College of Health Solutions, Arizona State University, Scottsdale, AZ 85259; [17] The Department of Biological and Environmental Engineering, Cornell University, Ithaca, NY 14853; [18] Department of Mathematics, City College of The City University of New York, NY, NY 10031; [19] Department of Pediatrics and Immunology, University of Connecticut, Farmington, CT 06030; [20] Department of Pharmaceutical Sciences, University of Connecticut, Storrs, Connecticut 06269-3092; [21] Department of Physiology and Pharmacology, University of Calgary, Calgary, AB Canada T2N 4N1; [22] Center for Gastrointestinal Biology and Disease, Department of Medicine, Division of Gastroenterology and Hepatology, University of North Carolina at Chapel Hill, Chapel Hill, NC 27599, USA

[$] Present Address: St. Edmund's College, Shillong, Old Jowai Road, Shillong, Meghalaya 793003, India

**KEYWORDS: Pregnane X Receptor, microbial metabolite, mimics, drugs, therapy, xenobiotic receptors, tryptophan, indole, indole propionate**

Table of Contents

| | |
|---|---|
| Synthesis of the Indole Metabolite Mimics FKK1-FKK9 | Page 3-22 |
| NMR Traces | Page 23-32 |
| Ligand efficiency metric (LEM) analysis for FKK ligand efficiency | Page 33 |
| X-Ray Analysis | Page 34 |
| In silico Experiments | Page 34-36 |
| Immunoblotting | Page 36-37 |
| Kinase Assays | Page 38-41 |
| Labeled[11,12-$^{3}$H(N)] 9-*cis* retinoic acid (9*c*RA) binding competition assay | Page 41 |
| hFXR TR-FRET | Page 41-42 |
| PPARγ reporter assay | Page 42 |
| Appendix Figure S1 | Page 43 |
| Appendix Figure S2 | Page 44 |
| Appendix Figure S3 | Page 45 |
| Appendix Figure S4 | Page 46 |
| Appendix Figure S5 | Page 47 |
| Appendix Figure S6 | Page 48 |
| Extended View Results (Tables) | Page 49 |
| Table EV1 | Page 50 |
| Table EV2 | Page 51 |
| Table EV3 | Page 52 |
| Table EV4 | Page 53 |
| Table EV5A | Page 54 |
| Table EV5B | Page 55 |
| Table EV5C | Page 56 |

#### Appendix

**Synthesis of the Indole Metabolite Mimics FKK1-FKK9**

##### Materials and Methods for the Chemical Syntheses

All chemical reagents and solvents were obtained from commercial sources (*Aldrich*, *Acros*, *Fisher*) and used without further purification unless otherwise noted. Anhydrous solvents (tetrahydrofurane, toluene, dichloromethane, diethyl ether) were obtained using a *Pure Solv* ™ AL-258 solvent purification system. Ethanol was dried over activated 4 Å molecular sieves. Chromatography was performed either on a *Teledyne ISCO CombiFlash* $R_f$ *200i* using disposable silica cartridges (4, 12, and 24 g). Analytical thin layer chromatography (TLC) was performed on aluminum-backed *Silicycle* silica gel plates (250 μm film thickness, indicator F254). Compounds were visualized using a dual wave length (254 and 365 nm) UV lamp, and/or staining with CAM (cerium ammonium molybdate) or $KMnO_4$ stains. NMR spectra were recorded on *Bruker DRX 300* and *DRX 600* spectrometers. $^1$H and $^{13}$C chemical shifts (δ) are reported relative to tetramethyl silane (TMS, 0.00/0.00 ppm) as internal standard or to residual solvent ($CD_3OD$: 3.31/49.00 ppm; $CDCl_3$: 7.26/77.16 ppm; dmso-$d_6$: 2.50/39.52 ppm; acetone-$d_6$: 2.05/29.84 ppm). Mass spectra (ESI-MS) were recorded on a *Shimadzu LCMS 2010EV* (direct injection unless otherwise noted). High resolution mass spectra (HRMS) and some full mass spectra (ESI-MS) were recorded on an *Orbitrap Velos* high resolution mass spectrometer (*Thermo Finnigan*).

Compounds samples submitted for screening were ≥90% pure as judged by the respective $^1$H-NMR traces. The samples were vialed and dried in high vacuum to constant weight to eliminate residual solvent.

#### Synthetic Procedures

**Figure EV1**: Overview of the synthetic routes. The screened compounds are highlighted in blue (DMF – *N,N*-dimethyl formamide, THF = tetrahydrofurane, μw = microwave); initial synthetic targets in green, lead compounds in red.

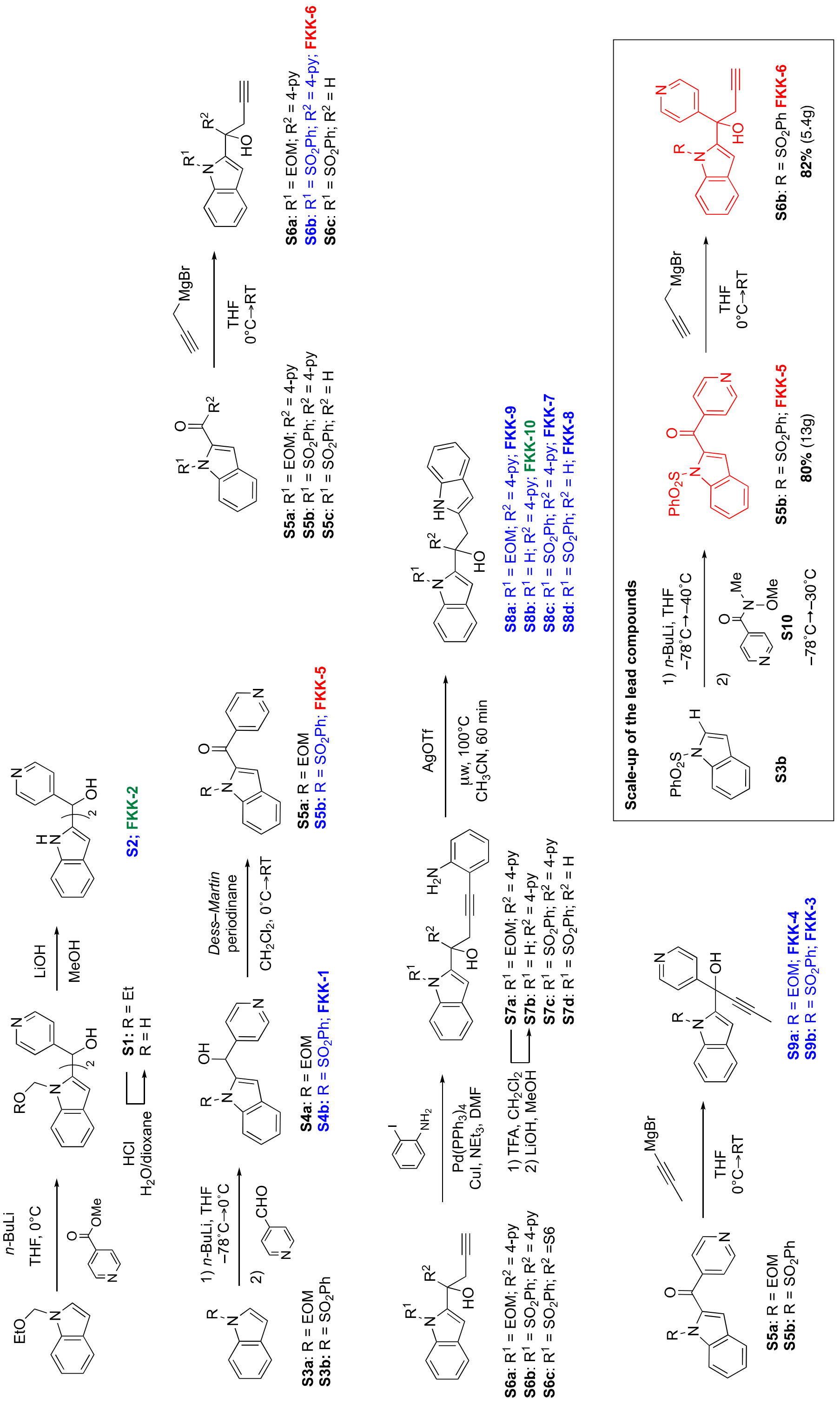

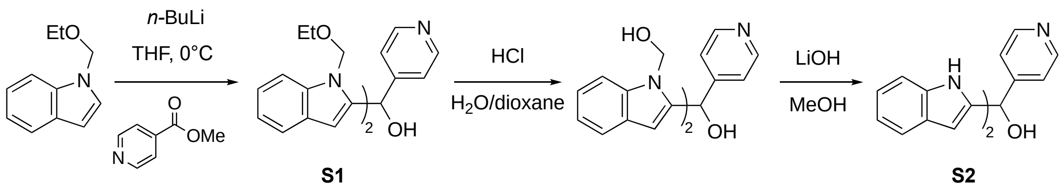

**S1** **S2**

**Scheme 1**: Synthesis of the symmetric bis-indolylcarbinol **S2**: Reacting methyl-4-picolinate with an excess of 2-lithiated 1-(ethoxymethyl)-1*H*-indole affords the intermediate carbinol **S1**. Various literature protocols for the deprotection of both EOM groups were examined, but to no avail. Interestingly, treatment with hydrochloric acid selectively removes the terminal ethyl ether from both EOM-protected indoles. The remaining hydroxymethyl groups can then be cleaved using lithium hydroxide in methanol (THF = tetrahydrofurane).

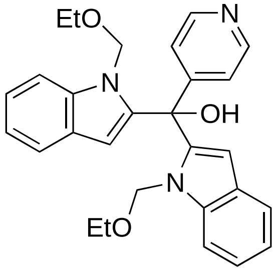

**Synthesis of bis(1-(ethoxymethyl)-1*H*-indol-2-yl)(pyridin-4-yl)methanol (S1):** In a 10mL screw-cap tube with stir bar and septum, 1-(ethoxymethyl)-1*H*-indole (53.7 mg, 0.306 mmol, 2.10 equiv) was dissolved in THF (0.50 mL) and cooled to –78 °C for 10 min. *n*-Butyllithium (0.200 mL, 0.320 mmol, 2.20 equiv) was added drop wise *via* syringe (reaction mixture turned yellow). The mixture was stirred for 1 h, at –78 °C, then warmed to 0 °C (mixture turned red). After stirring at 0 °C for over-all 0.5 h, the mixture was re-cooled to –78 °C. Methyl isonicotinate (20.0 mg, 0.146 mmol) was added neat (immediate color change to lighter red, then dark green), the mixture stirred at –78 °C for 5 min, then warmed to RT (color change to red after a while). TLC analysis of a reaction aliquot (satd. aq. $NH_4Cl$/EtOAc micro-workup) indicated full conversion after 0.5 h, at RT. Satd. aq. $NH_4Cl$ (1.0 mL) was added, followed by ethyl acetate (3.0 mL). The layers were thoroughly mixed, separated and the aqueous layer was extracted with EtOAc (2 × 2.0 mL). The combined organic layers were dried ($MgSO_4$), filtered and evaporated *in vacuo* to obtain the product as orange oil. Chromatography (silica; 0→15 % EtOAc in hexane) afforded bis(1-(ethoxymethyl)-1*H*-indol-2-yl)(pyridin-4-yl)methanol (**S1**; 11.7 mg, 0.026 mmol, 18 %) as colorless resin that solidifies by time.

**TLC**: $R_f$ 0.47 (hex/EtOAc). **$^1$H-NMR** (600 MHz, $CDCl_3$): δ 8.62 (d, *J* = 4.2 Hz, 2H), 7.50–7.43 (m, 6H), 7.29–7.26 (m, 2H), 7.14 (t, *J* = 7.6 Hz, 2H), 6.35 (s, 1H), 5.90 (s, 2H), 5.66 (d, *J* = 11.1 Hz, 2H), 5.48 (d, *J* = 11.1 Hz, 2H), 3.45–3.35 (m, 4H), 1.08 (t, *J* = 7.0 Hz, 6H). **$^{13}$C-NMR** (151 MHz, $CDCl_3$): δ

152.7, 149.9, 141.2, 139.1, 126.7, 123.3, 122.5, 121.3, 120.9, 110.1, 106.6, 75.0, 73.8, 64.1, 14.9. **ESI-MS** *m/z* (rel int): (pos) 478.2 ($[M+Na]^+$, 38), 456.2 ($[M+H]^+$, 100); (neg) 490.2, ($[M+Cl]^-$, 40), 456.2 ($[M–H]^-$, 100).

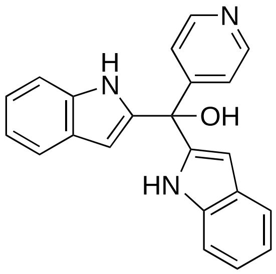

**Synthesis of di(1*H*-indol-2-yl)(pyridin-4-yl)methanol (S2, FKK2):** In a 3 mL screw-cap vial with stir bar, bis(1-(ethoxymethyl)-1*H*-indol-2-yl)(pyridin-4-yl)methanol (**S1**; 14.0 mg, 32.0 μmol) were dissolved in a mixture of dioxane (0.64 mL) and 1 N HCl (0.320 mL, 0.320 mmol, 10.0 equiv). The mixture was stirred at 50 ˚C. The progress of the reaction was closely followed by TLC analysis of reaction aliquots (satd. aq. $NaHCO_3$/EtOAc micro-workup). Previous experiments had shown that prolonged exposure can lead to decomposition. After 3.5 h, all starting material had been consumed and 2 new spots had formed (identified as mono-deprotected and double-deprotected intermediates by MS analysis). The reaction mixture was poured on satd. aq. $NaHCO_3$ (5.0 mL) and extracted with EtOAc (3 × 7.0 mL). The combined organic layers were dried ($MgSO_4$), filtered and evaporated *in vacuo*. Crude $^1$H-NMR analysis confirmed loss of the ethyl group. The residue from the first reaction was dissolved in methanol (0.64 mL) and lithium hydroxide (3.8 mg, 0.16 mmol) was added at RT and the mixture was stirred at the same temperature. TLC analysis of reaction aliquots (satd. aq. $NH_4Cl$/EtOAc micro-workup) after 2 h indicated complete conversion (*NOTE*: TLC conditions: silica, 5% MeOH in $CH_2Cl_2$, 3 × elution. minimal $R_f$ difference!). Crude MS analysis confirmed the presence of the product mass. The reaction was terminated by addition of satd. aq. $NH_4Cl$ (4.0 mL) and the resulting mixture was extracted with EtOAc (3 × 5 mL). The combined organic layers were dried ($MgSO_4$), filtered, and evaporated *in vacuo*. Chromatography (silica; 0.2→7.0 % MeOH in $CH_2Cl_2$) afforded di(1*H*-indol-2-yl)(pyridin-4-yl)methanol (2.5 mg, 7.4 μmol, 23 %) as pale brown solid (despite extended drying over 3d, the sample contained 8% wt residual EtOAc).

**TLC**: $R_f$ 0.76 (5% MeOH in $CH_2Cl_2$, triple elution). **$^1$H-NMR** (300 MHz, MeOD): $\delta$ 8.51 (d, *J* = 6.2 Hz, 2H), 7.60 (dd, *J* = 4.6, 1.6 Hz, 2H), 7.47 (dt, *J* = 7.8, 1.0 Hz, 2H), 7.34 (dd, *J* = 8.1, 0.9 Hz, 2H), 7.08 (ddd, *J* = 8.1, 7.0, 1.2 Hz, 2H), 6.98 (ddd, *J* = 7.9, 7.0, 1.0 Hz, 2H), 6.17 (d, *J* = 0.8 Hz, 2H), 1.29 (s,

1H). $^{13}$**C-NMR** (75 MHz; MeOD): δ 156.5, 149.7, 142.5, 138.3, 129.0, 123.7, 122.8, 121.4, 120.3, 112.2, 102.8, 75.4. **ESI-MS** *m/z* (rel int): (pos) 340.0 ($[M+H]^+$, 100); (neg) 374 ($[M+Cl]^-$, 20), 338.0 ($[M–H]^-$, 100). **HRMS** (for $C_{22}H_{18}N_3O$): calculated: 340.1444; found: 340.1452.

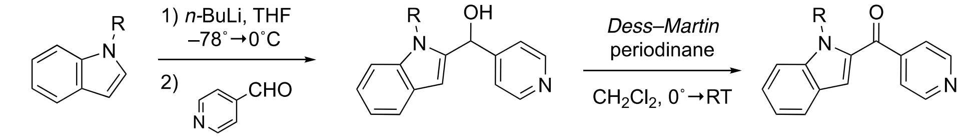

**S3a**: R = EOM
**S3b**: R = $SO_2Ph$

**S4a**: R = EOM
**S4b**: R = $SO_2Ph$

**S5a**: R = EOM
**S5b**: R = $SO_2Ph$

**Scheme 2**: Synthesis of ketones **6a**–**b**: The protected indoles **3a**–**b** were lithiated and and reacted with 4-pyridinecarbaldehyde; the resulting alcohols were oxidized under *Dess–Martin* conditions (THF = tetrahydrofurane).

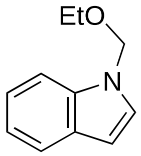

**Synthesis of 1-(ethoxymethyl)-1*H*-indole (3a):** In a flame-dried and argon-flushed 10 mL round-bottom flask, sodium hydride (1.13 g, 28.2 mmol, 1.10 equiv) was suspended in dry DMF (10.0 mL) and cooled to 0 °C. Indole (3.00 g, 25.6 mmol) in dry DMF (10.0 mL) was slowly added over the course of 15 min. The flask used for dissolving the indole was rinsed with additional DMF (5.0 mL). Strong gas formation was observed, indicating deprotonation. After stirring the resulting, more and more viscous slurry for 10 min at 0 °C, the mixture was allowed to warm to RT and stirred at RT for 1.5 h. The mixture was cooled to 0 °C again, then (chloromethoxy)ethane (3.56 mL, 30.7 mmol, 1.20 equiv) was added neat over 10 min in small portions. Most of the material seems to dissolve. The mixture was stirred at 0 °C and followed TLC analysis of reaction aliquots (micro-workup: satd. aq. $NH_4Cl$/EtOAc). After 3 h, complete conversion of the starting material was observed.[1] The milky reaction mixture was poured on 200 mL ice-cold satd. aq. $NaHCO_3$. The mixture was transferred into a separatory funnel, extracted with ethyl acetate (300 mL in total). The organic layer was washed with satd. aq. sodium bicarbonate (100 mL), water (100 mL), satd. aq. $NaHCO_3$ (100 mL), and brine (100 mL). The organic layer was dried ($MgSO_4$), filtered and evaporated *in vacuo* to receive the crude product as pale yellow oil. Chromatography (silica; 0→15 % EtOAc in hexane) afforded 1-(ethoxymethyl)-1*H*-indole (3.17 g, 18.1 mmol, 71 %) as colorless oil.

[1] In other runs, the mixture was stirred overnight, which turned out to be disadvantageous, as considerable decomposition was observed. It is hence recommended to follow the reaction closely and work up as soon as complete conversion is reached.

**TLC**: $R_f$ 0.63 ($CH_2Cl_2$). **$^1$H-NMR** (600 MHz, $CDCl_3$): δ 7.63 (d, *J* = 7.9 Hz, 1H), 7.50 (dd, *J* = 8.2, 0.7 Hz, 1H), 7.24 (ddd, *J* = 8.2, 7.1, 1.0 Hz, 1H), 7.17 (d, *J* = 3.2 Hz, 1H), 7.14 (ddd, *J* = 7.9, 7.1, 0.9 Hz, 1H), 6.53 (dd, *J* = 3.2, 0.7 Hz, 1H), 5.50 (s, 2H), 3.42 (q, *J* = 7.0 Hz, 2H), 1.14 (t, *J* = 7.0 Hz, 3H). **$^{13}$C-NMR** (151 MHz, $CDCl_3$): δ 136.5, 129.2, 128.1, 122.2, 121.1, 120.3, 110.0, 102.6, 76.1, 64.0, 15.0.

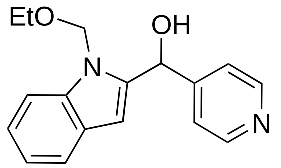

**Synthesis of (1-(thoxymethyl)-1*H*-indol-2-yl)(pyridin-4-yl)methanol (S4a):** In a flame-dried 50 mL centrifuge tube with stir bar and septum, 1-(ethoxymethyl)-1*H*-indole (300 mg, 1.71 mmol) was dissolved in THF (2.85 mL) and cooled to –78 °C. After stirring for 10 min at that temperature, *n*-butyllithium in hexane (1.04 mL, 1.80 mmol, 1.05 equiv) was added drop wise *via* syringe. The mixture was stirred for 10 min, then warmed to 0 °C, upon which it first turned orange, later deeper red. The reaction mixture was stirred at 0 °C for 40 min overall, then re-cooled to –78°C. Isonicotinaldehyde (202 mg, 1.88 mmol, 1.10 equiv) was added at –78°C and the reaction mixture was allowed to slowly warm to RT overnight. TLC analysis of a reaction aliquot (micro-workup: satd. aq. $NH_4Cl$/EtOAc) indicated complete conversion. 10 mL satd. aq. $NH_4Cl$ were added, and the resulting mixture was extracted with EtOAc (1 × 20 mL and 2 × 10 mL). The combined organic layers were dried ($MgSO_4$), filtered and evaporated *in vacuo*. Chromatography (silica; 0.5→5.0 % MeOH in $CH_2Cl_2$) afforded (1-(ethoxymethyl)-1*H*-indol-2-yl)(pyridin-4-yl)methanol (**S4a**; 245 mg, 0.87 mmol, 51 %) as colorless resin.

**TLC**: $R_f$ 0.19 (1:1, hex:EtOAc). **$^1$H-NMR** (600 MHz, $CDCl_3$): δ 8.58 (d, *J* = 5.9 Hz, 2H), 7.56 (d, *J* = 7.9 Hz, 1H), 7.43–7.38 (m, 3H), 7.26 (d, *J* = 5.9 Hz, 1H), 7.14 (t, *J* = 7.4 Hz, 1H), 6.30 (s, 1H), 6.08 (s, 1H), 5.52 (d, *J* = 11.6 Hz, 1H), 5.39 (d, *J* = 11.6 Hz, 1H), 4.47 (s, 1H), 3.44 (q, *J* = 7.0 Hz, 2H), 1.15 (t, *J* = 7.0 Hz, 3H). **$^{13}$C-NMR** (151 MHz, $CDCl_3$): δ 150.7, 149.9, 140.2, 138.6, 127.2, 123.2, 121.6, 121.4, 120.8, 109.3, 105.3, 72.8, 68.3, 64.5, 15.0. **ESI-MS** *m/z* (rel int): (pos) 282.9 ($[M+H]^+$, 100); (neg) 563.3 ($[2M–H]^-$, 50), 317.1 ($[M+Cl]^-$, 45), 281.0 ($[M–H]^-$, 100).

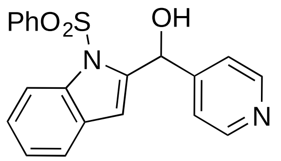

**Synthesis of (1-(phenylsulfonyl)-1*H*-indol-2-yl)(pyridin-4-yl)methanol (S4b, FKK-1):** In a flame-dried and Ar-flushed 60 mL centrifuge tube with stir bar and septum, diisopropylamine (0.130 mL, 0.890 mmol, 1.15 equiv) was dissolved in THF (0.79 mL) and cooled to –78 °C. *n*-Butyllithium (0.530 mL, 0.860 mmol, 1.10 equiv) was slowly added, the mixture stirred first at –78 °C (5 min), then at 0° C (20 min). A solution of 1-(phenylsulfonyl)-1*H*-indole (200 mg, 0.780 mmol) in dry THF (1.57 mL) was the slowly added over 10 min at 0 °C. After stirring for an additional 30 min at 0 °C, the mixture was re-cooled to –78 °C for 10 min. Neat 4-pyridine carbaldehyde (95.0 μL, 1.01 mmol, 1.30 equiv) was added, and the mixture was allowed to warm to RT. TLC analysis of a reaction aliquot (micro-workup: satd. aq. $NH_4Cl$/EtOAc) indicated complete conversion after 2 h 15 min. 10 mL satd. aq. $NH_4Cl$ were added, and the resulting mixture was extracted three times with EtOAc (1 × 20 mL and 2 × 10 mL). The combined organic layers were dried ($MgSO_4$), filtered and evaporated *in vacuo*. Chromatography (silica; 3→15 % MeOH in $CH_2Cl_2$) afforded (1-(phenylsulfonyl)-1*H*-indol-2-yl)(pyridin-4-yl)methanol (**S4b**; 142 mg, 0.39 mmol, 50 %) as off-white solid.

**TLC**: $R_f$ 0.24 (19:1; $CH_2Cl_2$:MeOH). **$^1$H-NMR** (600 MHz, dmso-$d_6$): δ 8.56 (d, *J* = 6.1 Hz, 2H), 8.01 (dd, *J* = 8.4, 0.7 Hz, 1H), 7.91 (dd, *J* = 8.5, 1.1 Hz, 2H), 7.69—7.66 (m, 1H), 7.57—7.53 (m, 3H), 7.40 (d, *J* = 6.1 Hz, 2H), 7.32 (ddd, *J* = 8.4, 7.3, 1.2 Hz, 1H), 7.25—7.22 (m, 1H), 6.62 (s, 1H), 6.49 (d, *J* = 5.8 Hz, 1H), 6.45 (d, *J* = 5.8 Hz, 1H). **$^{13}$C-NMR** (151 MHz; dmso-$d_6$): δ 149.6, 137.4, 136.4, 134.5, 129.6, 128.9, 126.5, 124.9, 123.9, 122.1, 121.4, 114.4, 111.3, 67.2, 39.5. **ESI-MS** *m/z* (rel int): (pos) 365.0 ($[M+H]^+$, 60); (neg) 727.3 ($[2M–H]^-$, 60), 363.0 ($[M–H]^-$, 100). **HRMS** (for $C_{22}H_{15}N_2O_3S_1$): calculated: 365.0954; found: 365.0958.

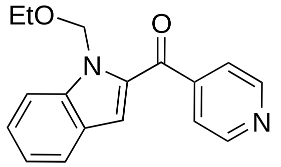

**(1-(Ethoxymethyl)-1*H*-indol-2-yl)(pyridin-4-yl)methanone (S5a)** was synthesized in analogy to **S5b** (*vide infra*) from (1-(ethoxymethyl)-1*H*-indol-2-yl)(pyridin-4-yl)methanol (**S4a**; 38.7 mg, 0.137 mg). Chromatography (silica; 0.2→5.0 % MeOH in $CH_2Cl_2$) afforded (1-(ethoxymethyl)-1*H*-indol-2-yl)(pyridin-4-yl)methanone (**S5a**; 22.1 mg; 0.079 mmol, 58 %) as colorless resin.

**TLC**: $R_f$ 0.52 (95:5 $CH_2Cl_2$:MeOH). **$^1$H-NMR** (600 MHz, $CDCl_3$): δ 8.83 (d, *J* = 6.0 Hz, 2H), 7.70–7.67 (m, 3H), 7.64 (dd, *J* = 8.4, 0.5 Hz, 1H), 7.47 (ddd, *J* = 8.4, 7.1, 1.2 Hz, 1H), 7.23 (ddd, *J* = 7.9, 7.1, 0.8 Hz, 1H), 7.08 (d, *J* = 0.5 Hz, 1H), 6.08 (s, 2H), 3.58 (q, *J* = 7.0 Hz, 2H), 1.16 (t, *J* = 7.0 Hz, 3H).

$^{13}$**C-NMR** (151 MHz, $CDCl_3$): δ 187.0, 150.4, 146.1, 141.0, 133.6, 127.5, 126.3, 123.4, 122.9, 122.1, 118.4, 111.8, 74.0, 64.2, 15.2. **ESI-MS** *m/z* (rel int): (pos) 303.1 ($[M+Na]^+$, 20), 281.1 ($[M+H]^+$, 100).

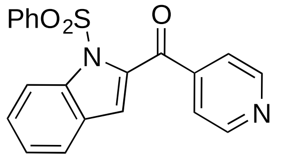

**Synthesis of (1-(phenylsulfonyl)-1*H*-indol-2-yl)(pyridin-4-yl)methanone (S5b, FKK-5)**: In a flame-dried 50 mL round-bottom flask with stir-bar and septum, *Dess–Martin* periodinane (349 mg, 0.823 mmol, 1.50 equiv) was suspended in dichloromethane (5.50 mL) and cooled to 0 °C (wet ice/water bath). A suspension of (1-(phenylsulfonyl)-1*H*-indol-2-yl)(pyridin-4-yl)methanol (200 mg, 0.549 mmol) in dry dichloromethane (22.0 mL) was slowly added over 10 min. The mixture was kept stirring, while the ice bath was allowed to melt slowly. After 35 min, TLC analysis of a reaction aliquot (micro-workup: satd. aq. $Na_2S_2O_3$/EtOAc) indicated complete conversion. The mixture was poured onto 30 mL satd. aq. $Na_2S_2O_3$ and 50 mL EtOAc were added. The resulting mixture was shaken vigorously, and the layers separated. The organic layer was washed with satd. aq. $NaHCO_3$ (30 mL), and brine (30 mL), dried ($MgSO_4$), filtered and evaporated *in vacuo*. Chromatography (silica; 0→40 % EtOAc in hexane) afforded (1-(phenylsulfonyl)-1*H*-indol-2-yl)(pyridin-4-yl)methanone (**S5b**; 169 mg, 0.466 mmol, 85 %) as colorless solid.

**TLC**: $R_f$ 0.37 (1:1, hex:EtOAc). $^{1}$**H-NMR** (600 MHz, $CDCl_3$): δ 8.84 (d, *J* = 6.0 Hz, 2H), 8.15 (dd, *J* = 8.5, 0.7 Hz, 1H), 7.99 (dd, *J* = 8.5, 1.2 Hz, 2H), 7.75 (d, *J* = 6.0 Hz, 2H), 7.59 (t, *J* = 7.6 Hz, 2H), 7.52—7.48 (m, 3H), 7.35—7.32 (m, 1H), 7.04 (d, *J* = 0.7 Hz, 1H). $^{13}$**C-NMR** (151 MHz, $CDCl_3$): δ 186.4, 150.9, 143.9, 138.3, 137.8, 137.2, 134.3, 129.2, 128.6, 128.0, 127.6, 124.9, 123.1, 122.7, 118.9, 115.5. **ESI-MS** *m/z* (rel int): (pos) 425.9 ($[M+Na+MeCN]^+$, 95), 385.9 ($[M+Na]^+$, 100), 363.0 ($[M+H]^+$, 70); (neg) 361.0 ($[M–H]^-$, 100). **HRMS** (for $C_{20}H_{15}N_2O_3S_1$): calculated: 363.0798; found: 363.0799.

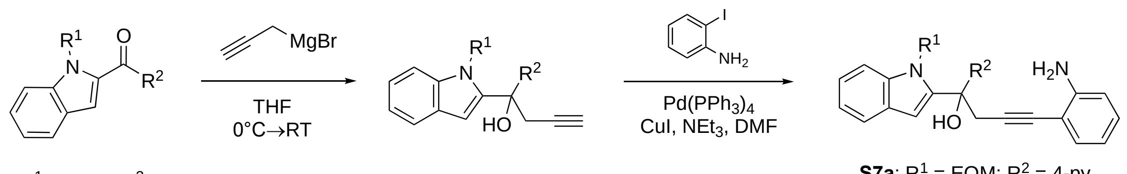

**S5a**: $R^1$ = EOM; $R^2$ = 4-py
**S5b**: $R^1$ = $SO_2Ph$; $R^2$ = 4-py
**S5c**: $R^1$ = $SO_2Ph$; $R^2$ = H

**S6a**: $R^1$ = EOM; $R^2$ = 4-py
**S6b**: $R^1$ = $SO_2Ph$; $R^2$ = 4-py
**S6c**: $R^1$ = $SO_2Ph$; $R^2$ = H

**S7a**: $R^1$ = EOM; $R^2$ = 4-py
**S7c**: $R^1$ = $SO_2Ph$; $R^2$ = 4-py
**S7d**: $R^1$ = $SO_2Ph$; $R^2$ = H

**Scheme 3**: Synthesis of cyclization precursors **7a–d**: Prop-2-yn-1-ylmagnesium bromide was added to the ketones **5a–b** or to 1-(phenylsulfonyl)-1*H*-indole-2-carbaldehyde (**5c**), respectively, to give the corresponding alcohols **6a–c**; *Sonogashira* couplings with 2-iodoaniline furnished the 2-alkynyl anilines **7Xa**,**c** and **d**. For **S7b**, see below. (THF = tetrahydrofurane; DMF = *N,N*-diemthyl formamide; TFA = trifluoroacetic acid).

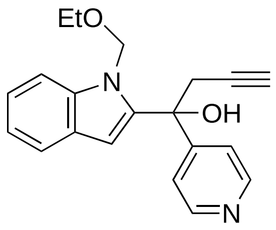

**Synthesis of 1-(1-(ethoxymethyl)-1*H*-indol-2-yl)-1-(pyridin-4-yl)but-3-yn-1-ol (S6a)**: In a dry and argon-flushed 60 mL centrifuge tube, equipped with stir bar and rubber septum, (1-(ethoxymethyl)-1*H*-indol-2-yl)(pyridin-4-yl)methanone (**S5a**; 95.0 mg, 0.339 mmol) was dissolved in dry THF (3.2 mL) and cooled to 0 °C (ice bath) for ca. 10 min. Freshly prepared and titrated prop-2-yn-1-ylmagnesium bromide (0.96M; 1.06 mL, 1.02 mmol, 3.0 equiv)[2] was added drop-wise *via* syringe. TLC analysis of a reaction aliquot (micro-workup: satd. aq. $NH_4Cl$/EtOAc) indicated full conversion after 1.5 h (*NOTE:* can be stirred over night without any problem). The reaction was terminated by the addition of satd. aq. $NH_4Cl$ (1.0 mL). The resulting mixture was warmed to room temperature and extracted with EtOAc (3 × 3.0 mL). The combined organic layers were dried ($MgSO_4$), filtered and evaporated *in vacuo*. Chromatography (silica; 20→55 % EtOAc in hexane) afforded 1-(1-(ethoxymethyl)-1*H*-indol-2-yl)-1-(pyridin-4-yl)but-3-yn-1-ol (**S6a**; 79.0 mg, 0.247 mmol, 73 %) as colorless solid.

**TLC**: $R_f$ 0.08 (4:1, hex:EtOAc, double elution). **$^1$H-NMR** (600 MHz, $CDCl_3$): δ 8.57 (d, *J* = 6.1 Hz, 2H), 7.68 (d, *J* = 7.9 Hz, 1H), 7.37—7.34 (m, 3H), 7.27 (t, *J* = 7.7 Hz, 1H), 7.20—7.18 (m, 1H), 6.95 (s, 1H), 5.28 (d, *J* = 11.5 Hz, 1H), 5.07 (s, 1H), 4.84 (d, *J* = 11.5 Hz, 1H), 3.38—3.29 (m, 2H), 3.21 (dd, *J* = 16.6, 2.6 Hz, 1H), 3.10 (dd, *J* = 16.5, 2.6 Hz, 1H), 2.09 (t, *J* = 2.6 Hz, 1H), 1.10 (d, *J* = 14.1 Hz, 3H). **$^{13}$C-NMR** (151 MHz, $CDCl_3$): δ 153.2, 150.0, 140.5, 138.7, 126.8, 123.5, 121.6, 121.2, 121.0, 109.3, 104.9, 79.3, 73.6, 72.9, 72.8, 64.4, 34.2, 14.8. **ESI-MS** *m/z* (rel int): (pos) 321.1 ($[M+H]^+$, 100).

[2] Prepared according to: Snyder, B. B.; Lin, H. Y.; *Org. Lett.*, **2011**, *13*, 1234–1237; the reagent was used directly in the next step (brief storage at 0 ˚C) and was titrated prior to use according to Paquette's protocol: *Lin, H.-S.; Paquette, L. A.; Synth. Commun.* ***1994****, 24, 2503–2506.*

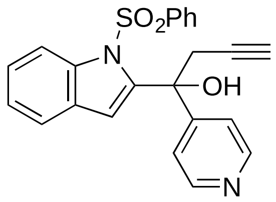

**1-(1-(Phenylsulfonyl)-1*H*-indol-2-yl)-1-(pyridin-4-yl)but-3-yn-1-ol (S6b; FKK-6)** was synthesized in analogy to **S6a** from (1-(phenylsulfonyl)-1*H*-indol-2-yl)(pyridin-4-yl)methanone (**S5b**; 145 mg, 0.400 mmol) and was obtained (149 mg, 0.370 mmol, 92 %) as colorless resin (*NOTE:* no chromatography required).

**TLC**: $R_f$ 0.28 (1:1, hex:EtOAc, double elution). $^{1}$**H-NMR** (600 MHz, $CDCl_3$): δ 8.46 (d, *J* = 5.6 Hz, 2H), 8.04 (d, *J* = 8.3 Hz, 1H), 7.61—7.59 (m, 1H), 7.47 (t, *J* = 7.5 Hz, 1H), 7.37—7.27 (m, 5H), 7.21 (d, *J* = 5.9 Hz, 2H), 7.17 (s, 1H), 5.61 (s, 1H), 3.16—3.09 (m, 2H), 2.08 (t, *J* = 2.6 Hz, 1H). $^{13}$**C-NMR** (151 MHz, $CDCl_3$): δ 152.9, 149.7, 142.1, 138.4, 137.9, 134.0, 129.3, 128.3, 126.2, 126.1, 124.5, 121.8, 121.0, 115.3, 114.1, 79.3, 75.0, 72.8, 35.5. **ESI-MS** *m/z* (rel int): (pos) 425.9 ([M+Na+MeCN]$^{+}$, 95), 403.0 ([M+H]$^{+}$, 100); (neg) 437.1 ([M+Cl]$^{-}$, 15), 401.0 ([M–H]$^{-}$, 50), 156.8 (100). **HRMS** (for $C_{23}H_{19}N_2O_3S_1$): calculated: 403.1111; found: 403.1117.

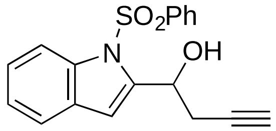

**1-(1-(Phenylsulfonyl)-1*H*-indol-2-yl)but-3-yn-1-ol** (**S6c**) was synthesized in analogy to **S6a** from 1-(phenylsulfonyl)-1*H*-indole-2-carbaldehyde (114 mg, 0.40 mmol) and was obtained (129 mmol, 0.396 mmol, 99 %) as colorless resin (*NOTE:* no chromatography required).

**TLC**: $R_f$ 0.80 (1:1, hex:EtOAc, double elution). $^{1}$**H-NMR** (600 MHz, $CDCl_3$): δ 8.09 (dd, *J* = 8.4, 0.6 Hz, 1H), 7.78 (dd, *J* = 8.6, 1.2 Hz, 2H), 7.52—7.48 (m, 2H), 7.39 (t, *J* = 7.8 Hz, 2H), 7.30 (ddd, *J* = 8.4, 7.2, 1.2 Hz, 1H), 7.25–7.22 (m, 1H), 6.81 (s, 1H), 5.48 (q, *J* = 5.8 Hz, 1H), 3.36 (s, 1H), 3.01 (ddd, *J* = 16.8, 6.2, 2.7 Hz, 1H), 2.90 (ddd, *J* = 16.8, 6.3, 2.7 Hz, 1H), 2.07 (t, *J* = 2.6 Hz, 1H). $^{13}$**C-NMR** (151 MHz, $CDCl_3$): δ 141.9, 138.4, 137.4, 134.1, 129.4, 129.2, 126.5, 125.3, 124.1, 121.5, 114.9, 110.2, 80.2, 71.6, 65.9, 26.8. **ESI-MS** *m/z* (rel int): (pos) 348.1 ([M+Na]$^{+}$, 40), 308.1 ([M +H–$H_2O$]$^{+}$, 100).

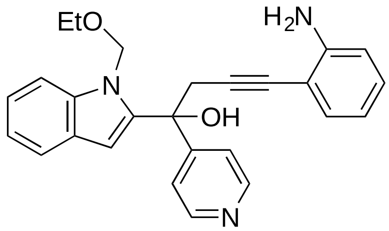

**Synthesis of 4-(2-aminophenyl)-1-(1-(ethoxymethyl)-1*H*-indol-2-yl)-1-(pyridin-4-yl)but-3-yn-1-ol (S7a)**: In a dry and argon-flooded 5 mL vial, equipped with stir bar and septum, 1-(1-(ethoxymethyl)-*1H*-indol-2-yl)-1-(pyridin-4-yl)but-3-yn-1-ol (**S6a**; 33.7 mg, 105 μmol), copper(I) iodide (8.01 mg, 42.0 μmol), and 2-iodoaniline (23.0 mg, 105 μmol) were dissolved in dry DMF (1.50 mL). Triethyl amine (73.0 μl, 53.0 μmol, 5.00 equiv) was added, followed by tetrakis(triphenylphosphino) palladium (24.3 mg, 20.0 μmol, 20.0 mol%). The vial was purged with argon and sealed with a screw cap. The reaction mixture was stirred at room temperature over night. TLC analysis of a reaction aliquot (micro-workup: satd. aq. $NH_4Cl$, 1 dr. $NH_4OH$/EtOAc) indicated full conversion. The mixture was poured onto satd. aq. $NH_4Cl$ containing 2% (v/v) $NH_4OH$ (10 mL), the vial rinsed with little water and EtOAc. The mixture was extracted with EtOAc (2 x 20 mL). The organic layers were combined, washed with water (10 mL) and brine (10 mL), dried ($MgSO_4$), filtered and evaporated *in vacuo*. Repeated chromatography[3] (silica; 20→70 % EtOAc in hexane) afforded 4-(2-aminophenyl)-1-(1-(ethoxymethyl)-1*H*-indol-2-yl)-1-(pyridin-4-yl)but-3-yn-1-ol (**S7a**; 26.9 mg, 65.0 μmol, 62 %) as colorless solid.

**TLC**: $R_f$ 0.25 (1:1, hex:EtOAc). **$^1$H-NMR** (600 MHz, $CDCl_3$): δ 8.57 (d, *J* = 6.0 Hz, 2H), 7.68 (d, *J* = 7.8 Hz, 1H), 7.42 (d, *J* = 6.0 Hz, 2H), 7.36 (d, *J* = 8.3 Hz, 1H), 7.28 (t, *J* = 7.6 Hz, 1H), 7.20—7.18 (m, 1H), 7.16 (dd, *J* = 7.7, 1.3 Hz, 1H), 7.06 (td, *J* = 7.7, 1.1 Hz, 1H), 7.02 (s, 1H), 6.63—6.59 (m, 2H), 5.31 (d, *J* = 11.5 Hz, 1H), 5.10 (s, 1H), 4.90 (d, *J* = 11.5 Hz, 1H), 3.97 (s, 2H), 3.47 (d, *J* = 16.6 Hz, 1H), 3.38 (d, *J* = 16.6 Hz, 1H) 1.08 (t, *J* = 7.0 Hz, 2H). **$^{13}$C-NMR** (151 MHz, $CDCl_3$): δ 153.5, 150.0, 148.4, 140.8, 138.7, 132.1, 129.5, 126.9, 123.5, 121.5, 121.19, 121.04, 117.8, 114.2, 109.5, 107.8, 104.9, 90.1, 81.6, 74.1, 73.0, 64.4, 35.5, 14.8. **ESI-MS** *m/z* (rel int): (pos) 434.1 ($[M+Na]^+$, 60), 412.1 ($[M+H]^+$, 100), 366.0 (50), 348.0 (90); (neg) 446.1 ($[M+Cl]^-$, 65), 410.1 ($[M–H]^-$, 100).

[3] Two columns were required to reduce contamination with triphenylphospine oxide. Compounds were used in the next step if the purity was 95% or better as judged by $^1$H-NMR.

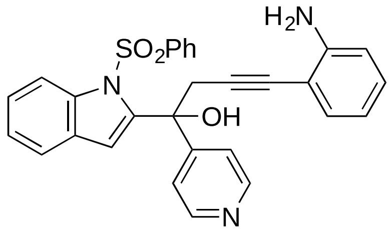

**4-(2-Aminophenyl)-1-(1-(phenylsulfonyl)-1*H*-indol-2-yl)-1-(pyridin-4-yl)but-3-yn-1-ol (S7c)** was synthesized in analogy to **S7a** from 1-(1-(phenylsulfonyl)-1*H*-indol-2-yl)-1-(pyridin-4-yl)but-3-yn-1-ol (**S6b**; 60.0 mg, 0.149 mmol) and 2-iodoaniline (32.7 mg, 0.149 mmol, 1.00 equiv). The reaction was complete after 17 h. Repeated chromatography[3] (silica; 20→90 % EtOAc in hexane) afforded 4-(2-aminophenyl)-1-(1-(phenylsulfonyl)-1*H*-indol-2-yl)-1-(pyridin-4-yl)but-3-yn-1-ol (**S7c**; 32.0 mg, 65.0 μmol, 44 %) as light yellow solid.

**TLC**: $R_f$ 0.15 (1:1, hex:EtOAc). **$^{1}$H-NMR** (600 MHz, $CDCl_3$): δ **$^{1}$H-NMR** (600 MHz, $CDCl_3$): δ 8.47 (d, *J* = 4.1 Hz, 2H), 8.04 (d, *J* = 8.3 Hz, 1H), 7.60 (d, *J* = 7.4 Hz, 1H), 7.47 (t, *J* = 7.5 Hz, 1H), 7.39 (d, *J* = 7.8 Hz, 2H), 7.35 (td, *J* = 7.8, 1.0 Hz, 1H), 7.31—7.24 (m, 2H), 7.13 (dd, *J* = 7.9, 1.3 Hz, 1H), 7.05 (td, *J* = 7.7, 1.3 Hz, 1H), 6.62—6.59 (m, 2H), 5.64 (s, 1H), 3.95 (brs, 2H), 3.44—3.37 (m, 2H). **$^{13}$C-NMR** (151 MHz, $CDCl_3$): δ 153.3, 149.7, 148.3, 142.4, 138.4, 138.0, 134.1, 132.0, 129.5, 129.3, 128.3, 126.2, 126.2, 124.5, 121.8, 121.1, 117.7, 115.3, 114.3, 114.2, 107.8, 90.3, 81.6, 75.4, 36.7. **ESI-MS** *m/z* (rel int): (pos) 516.1 ($[M+Na]^+$, 50), 494.1 ($[M+H]^+$, 100); (neg) 528.1 ($[M+Cl]^-$, 100), 492.1 ($[M–H]^-$, 33).

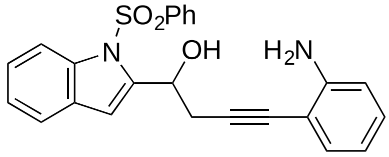

**4-(2-Aminophenyl)-1-(1-(phenylsulfonyl)-1*H*-indol-2-yl)but-3-yn-1-ol (S7d)** was synthesized in analogy to **S7a** from 1-(1-(phenylsulfonyl)-1*H*-indol-2-yl)but-3-yn-1-ol (**S6c**; 65.0 mg, 20.0 μmol) and 2-iodoaniline (43.8 mg, 20.0 μmol, 1.00 equiv). The reaction was complete after 4.5 h. Repeated chromatography[3] (silica; 10→60 % EtOAc in hexane) afforded 4-(2-aminophenyl)-1-(1-(phenylsulfonyl)-1*H*-indol-2-yl)but-3-yn-1-ol (**S7d**; 25.0 mg, 60.0 μmol, 30 %) as colorless solid.

**TLC**: $R_f$ 0.65 (1:1, hex:EtOAc, double elution). **$^{1}$H-NMR** (600 MHz, $CDCl_3$): δ 8.11 (dd, *J* = 8.4, 0.6 Hz, 1H), 7.78 (d, *J* = 7.5 Hz, 2H), 7.50—7.46 (m, 2H), 7.36 (dd, *J* = 8.3, 7.6 Hz, 2H), 7.32 (td, *J* = 7.8, 1.2 Hz, 1H), 7.257.23 (m, 1H), 7.16 (dd, *J* = 7.6, 1.2 Hz, 1H), 7.07 (td, *J* = 7.7, 1.1 Hz, 1H), 6.89 (s, 1H), 6.64—6.60 (m, 2H), 5.56 (t, *J* = 6.2 Hz, 1H), 3.95 (s, 2H), 3.35 (s, 1H), 3.29 (dd, *J* = 16.9, 6.6 Hz, 1H), 3.20 (dd, *J* = 16.9, 5.9 Hz, 1H). **$^{13}$C-NMR** (151 MHz, $CDCl_3$): δ 148.2, 142.3, 138.3, 137.5,

134.1, 132.1, 129.5, 129. 5, 129.3, 126.4, 125.4, 124.2, 121.4, 117.8, 115.0, 114.2, 110.4, 108.0, 90.8, 80.6, 66.4, 28.0. **ESI-MS** *m/z* (rel int): (pos) 458.0 ($[M+H+H_3CCN]^+$, 65), 439.0 ($[M+Na]^+$, 100), 417.0 ($[M+H]^+$, 25), 399.0 (27); (neg) 451.1 ($[M+Cl]^-$, 100), 415.2 ($[M–H]^-$, 32), 156.8 (90).

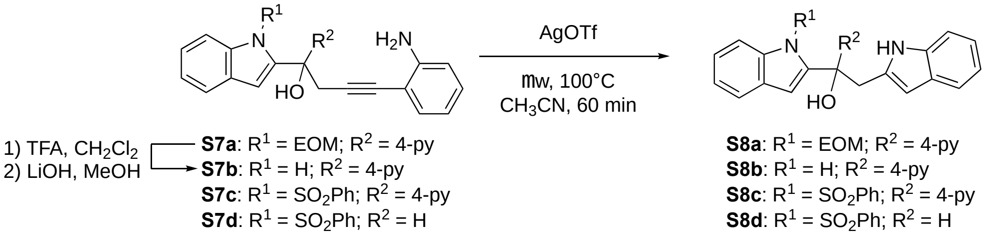

**Scheme 4**: Silver-catalyzed cyclization of precursors **7a**–**d**: Of different protocols examined, the treatment of the cyclization precursors of type **7** with silver(I) triflate in acetonitrile under microwave irradiation proved most efficient. Interestingly, a number of attempts to deprotect **8a** failed. However, a two-step procedure, consisting of treatment with TFA followed by reaction with LiOH (see below), was used successfully to obtain **7b**, which was subsequently cyclized using the same silver-catalyzed protocol as described above (μw = microwave) to obtain **8a**.

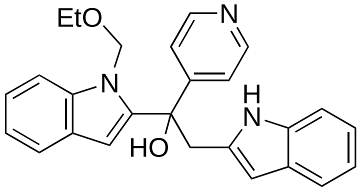

**Synthesis of 1-(1-(ethoxymethyl)-1*H*-indol-2-yl)-2-(1*H*-indol-2-yl)-1-(pyridin-4-yl)ethan-1-ol (S8a; FKK-9):** In a flame-dried microwave vial with septum and stir bar, 4-(2-aminophenyl)-1-(1-(ethoxymethyl)-1*H*-indol-2-yl)-1-(pyridin-4-yl)but-3-yn-1-ol (**S7a**; 15.0 mg, 36.0 μmol) was dissolved in dry acetonitrile (1.0 mL). Silver(I) triflate (3.8 mg, 20 μmol, 0.40 equiv) was added and the vial sealed with a cap under argon. The reaction was heated in the microwave reactor at 120 °C for 1h. TLC analysis of a reaction aliquot indicated complete conversion. The mixture was diluted with EtOAc (ca. 2.0 mL) and filtered through a plug of *Celite* to remove silver salts that had precipitated. The residue was rinsed thoroughly with more EtOAc. The combined organic layers (ca. 5.0 mL) were washed with satd. aq. $NH_4Cl$ (3.0 mL) and brine (3.0 mL), dried ($MgSO_4$), filtered, and evaporated *in vacuo.* Chromatography (silica; 15→60 % EtOAc in hexane) afforded 1-(1-(ethoxymethyl)-1*H*-indol-2-yl)-2-(1*H*-indol-2-yl)-1-(pyridin-4-yl)ethan-1-ol (**S8a**; 6.5 mg, 16 μmol mmol, 43 %) as reddish solid.

**TLC**: $R_f$ 0.61 (1:1, hex:EtOAc, double elution). **$_1$H-NMR** (600 MHz, $CDCl_3$): δ 8.95 (s, 1H), 8.43 (d, *J* = 5.9 Hz, 2H), 7.67 (d, *J* = 7.8 Hz, 1H), 7.45 (d, *J* = 7.8 Hz, 1H), 7.33—7.31 (m, 2H), 7.28—7.26 (m, 1H), 7.20—7.16 (m, 3H), 7.12 (ddd, *J* = 8.1, 7.1, 1.1 Hz, 1H), 7.04–7.01 (m, 2H), 6.04 (d, *J* = 1.0 Hz,

1H), 5.72 (s, 1H), 5.27 (d, *J* = 11.8 Hz, 1H), 4.73 (d, *J* = 11.8 Hz, 1H), 3.93 (d, *J* = 14.4 Hz, 1H), 3.55 (d, *J* = 14.4 Hz, 1H), 3.47–3.37 (m, 2H), 1.18 (t, *J* = 7.0 Hz, 3H). **$^{13}$C-NMR** (151 MHz, $CDCl_3$): δ 154.0, 150.0, 141.8, 138.6, 136.5, 134.0, 128.1, 126.9, 123.5, 121.7, 121.4, 121.1, 120.7, 120.1, 119.4, 110.7, 108.9, 104.9, 102.9, 75.1, 72.6, 64.7, 41.7, 14.9. **ESI-MS** *m/z* (rel int): (pos) 434.1 ($[M+Na]^+$, 100), 412.1 ($[M+H]^+$, 13); (neg) 446.1 ($[M+Cl]^-$, 54), 410.1 ($[M–H]^-$, 100). **HRMS** (for $C_{26}H_{26}N_3O_2$): calculated: 412.2020 ; found: 412.2029.

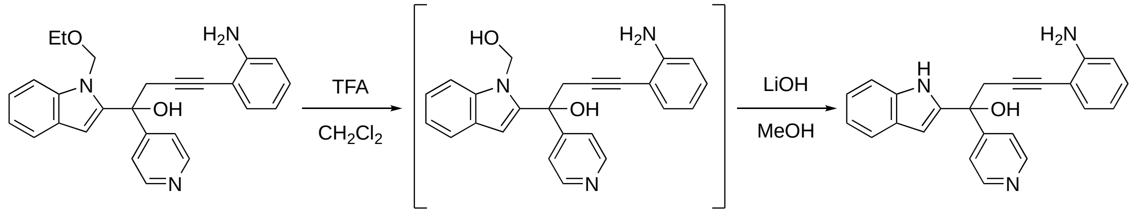

**S7a** **S7b**

**Scheme 5**: Deprotecting the cyclization precursor **S7a**: A two-step deprotection similar to the one developed for **S1** proved useful here. Nevertheless, the overall yield is low, with losses occurring mainly in the second step and probably during purification (TFA = trifluoroacetic acid).

**Synthesis of 4-(2-aminophenyl)-1-(1*H*-indol-2-yl)-1-(pyridin-4-yl)but-3-yn-1-ol (S7b)**: 4-(2-Amino-phenyl)-1-(1-(ethoxymethyl)-1H-indol-2-yl)-1-(pyridin-4-yl)but-3-yn-1-ol (**S7a**; 25.0 mg, 61.0 μmol) was dissolved in $CH_2Cl_2$ (2.53 mL). Trifluoroacetic acid (0.650 mL, 8.51 mmol, 140 equiv) was added at RT and stirred at the same temperature. TLC monitoring (micro-workup, satd. aq. $NaHCO_3$/EtOAc) indicated complete conversion after 5 min. The mixture was poured on satd. aq. $NaHCO_3$ (15 mL). The flask was rinsed with more $CH_2Cl_2$, to reach a final $CH_2Cl_2$ volume of ca. 20 mL. After gas formation ceased, the layers were mixed vigorously, then separated. The aqueous layer was extracted with $CH_2Cl_2$ (4 × 15 mL). The combined organic layers were dried ($MgSO_4$), filtered and evaporated *in vacuo*.

The crude product was dissolved in MeOH (4.40 mL). Lithium hydroxide (7.3 mg, 0.31 mmol, 5.0 equiv) was added and the mixture stirred at RT. TLC analysis after 0.5 h (spotted directly; 1:1 hex:EtOAc, 4 × elution). The mixture was added to satd. aq. $NH_4Cl$ (20 mL), little water (ca. 2.0 mL) was used to rinse the flask. The mixture was extracted with EtOAc (4 × 20mL). The organic layers were dried ($MgSO_4$), filtered, and evaporated *in vacuo*. Chromatography (silica; 20→80 % EtOAc in hexane) afforded 4-(2-aminophenyl)-1-(1*H*-indol-2-yl)-1-(pyridin-4-yl)but-3-yn-1-ol (5.7 mg, 1.5 μmol, 24%) as white resin. The material was obtained in ca. 85 % purity (as estimated by $^1$H-NMR) and used directly in the cyclization reaction.

**TLC**: $R_f$ 0.35 (1:1, hex:EtOAc, double elution). $^{1}$**H-NMR** (600 MHz, CD$_3$CN): δ 9.35 (s, 1H), 8.54 (d, *J* = 6.1 Hz, 2H), 7.55 (dd, *J* = 7.9, 1.0 Hz, 1H), 7.51 (d, *J* = 6.1 Hz, 2H), 7.34—7.33 (m, 1H), 7.12 (ddd, *J* = 8.2, 7.0, 1.2 Hz, 1H), 7.05—7.01 (m, 3H), 6.60 (dd, *J* = 8.2, 0.6 Hz, 1H), 6.57 (dd, *J* = 2.2, 0.9 Hz, 1H), 6.53 (td, *J* = 7.5, 1.1 Hz, 1H), 4.53 (s, 1H), 4.27 (s, 2H), 3.53—3.47 (m, 2H). $^{13}$**C-NMR** (600 MHz, CD$_3$CN): δ 154.3, 150.6, 150.1, 142.9, 132.5, 130.3, 128.8, 122.9, 121.8, 121.4, 120.6, 117.6, 114.7, 112.1, 107.9, 100.3, 91.5, 81.2, 75.0, 61.0, 34.4. **ESI-MS** *m/z* (rel int): (pos) 353.9 ([M+H]$^+$, 100); (neg) 352.0 ([M–H]$^-$, 100).

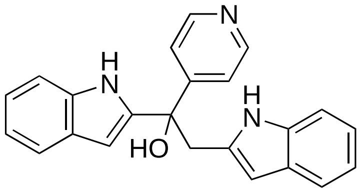

**1,2-Di(1*H*-indol-2-yl)-1-(pyridin-4-yl)ethan-1-ol (S8b, FKK10)** was synthesized in analogy from 4-(2-aminophenyl)-1-(1*H*-indol-2-yl)-1-(pyridin-4-yl)but-3-yn-1-ol (**S7b**; 5.7 mg, 16 μmol). Chromatography (silica; 10→50 % EtOAc in hexane) afforded the desired product (3.5 mg, 10 μmol, 61 %) as off-white solid.

**TLC**: $R_f$ 0.67 (1:1, hex:EtOAc, triple elution). $^{1}$**H-NMR** (600 MHz, DMF-$d_7$): δ 10.99 (brs, 1H), 10.56 (brs, 1H), 8.43 (dd, *J* = 4.6, 1.5 Hz, 2H), 7.55—7.53 (m, 3H), 7.38—7.37 (m, 1H), 7.34 (dd, *J* = 10.7, 8.1 Hz, 2H), 7.06 (t, *J* = 7.5 Hz, 1H), 6.99—6.94 (m, 2H), 6.87 (t, *J* = 7.4 Hz, 1H), 6.58 (d, *J* = 1.6 Hz, 1H), 6.44 (s, 1H), 6.08 (s, 1H), 3.98–3.93 (m, 2H). $^{13}$**C-NMR** (151 MHz, DMF-$d_7$): δ 155.1, 149.9, 145.0, 137.5, 137.0, 135.7, 129.1, 128.7, 121.9, 121.8, 120.8, 120.7, 119.8, 119.6, 119.1, 112.0, 111.6, 102.2, 99.4, 75.1, 40.5. **ESI-MS** *m/z* (rel int): (neg) 220 (20), 388.0 ([M+Cl]$^-$, 100), 352.0 ([M–H]$^-$, 50). **HRMS** (for $C_{23}H_{20}N_3O$): calculated: 354.1601; found: 354.1620.

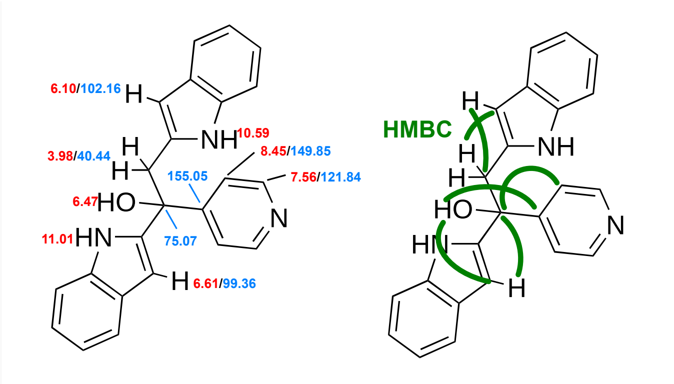

**2D NMR analysis of FKK-10**

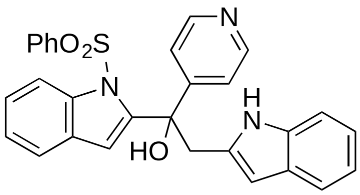

**2-(1*H*-Indol-2-yl)-1-(1-(phenylsulfonyl)-1*H*-indol-2-yl)-1-(pyridin-4-yl)ethan-1-ol (S8c; FKK-7)** was synthesized in analogy from 4-(2-aminophenyl)-1-(1*H*-indol-2-yl)-1-(pyridin-4-yl)but-3-yn-1-ol (**S8c**; 11.0 mg, 22.0 μmol). Chromatography (silica; 15→60 % EtOAc in hexane) afforded the desired product (7.0 mg, 14 μmol, 64 %) as off-white solid.

**TLC**: $R_f$ 0.32 (1:1, hex:EtOAc). $^1$**H-NMR** (600 MHz, $CDCl_3$): δ 9.13 (s, 1H), 8.33 (d, *J* = 4.3 Hz, 2H), 8.06 (d, *J* = 8.4 Hz, 1H), 7.54 (d, *J* = 7.8 Hz, 1H), 7.47—7.45 (m, 2H), 7.35—–7.31 (m, 2H), 7.30—7.27 (m, 1H), 7.23—7.16 (m, 5H), 7.14—7.11 (m, 1H), 7.08—7.02 (m, 3H), 6.06 (s, 1H), 6.00 (s, 1H), 3.80 (d, *J* = 14.7 Hz, 1H), 3.47 (d, *J* = 14.8 Hz, 1H). $^{13}$**C-NMR** (151 MHz, $CDCl_3$): δ 153.7, 149.7, 142.6, 138.4, 137.9, 136.4, 134.2, 134.0, 129.3, 128.3, 128.0, 126.1, 126.0, 124.5, 121.9, 121.5, 120.6, 120.1, 119.5, 115.3, 113.8, 110.8, 102.8, 76.4, 43.6. **ESI-MS** *m/z* (rel int): (pos) 516 ($[M+Na]^+$, 27), 494.1 ($[M+H]^+$, 100); (neg) 528.2 ($[M+Cl]^-$, 20), 492.1 ($[M–H]^-$, 15), 156.8 (100). **HRMS** (for $C_{29}H_{24}N_3O_3S_1$): calculated: 494.1533 ; found: 494.1533.

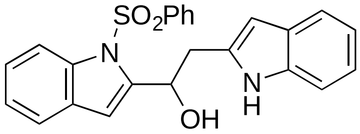

**2-(1*H*-Indol-2-yl)-1-(1-(phenylsulfonyl)-1*H*-indol-2-yl)ethan-1-ol (S8d; FKK-8)** was synthesized in analogy from 4-(2-aminophenyl)-1-(1*H*-indol-2-yl)-1-(pyridin-4-yl)but-3-yn-1-ol (**S7d**; 30.0 mg, 72.0 μmol). Chromatography (silica; 15→60 % EtOAc in hexane) afforded the desired product (12.1 mg, 29.0 μmol, 40 %) as colorless resin.

**TLC**: $R_f$ 0.63 (1:1, hex:EtOAc, double elution). $^1$**H-NMR** (600 MHz, $CDCl_3$): δ 8.80 (s, 1H), 8.18 (d, *J* = 8.4 Hz, 1H), 7.75 (d, *J* = 7.6 Hz, 2H), 7.59 (d, *J* = 7.8 Hz, 1H), 7.58—7.50 (m, 2H), 7.44—7.32 (m, 4H), 7.28 (t, *J* = 7.3 Hz, 1H), 7.20—7.17 (m, 1H), 7.13—7.11 (m, 1H), 6.82 (s, 1H), 6.37 (s, 1H), 5.53 (d, *J* = 6.7 Hz, 1H), 3.58 (dd, *J* = 14.9, 2.8 Hz, 1H), 3.48 (s, 1H), 3.35 (dd, *J* = 14.9, 8.8 Hz, 1H). $^{13}$**C-NMR** (151 MHz, $CDCl_3$): δ 142.9, 138.4, 137.4, 136.4, 136.1, 134.2, 129.5, 129.3, 128.4, 126.3, 125.4, 124.3, 121.6, 121.5, 120.1, 119.8, 115.0, 110.9, 109.8, 101.4, 67.5, 35.3. **ESI-MS** *m/z* (rel int): (pos) 439.0 ($[M+Na]^+$, 100), 417.0 ($[M+H]^+$, 25), 399.0 (27); (neg) 451.1 ($[M+Cl]^-$, 30), 415.1 ($[M–H]^-$, 50), 156.8 (100). **HRMS** (for $C_{24}H_{21}N_2O_3S$): calculated: 417.1267 ; found: 417.1300.

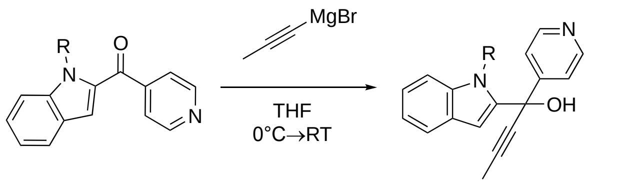

**S5a**: R = EOM
**S5b:** R = $SO_2Ph$

**S9a**: R = EOM; **FKK-4**
**S9b:** R = $SO_2Ph$; **FKK-3**

**Scheme 6**: Synthesis of carbinols **S9a** and **b**: The target alcohols were synthesized by simple addition of commercial prop-1-yn-1-ylmagnesium bromide to the ketons **S5a** and **b** (THF = tetrahydrofurane).

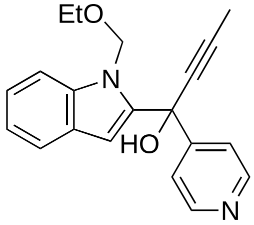

**Synthesis of 1-(1-(ethoxymethyl)-1*H*-indol-2-yl)-1-(pyridin-4-yl)but-2-yn-1-ol (S9a; FKK-4):** (1-(ethoxymethyl)-1*H*-indol-2-yl)(pyridin-4-yl)methanone (**S5a**; 20.0 mg, 71.0 μmol) was dissolved in dry THF (1.43 mL) and cooled to 0 °C for 10 min. Prop-1-yn-1-ylmagnesium bromide (0.50 M; 0.29 mL, 0.14 mmol, 2.0 equiv) was added drop wise *via* syringe. TLC analysis of a reaction aliquot (micro-workup, satd. aq. $NH_4Cl$/EtOAc) indicated full conversion after 1.5 h. The reaction was terminated by addition of 1.0 mL satd. aq. $NH_4Cl$ and the resulting mixture was extracted with EtOAc (3 × 3.0 mL). The combined organic layers were dried ($MgSO_4$), filtered and evaporated. Chromatography (silica; 0→1.5 % MeOH in $CH_2Cl_2$) afforded the desired product (14.8 mg, 46.0 μmol, 65 %) as off-white solid.

**TLC**: $R_f$ 0.78 (5% MeOH in $CH_2Cl_2$). **$^1$H-NMR** (600 MHz, $CDCl_3$): ): δ 8.65 (d, *J* = 6.0 Hz, 2H), 7.61 (d, *J* = 6.0 Hz, 2H), 7.54 (d, *J* = 7.8 Hz, 1H), 7.39 (d, *J* = 8.3 Hz, 1H), 7.26 (d, J = 7.2 Hz. 1H), 7.15—7.12 (m, 1H), 6.32 (s, 1H), 5.94 (d, *J* = 11.5 Hz, 1H), 5.34 (brs, 1H), 5.33 (d, *J* = 11.5 Hz, 1H), 3.48—3.38 (m, 2H), 1.96 (s, 3H), 1.15 (t, *J* = 7.0 Hz, 3H). **$^{13}$C-NMR** (151 MHz, $CDCl_3$): δ 152.3, 149.9, 141.1, 139.1, 126.5, 123.5, 121.6, 121.6, 120.9, 109.3, 106.3, 84.0, 80.2, 73.0, 69.4, 64.5, 14.9, 4.0. **ESI-MS** *m/z* (rel int): (pos) 321.0 ($[M+H]^+$, 100); (pos) 320.9 ($[M+H]^+$, 100). (neg) 319.0 ($[M–H]^-$, 100). **HRMS** (for $C_{20}H_{21}N_2O_2$): calculated: 321.1598; found: 321.1603.

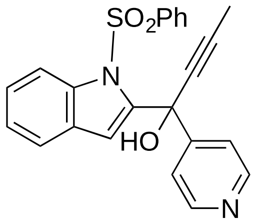

**1-(1-(Phenylsulfonyl)-1*H*-indol-2-yl)-1-(pyridin-4-yl)but-2-yn-1-ol (S9b; FKK-3)** was synthesized in analogy to **S5b** from (1-(phenylsulfonyl)-1*H*-indol-2-yl)(pyridin-4-yl)methanone (20.0 mg, 55.0 μmol). Chromatography (silica; 1.0→2.5 % MeOH in $CH_2Cl_2$) afforded the desired product (11.4 mg, 28.0 μmol, 51 %) as off-white solid.

**TLC**: $R_f$ 0.82 (5% MeOH in $CH_2Cl_2$). **$^1$H-NMR** (600 MHz, $CDCl_3$): δ 8.60 (d, *J* = 5.7 Hz, 2H), 7.95 (d, *J* = 8.4 Hz, 1H), 7.66 (d, *J* = 7.9 Hz, 2H), 7.52–7.50 (m, 3H), 7.46 (d, *J* = 7.6 Hz, 1H), 7.37 (t, *J* = 8.0 Hz, 2H), 7.29 (td, *J* = 7.9, 1.2 Hz, 1H), 7.23 (t, *J* = 7.6 Hz, 1H), 6.70 (s, 1H), 5.67 (s, 1H), 1.90 (s, 3H). **$^{13}$C-NMR** (151 MHz, $CDCl_3$): δ 152.7, 150.0, 143.0, 138.5, 138.3, 134.0, 129.4, 128.0, 126.5, 126.1, 124.4, 121.9, 121.3, 115.7, 115.2, 85.5, 80.0, 70.7, 4.1. **ESI-MS** *m/z* (rel int): (pos) 402.9 ($[M+H]^+$, 100). **HRMS** (for $C_{23}H_{19}N_2O_3S$): calculated: 403.1111; found: 403.1118.

##### Gram-Scale Synthesis of lead compounds FKK5 (S5b) and FKK6 (S6b)

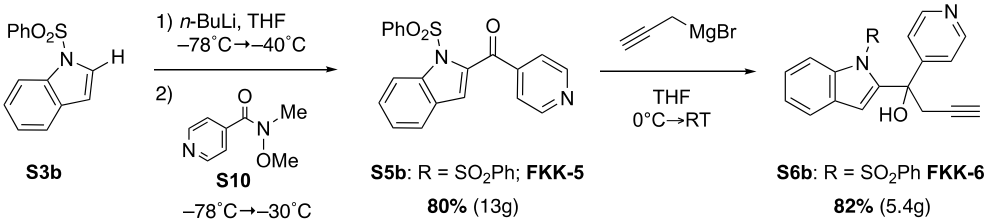

**Scheme 7**: Improved synthesis of **S5b** and **S6b**: For a more efficient synthesis of the two lead compounds, the synthetic route was slightly modified. The addition of a lithium reagent, directly generated from **S3b** and *n*-BuLi, to the Weinreb amide of isonicotinic acid (**S10**) gave easy and scalable access to **S5b** in only one synthetic step, starting exclusively from commercially available starting materials and reagents. The previously utilized addition of a propargylic Grignard reagent to **S5b** proved scalable and delivered **S6b** in excellent yield.

**Large scale synthesis of (1-(phenylsulfonyl)-1*H*-indol-2-yl)(pyridin-4-yl)methanone (S5b; FKK-5)**: In an oven-dried 250mL round-bottom flask with stir bar and septum cap, 1-(phenylsulfonyl)-1H-indole (11.6 g, 45.1 mmol, 1.00 equiv) was dissolved in THF (75.0 mL), and cooled to –78 °C (acetone/dry ice). After stirring for 15 min at that temperature, *n*-butyllithium in hexane (1.64 M; 28.9 mL, 47.4 mmol, 1.05 equiv) was added drop wise *via* syringe (over ca. 20 min). The mixture was stirred for 10 min, then warmed to –40 °C (5 min), then 0 °C (wet ice bath). Precipitate formation started at –78 °C, becoming more pronounced as the mixture warmed up). The slurry was stirred at 0 °C for 40 min, then re-cooled to –78 °C. *N*-Methoxy-*N*-methylisonicotinamide (9.00 g, 54.2 mmol, 1.20 equiv) was lyophilized first from toluene (50.0 mL), then from anhydrous benzene, dissolved in THF (75.0 mL). The resulting solution was added slowly over 15 min *via* cannula at –78 °C. After 5 min at –78 °C, the reaction mixture was warmed to –30 °C. After an additional 2.5 h, at –30 °C, almost complete conversion was observed by TLC analysis of a reaction aliquot (satd. aq. $NH_4Cl$/EtOAc micro-workup) and no discernible change was observed when compared to an earlier sample (t = 1.5 h). The reaction was terminated by addition of $NH_4Cl$ satd. aq. (40.0 mL) at –78 °C. The mixture was warmed to RT with vigorous stirring. Water (40.0 mL) was added, followed by EtOAc (200 mL), the mixture was vigorously shaken, the layers separated. The aqueous layer was extracted with EtOAc (2 × 150 mL), the combined organic layers were dried ($MgSO_4$), filtered and evaporated *in vacuo*. Crude NMR analysis indicated ca. 85 % purity of the crude product. Chromatography (silica; 0→15% EtOAc in $CH_2Cl_2$) afforded the desired product (13.1 g, 36.1 mmol, 80 %) as off-white solid.

**The large scale synthesis of 1-(1-(phenylsulfonyl)-1H-indol-2-yl)-1-(pyridin-4-yl)but-3-yn-1-ol** (**S6b**; **FKK-6**) followed the exact procedures described for the smaller scale synthesis, starting out from (1-(phenylsulfonyl)-1*H*-indol-2-yl)(pyridin-4-yl)methanone (**S5b**; 6.00 g; 16.6 mmol). The crude product was re-crystallized from EtOAc in three batches to obtain the desired product as off-white solid (5.44 g, 13.5 mmol, 82 %).

### Figure EV2. NMR traces of FKK1–10

#### FKK1

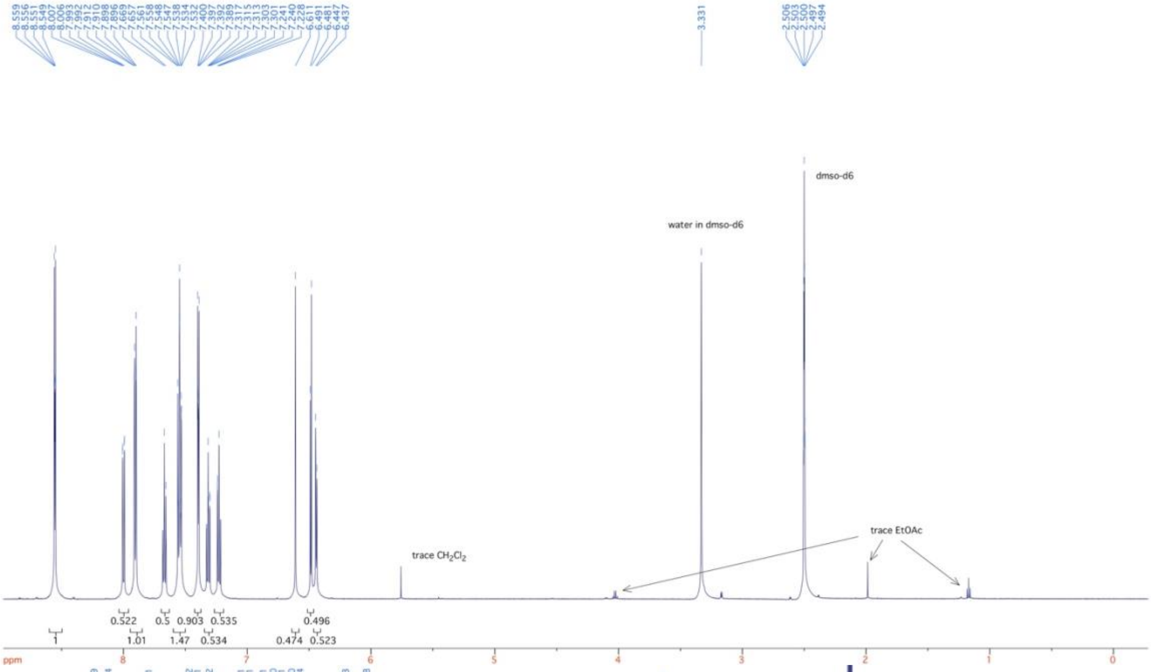

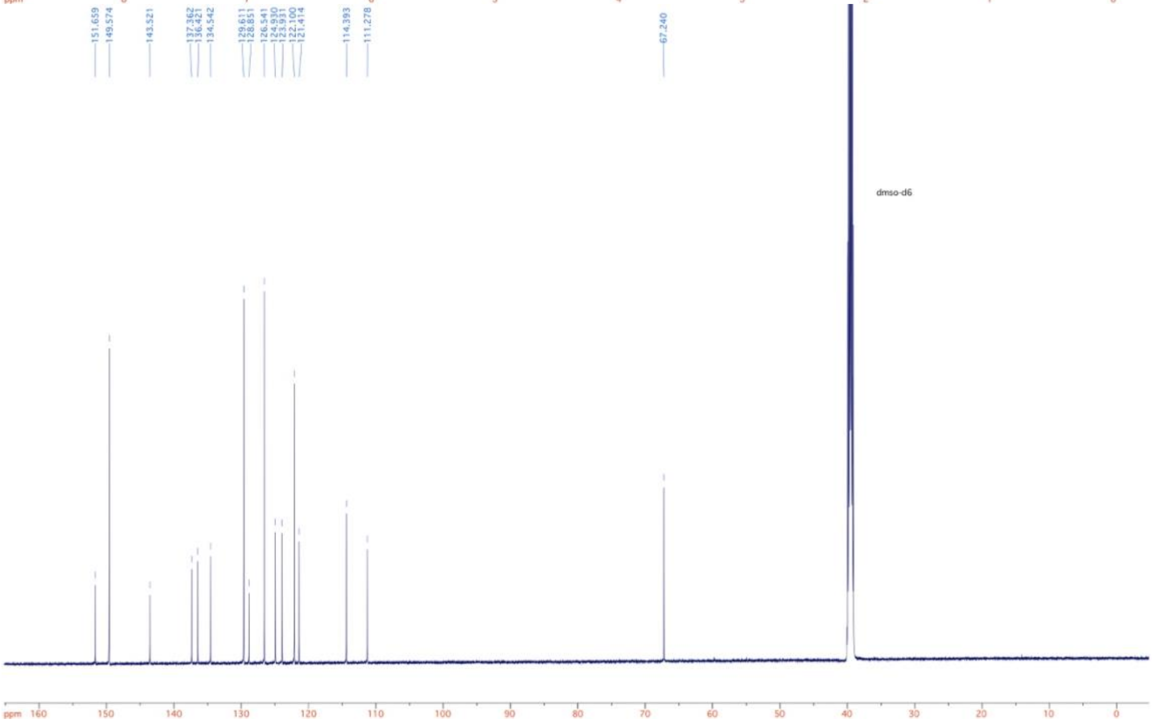

**FKK2**

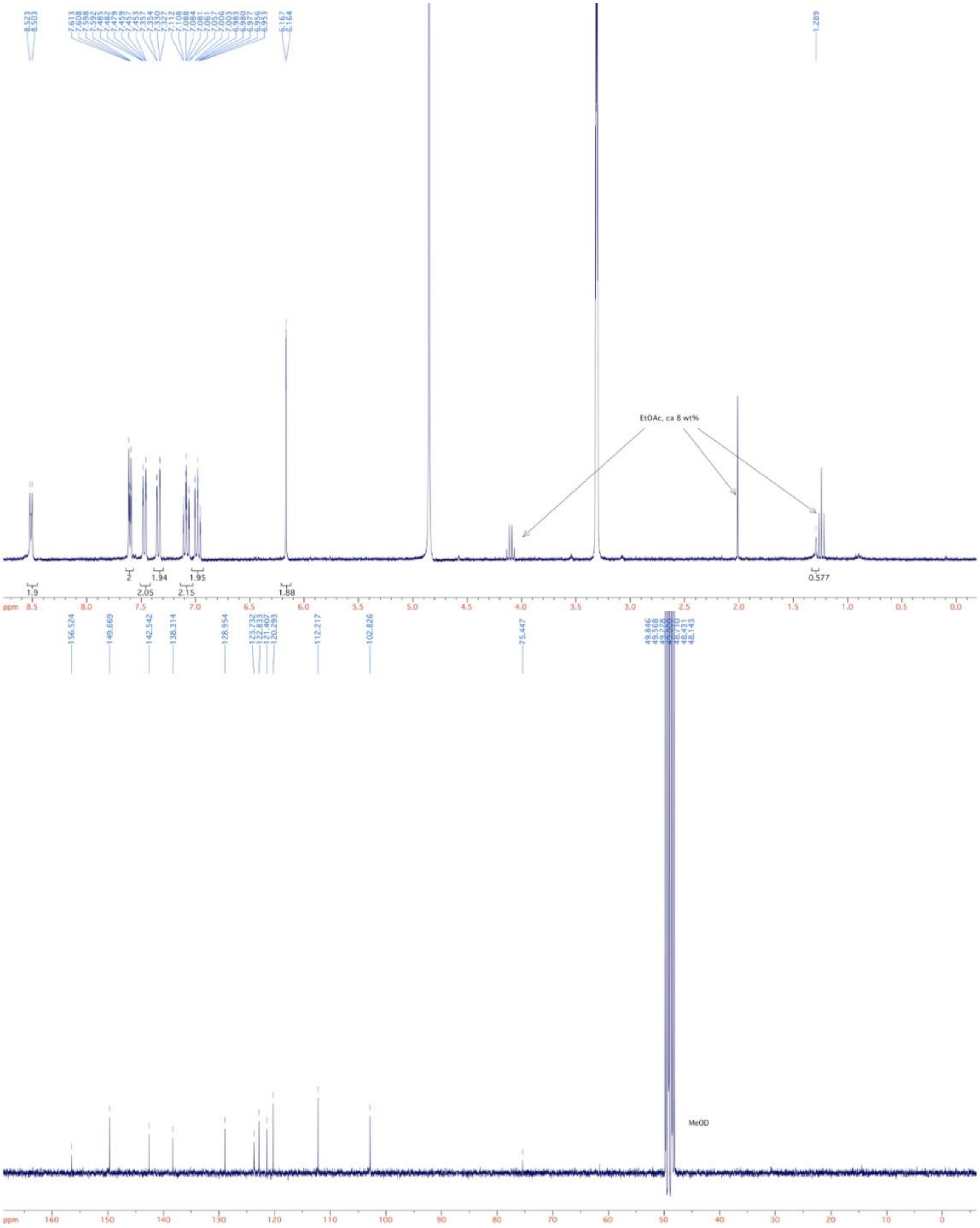

**FKK3**

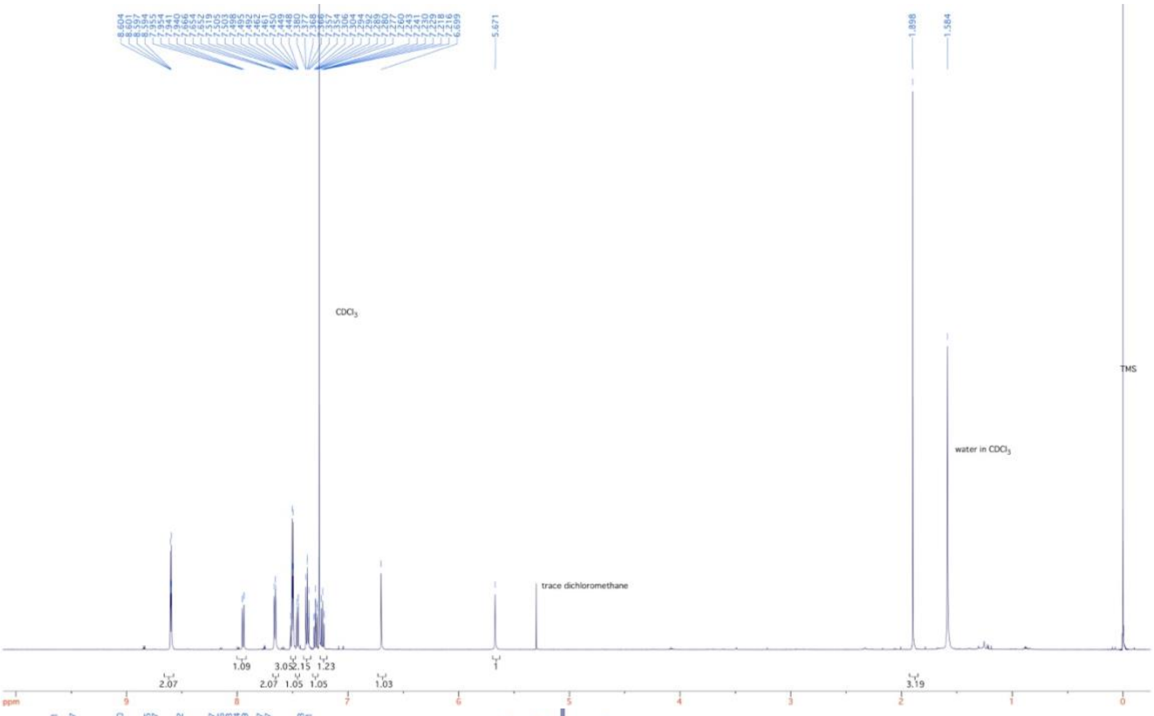
5.671
1.898
1.584
CDCl$_3$
TMS
water in CDCl$_3$
trace dichloromethane
ppm

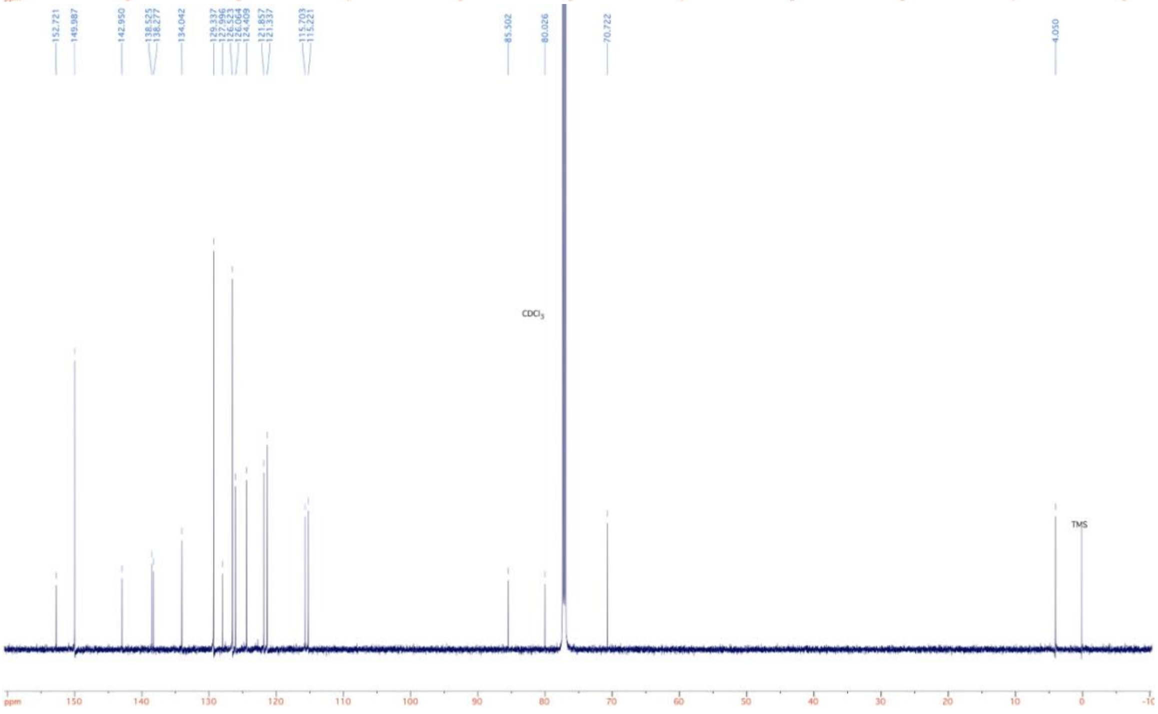
152.721
149.987
142.950
138.525
138.277
134.042
129.337
127.996
126.064
124.409
121.857
121.337
115.703
115.221
85.502
80.026
70.722
4.050
CDCl$_3$
TMS
ppm

**FKK4**

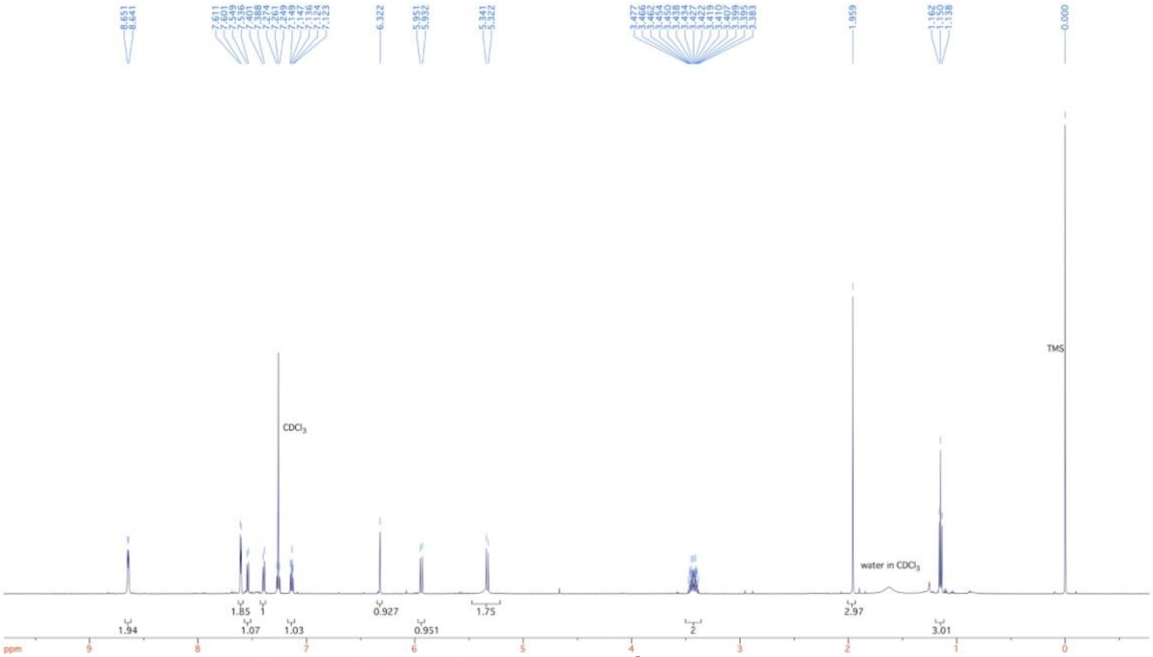

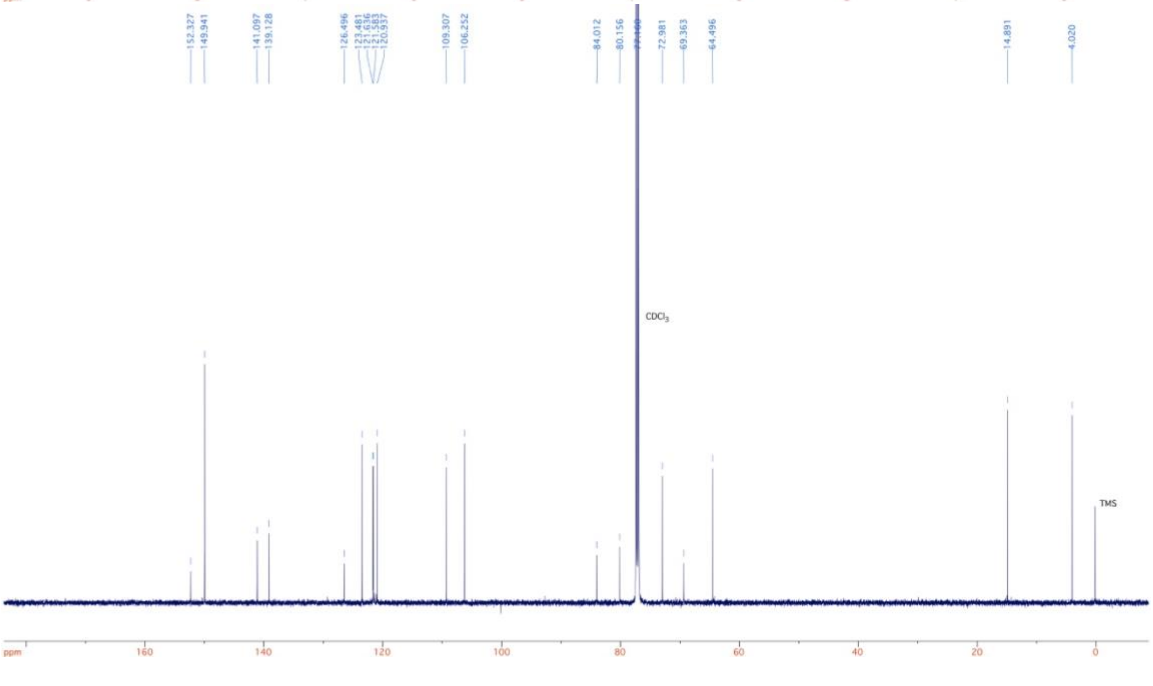

**FKK5**

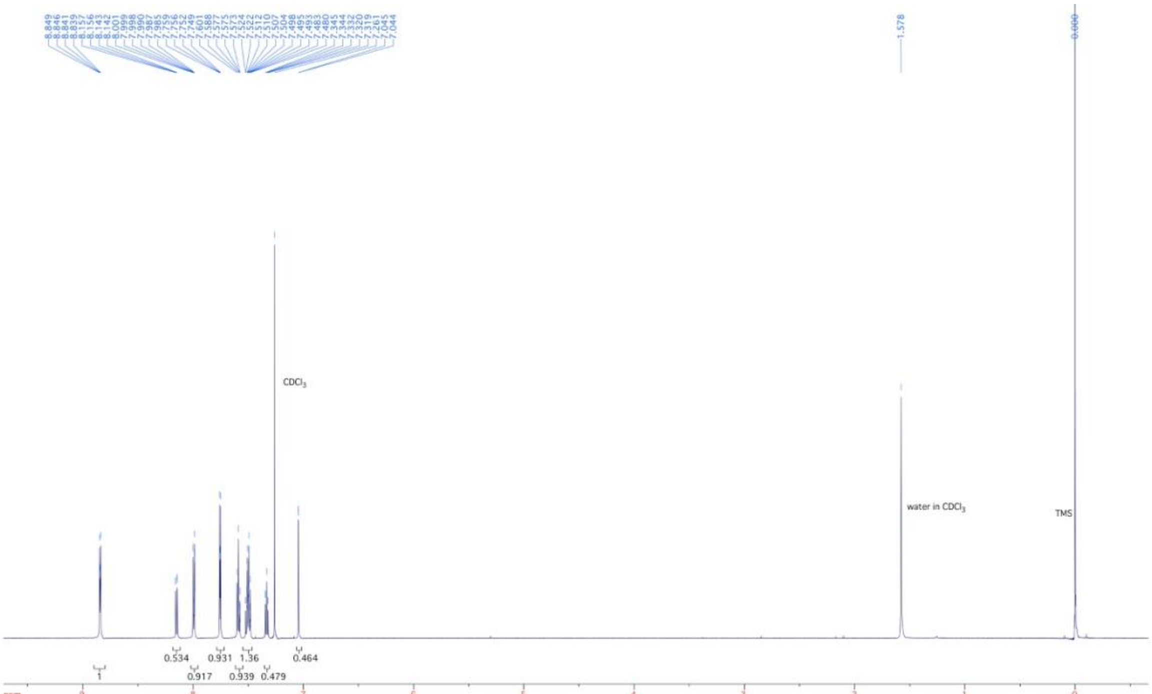

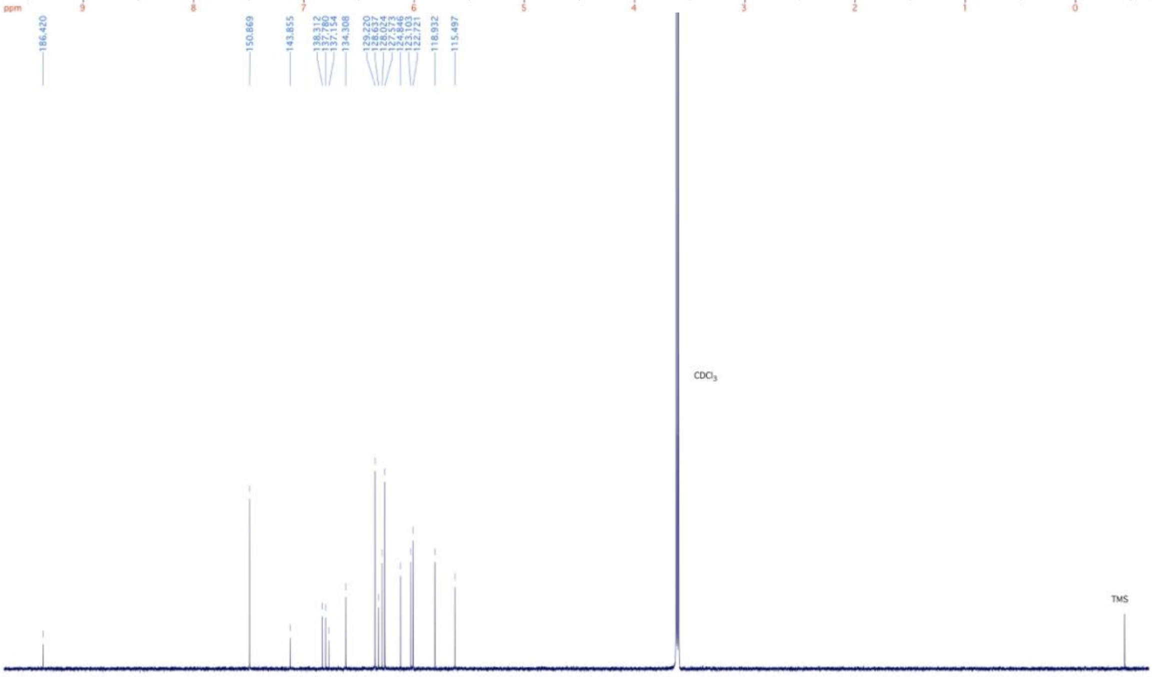

**FKK6**

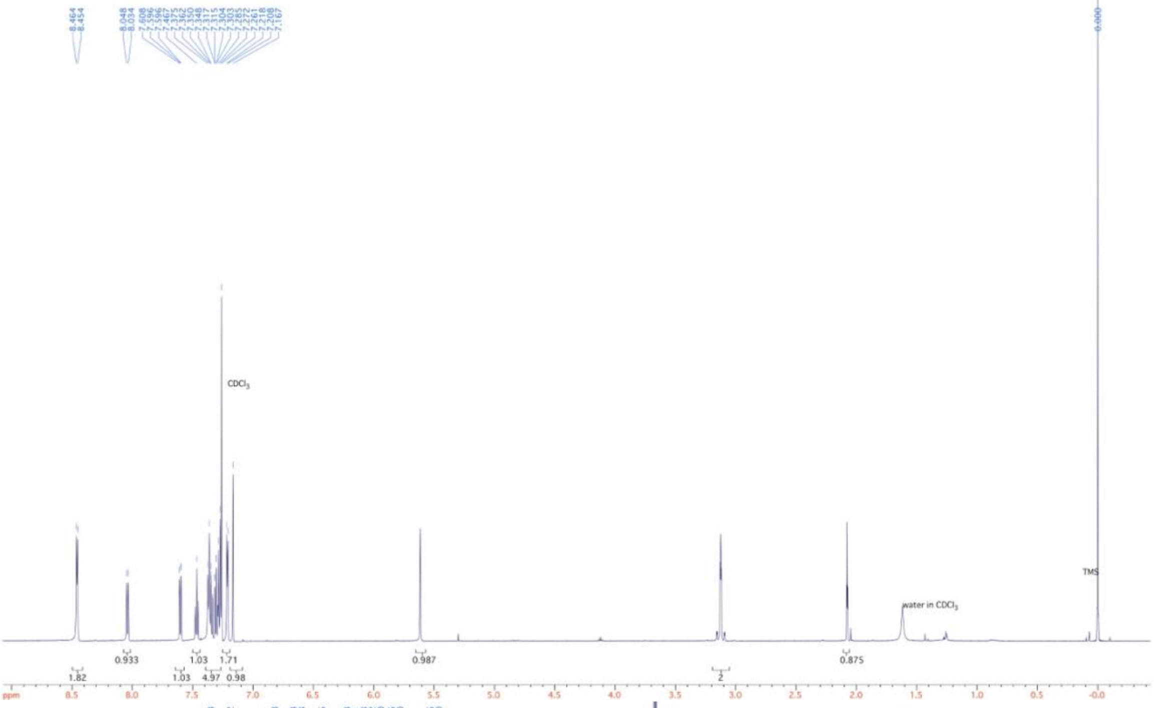

8.464
8.454
8.048
8.034
CDCl$_3$
water in CDCl$_3$
TMS
0.000
1.82
0.933
1.03
1.03
4.97
1.71
0.98
0.987
2
0.875
ppm

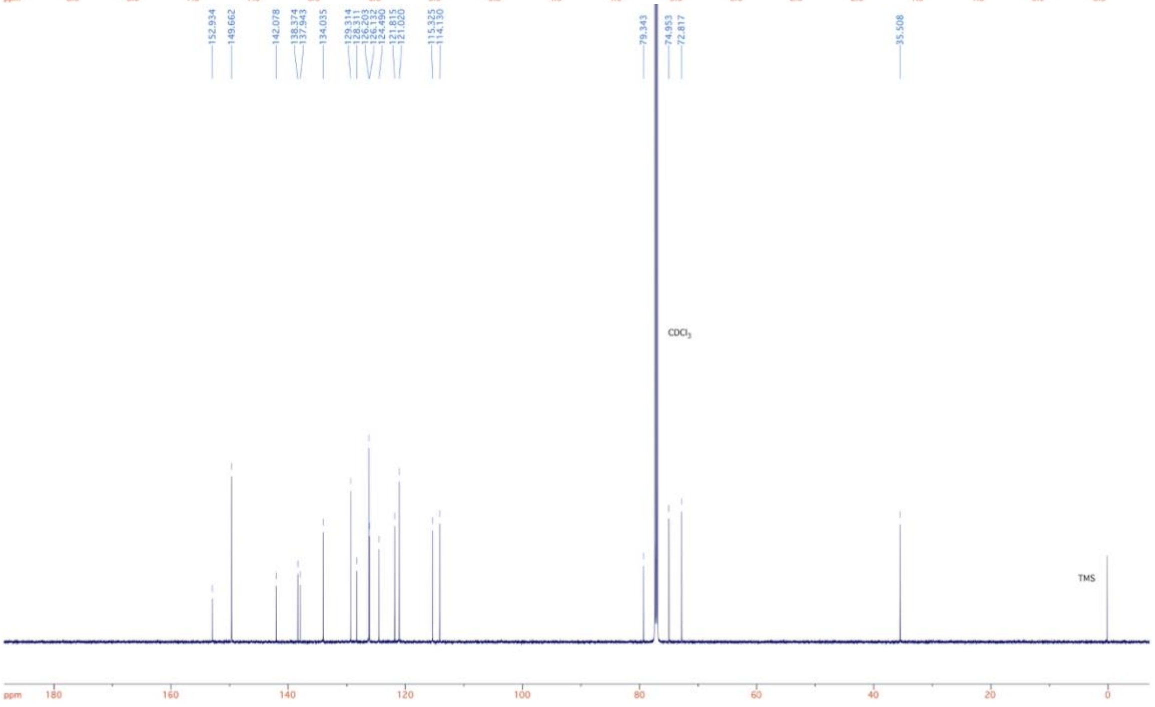

152.934
149.662
142.078
138.374
137.943
134.035
129.314
128.311
126.203
126.132
124.490
121.815
121.020
115.325
114.130
79.343
74.953
72.817
35.508
CDCl$_3$
TMS
ppm

**FKK7**

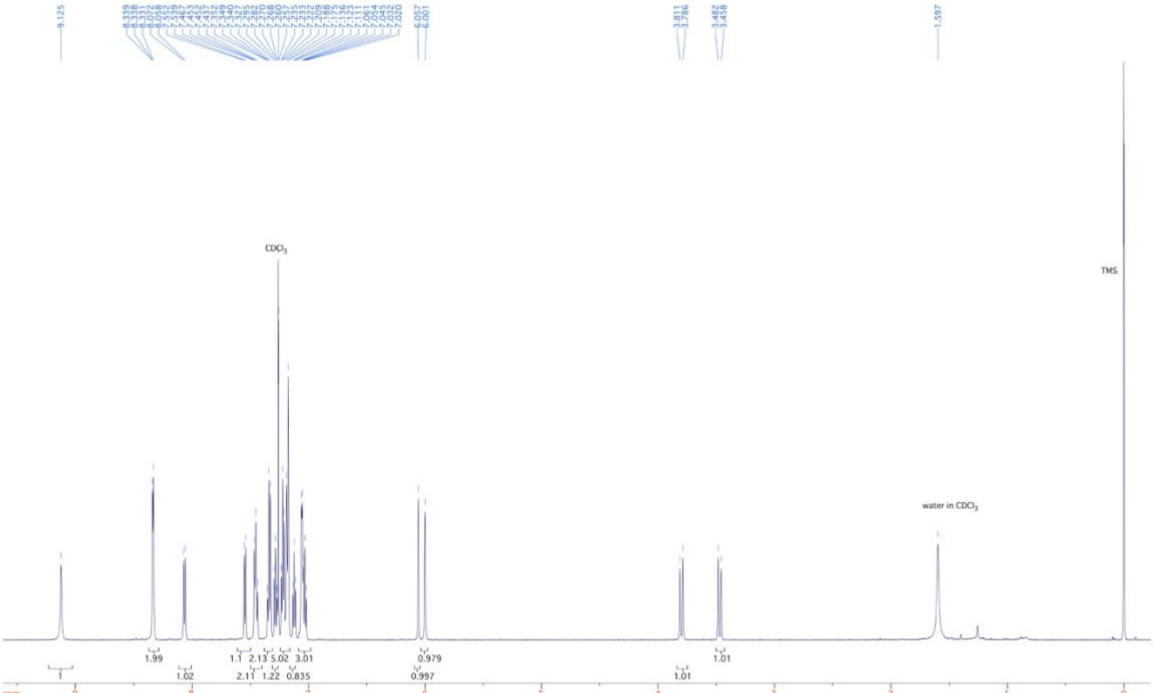

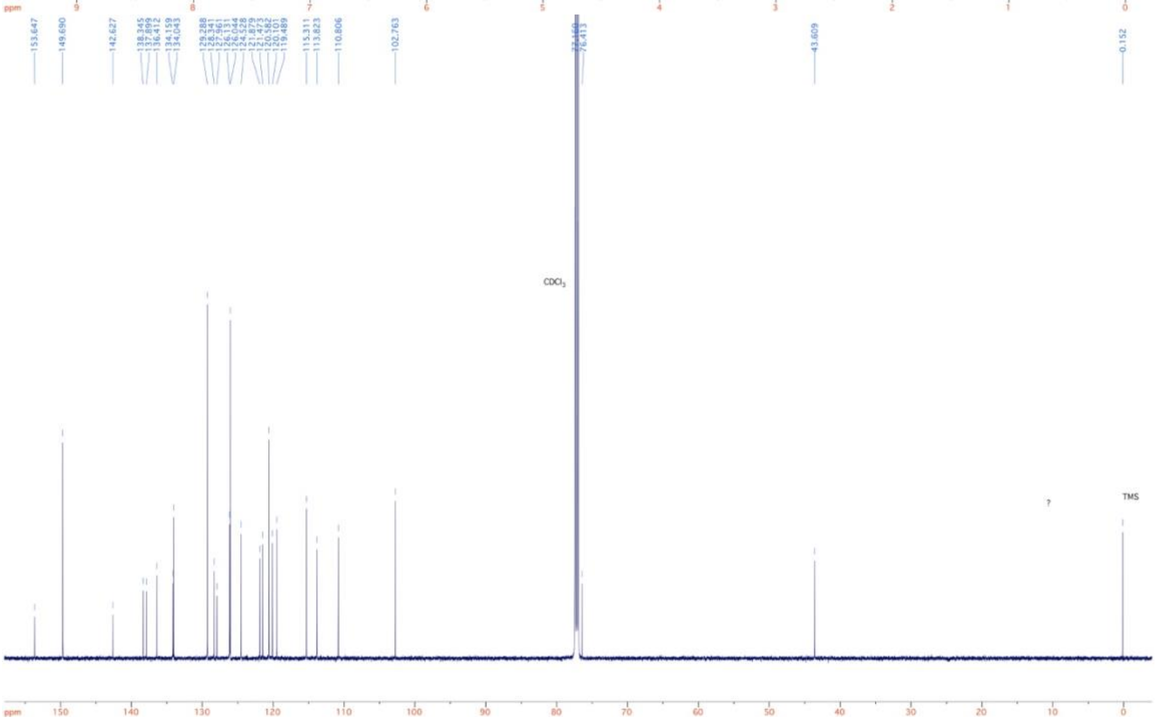

**FKK8**

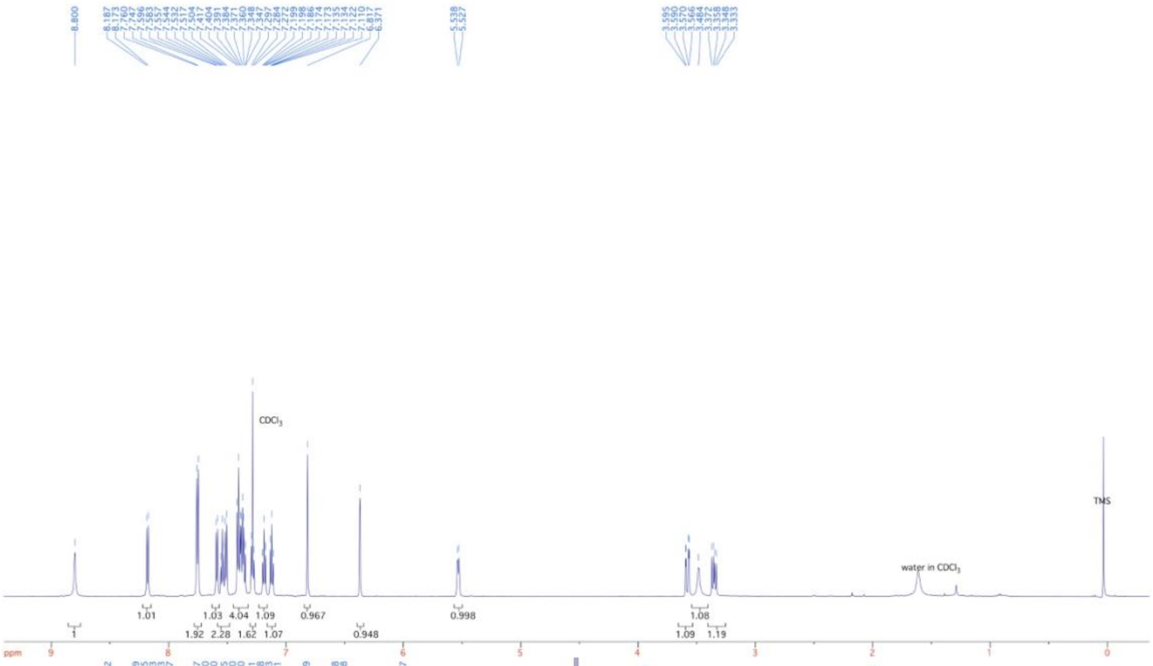

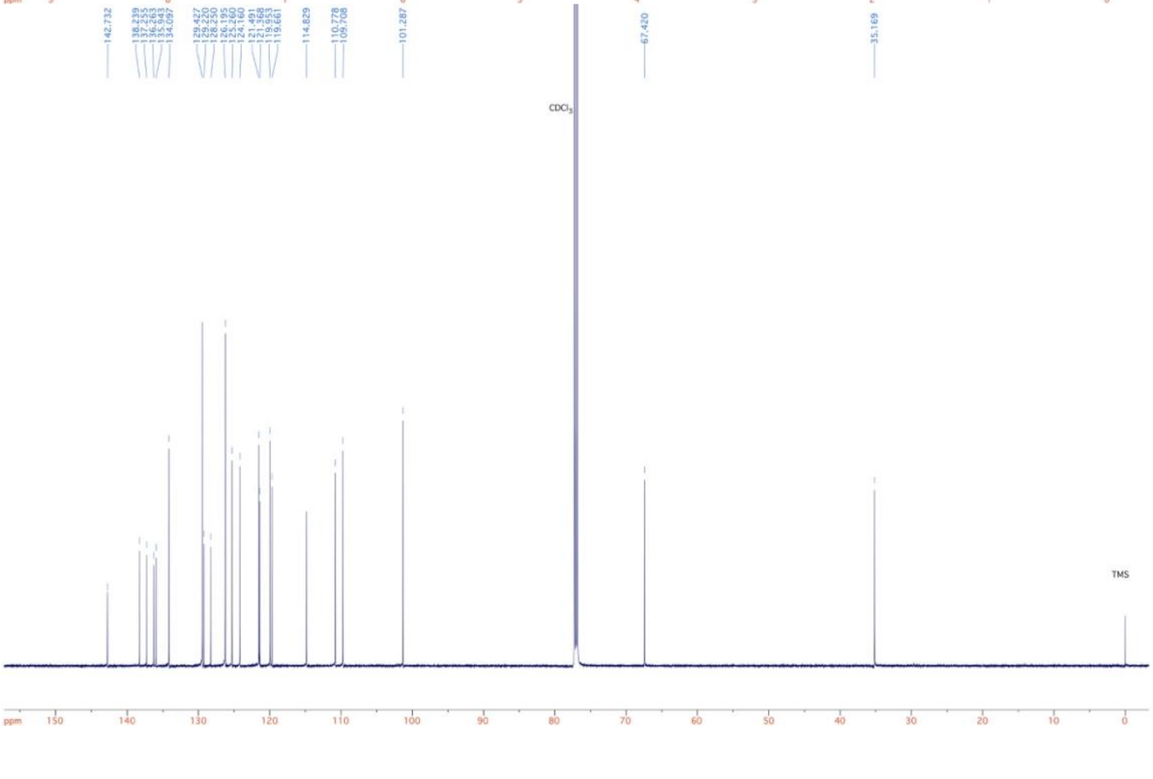

**FKK9**

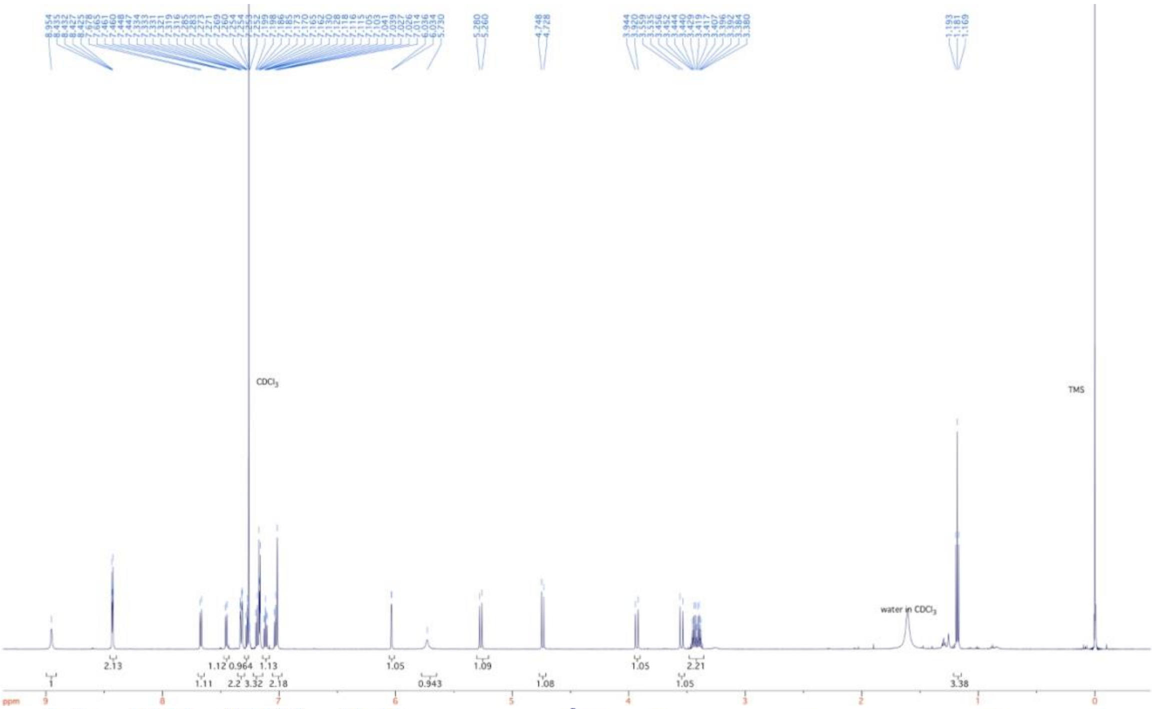
CDCl$_3$
TMS
water in CDCl$_3$
ppm

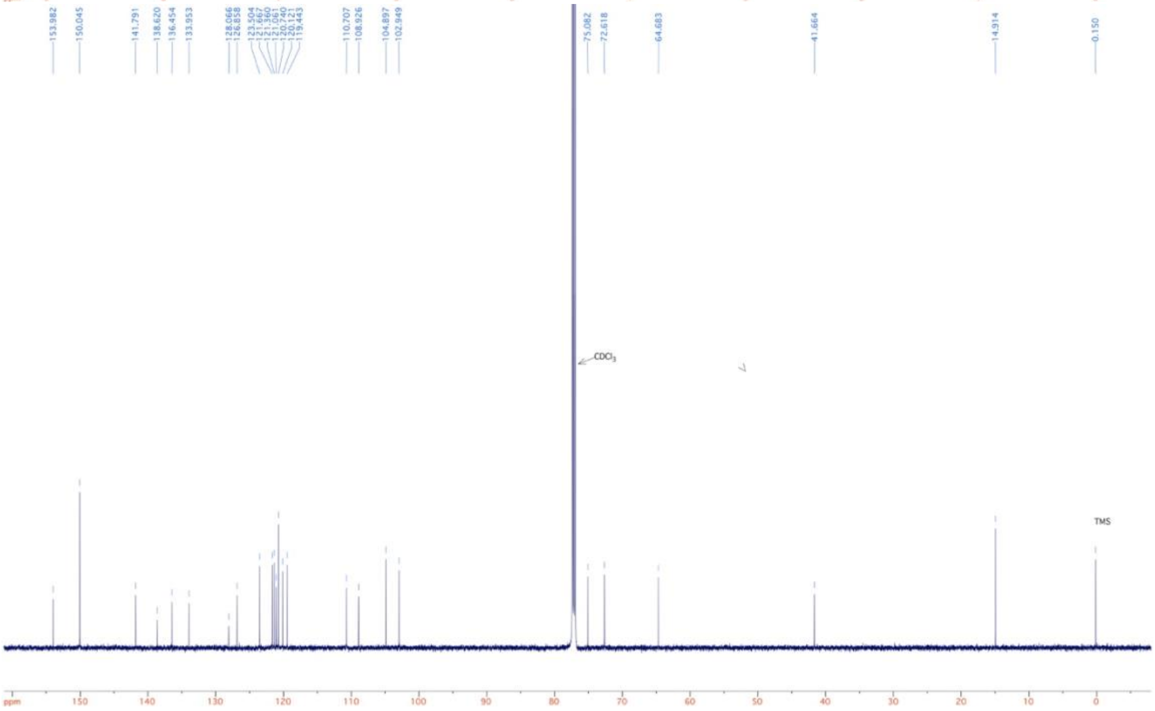
153.982
150.045
141.791
138.620
136.454
133.953
128.066
126.858
110.707
108.926
104.897
102.949
75.082
72.618
64.683
41.664
14.914
0.150
CDCl$_3$
TMS
ppm

**FKK10**

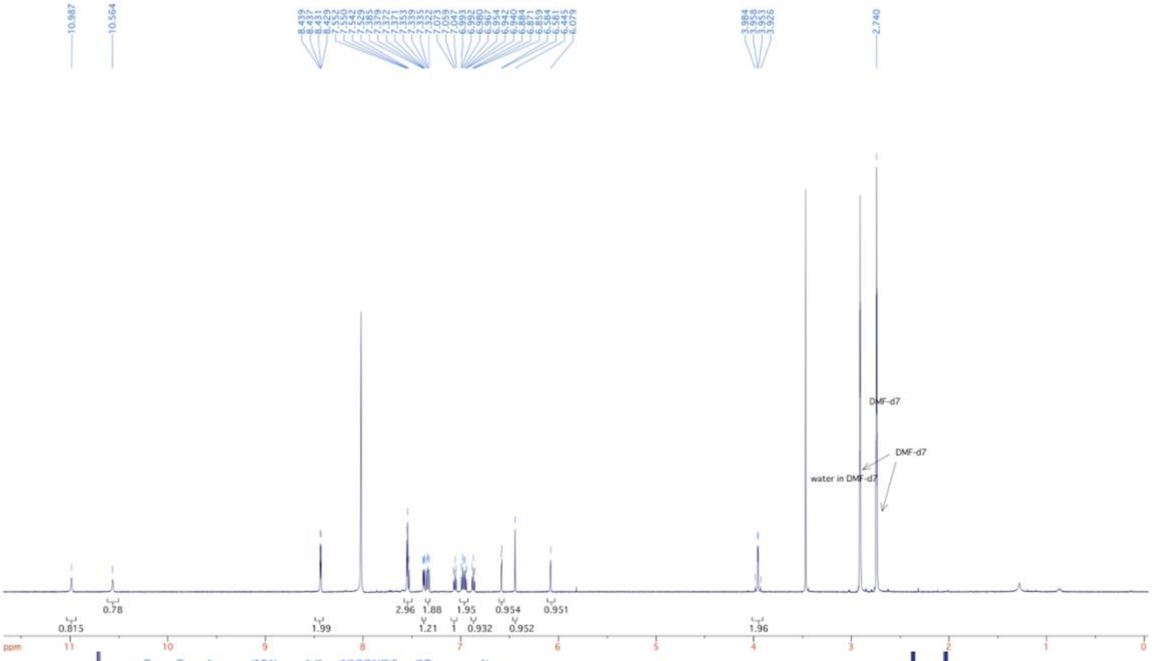
10.987
10.564
2.740
DMF-d7
DMF-d7
water in DMF-d7
0.815
0.78
1.99
2.96
1.88
1.21
1
1.95
0.932
0.954
0.952
0.951
1.96
ppm

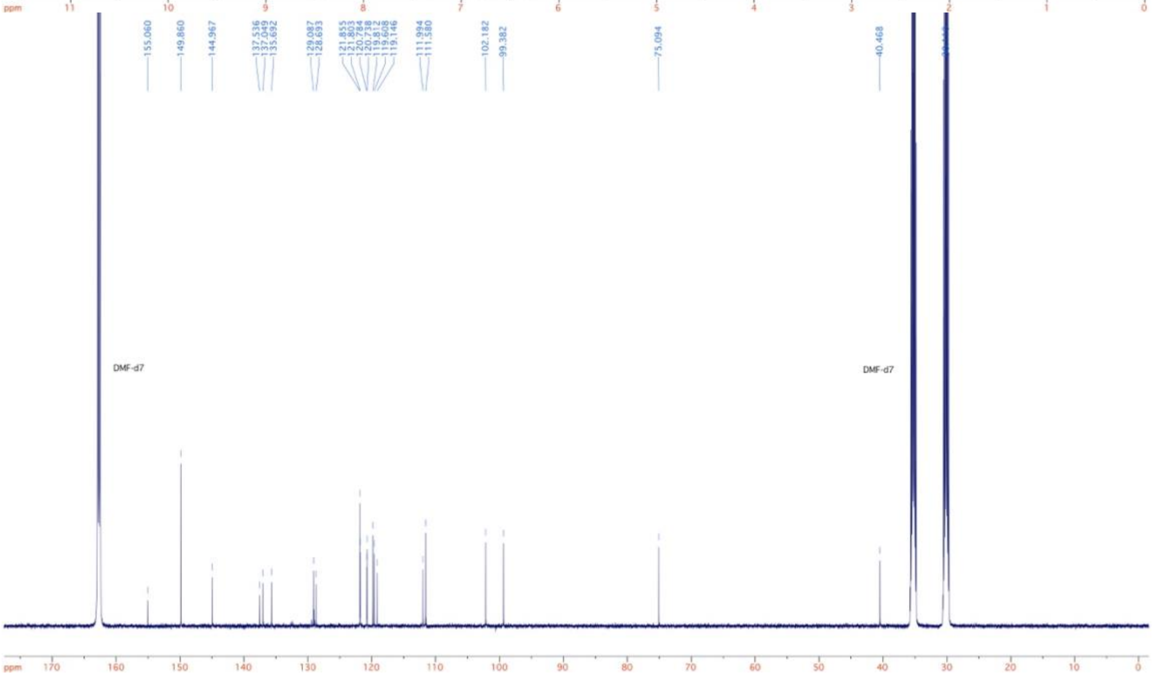
155.060
149.860
144.967
137.536
137.049
135.692
129.087
128.693
111.994
111.580
102.182
99.382
75.094
40.468
DMF-d7
DMF-d7
ppm

**Ligand efficiency metric (LEM) analysis for FKK ligand efficiency.** The ligand efficiency metrics (LEMs) of compounds FKK1 to 10 and FKK999 as a reference compound, have been analyzed by using a web tool named AtlasCBS which represents LEMs as variables for a Cartesian mapping of chemico-biological space (CBS). In particular, Cartesian planes combine three critical variables in drug discovery (affinity, size, and polarity) into easily interpretable 'efficiency planes' useful to guide a drug discovery process. In the Cartesian plane, each target:ligand pair appears as a point whose coordinates are given by pairs of values derived from the affinity of the ligand toward the target ($K_i$ or equivalent) in relation to its size (MW or number of non-hydrogen atoms NHA), and its polarity (expressed as Polar Surface Area, PSA, or number of polar atoms, NPOL = count of N + O). The first one represents a size-related LEM and the second a polarity-related LEM. Consistently, compounds that correspond to marketed drugs typically occupy the upper right quadrant of the plots ('North-East' direction, NE), where both the efficiency-per-size and efficiency-per-polarity are optimized [1,2].

Our results are represented in the efficiency plane NSEI-nBEI where:

$$NSEI = -\log K_i/NPOL\ (N, O) = pK_i/NPOL\ (N, O)$$
$$nBEI = -\log [(K_i/NHA)]$$

As a measure of activity of the above compounds, the reciprocal of the percentage efficacy values (1/Efficacy [%]) obtained in the PXR-dependent luciferase activity test ($E_{max}$ values from Fig. 1) were used in lieu of the affinity constants ($K_i$ in the above equations). FKK4 would appear as the most efficient compound since it is placed in the NE panel (GL, unpublished observations). However, this congener might suffer from metabolic liability with its *N*-protecting group possibly generating toxic aldehydes by oxidative cleavage in vivo. Unambiguously, the two most efficient compounds besides FKK4 are FKK5 and FKK6 which were therefore considered suitable lead compounds, as also suggested by PXR transactivation, target gene expression, and toxicity data (see the article main text).

**X-ray Analysis.** Single crystals of FKK5 and FKK6, respectively, were submitted for X-ray diffraction analysis. The data were collected on a Bruker X8 Kappa Apex II diffractometer using Mo Kα radiation. Crystal data, data collection and refinement parameters are summarized in Table EV1 (FKK5) and Table EV2 (FKK6), respectively. The structures were solved using a dual-space method and standard difference map techniques, and refined by full-matrix least-squares procedures on $F^2$ with SHELXTL (Version 2014/7) [3,4]. All hydrogen atoms were placed in calculated positions and refined with a riding model [$U_{iso}$(H) = 1.2–1.5$U_{eq}$(C)] with the exception of the hydrogen atoms bound to C17 and O3 in FKK6, which were located on the difference map and freely refined. TWINABS (Version 2012/1) was used to account for non-merohedral twinning in FKK5 [5]. FKK5 and FKK6 have been deposited in the Cambridge Structural Database under deposition numbers 1948848 and 1948849, respectively.

**In silico experiments**

A hybrid structure-based method has been adapted for modeling pharmacophore at PXR ligand binding pocket. Further, considering the dynamic nature of the protein, a conformational-based docking experiment has been conducted with identified FKK compounds at multiple sites in the PXR LBD (i.e., conventional ligand binding pocket, AF2 site as well as α8-pocket that was identified from earlier simulation study of PXR LBD [6]). The methodology for pharmacophore modeling and PXR ensemble-based docking are described below in sections (a) and (b).

**(a) Hybrid Structure Based (HSB) Method.** The crystal structure of hPXR (pdb code: 1M13; PMID-12578355 ) was used for the modeling and docking studies. The crystal structure was prepared by removing any bound ligands or cofactors and hydrogen atoms were added and minimized to remove any steric clashes. The protein was then solvated in a 5Å of water at each direction of an atom with largest coordination in a box with simple point-charge water model in a periodic boundary condition. The simulation parameter was performed by using CHARMM force field. All simulations were performed using NAMD program [7]. The full-system periodic electrostatic interaction was performed under Particle Mesh Ewald (PME) Sum method [8]. The temperature thermostats were a constant number, volume and temperature (NVT) steps [9,10] and constant pressure and temperature (NPT) equilibration and production MD [11,12]. A constant temperature of 310 K and the continuous pressure of 1 atm were implemented with steepest descent algorithm for energy minimization. The equilibration was performed for 0.5 ns with restraint position and followed by unrestrained position for another 0.5 ns. After the system fully equilibrated, production run was performed for 10 ns. Post simulation, the final snapshot was chosen for docking studies. Indole and Indole propionic acid (IPA) were modeled using ligand builder module of MOE program (Molecular Operating Environment; ver 2016.0801) and optimized for geometry. The two molecules were docked initially individually to the LBD of PXR using GOLD program (ver 5.2)[13] with conformational sampling of both the LBD and the ligand molecules. The best ranking pose of Indole and IPA was chosen, and a combined docking was performed and the best ranking complex of LBD with both indole and IPA was minimized and subject to 3 ns of MD simulation as described earlier.  This optimized structural complex was used to design the combined pharmacophore to screen our 3 million compounds library. The interaction profile of Indole and IPA co-docked to LBD was visually analyzed and a four-point combined pharmacophore was designed using the interaction profile of indole and IPA. Specific interactions of indole in LBD include Gln285, Phe281, Ser247, Arg287 and those of IPA include Cys301, Met323, Glu300 and a number of hydrophobic shared interactions including Trp299, Tyr306, Phe288 etc. The 4-point pharmacophore was then used to screen our library of vendor available small molecules and 5 hit molecules that strictly obeyed the pharmacophore were docked into the LBD of PXR using docking program GOLD and the complexes were scored using goldscore and chemscore functions adopted in GOLD program. FKK999 and BAS451 were chosen for in vitro testing and based on the results, FKK999's core was chosen for further medicinal chemistry optimization. Based on the core structure, ten molecules designated as FKK1–10 were designed and synthesized. All FKK molecules were modeled and docked as described previously and FKK5 and FKK6 were the lead ranking molecules.

**(b) Ensemble based-molecular docking to multiple sites.** Ensemble-based molecular docking of ten FKK compounds were performed using GOLD suite version 5.5.0 (CCDC, Cambridge, UK) [14]. GOLD uses a genetic algorithm (GA) to explore the conformational flexibility of the ligand and receptor side chains in the binding pocket. Thirty centroid conformations of *apo* hPXR generated using a RMSD-based clustering algorithm, obtained from previous work, were used for the docking [6]. In all the protein conformations, water and ions were removed prior to docking. For the docking purpose, a binding site was defined by considering all atoms within 12 Å from the geometrical center of the docking site. For each of the 30 independent GA runs, a maximum number of 200 GA operations were performed. The docked complexes were ranked with *goldscore* and then rescored using a *chemscore* fitness function [14]. All the FKK compounds were docked at the AF2 site and α8 pocket in hPXR. The scoring functions account for the hydrogen bonding, vdW interactions and steric complementarity between the ligand and receptor. For each ligand, the best-ranked docked pose with corresponding *chemscore* is considered for further analysis. Ligand interaction diagrams were prepared using LigPlot[+] software [15].

**Immunoblotting.** The cells were centrifuged (1500 rpm for 3 min), the medium was removed, and the pellet was resuspended in 150 µl of ice-cold lysis buffer (150 mM NaCl; 50 mM HEPES; 1% (v/v) Triton X-100; 5 mM EDTA; anti-protease cocktail, anti-phosphatase cocktail). The mixture was vortexed and incubated for 10 min on ice and then centrifuged (15000 rpm/13 min/4 °C). Supernatant was collected and the protein content was determined using the Bradford reagent. SDS–PAGE gels (10%) were run on a BioRad apparatus according to the general procedure followed by the protein transfer onto PVDF membrane. The membrane was saturated with 5% non-fat dried milk for 1 h at room temperature. Blots were probed with primary antibodies against PXR (mouse monoclonal, sc-48340, H-11, dilution 1:250), AHR (mouse monoclonal, sc-133088, A-3, dilution 1:500), both purchased from Santa Cruz Biotechnology (Santa Cruz, CA, USA), and β-actin (mouse monoclonal; mouse monoclonal; 3700S, 8H10D10, dilution 1:2000) purchased from Cell Signaling Technology (Danvers, Massachusetts, USA). Chemiluminescent detection was performed using horseradish peroxidase-conjugated secondary antibody and chemiluminescent substrate. As a second protocol, 1 x $10^6$ cells were centrifuged and the cell pellets were resuspended in 200 ul RIPA buffer with protein inhibitors, 20 mM Tris-HCl pH 7.0, 0.15 M NaCl, 1% NP-40, 0.1% SDS, 0.5% deoxycholic acid. The lysate was incubated for 30 minutes on ice and lysate clarification was achieved by centrifugation (12,000 rpm/10 min/4 °C). Supernatant was collected and protein content was determined using Protein A280 measurement by NanoDrop 1000 Spectrophotometer (Thermo Fisher Scientific). Twenty (20 µg) lysate protein was loaded on SDS-PAGE gels (10%) followed by transfer onto NC membrane. Goat PXR polyclonal antibody (ab109728, Abcam, Toronto, ON M5W 0E9) and goat PXR polyclonal antibody (sc-9690, Santa Cruz Biotechnology, CA) were used for blotting at 1:250 dilution, respectively.

Simple western blotting by Sally Sue™ was performed in total protein extracts from LS180 cells (± wild-type PXR). All reagents, capillaries and 384-well plates were purchased from ProteinSimple (San Jose, CA, USA) and handled according to the manufacturer's instructions. Human PXR (mouse monoclonal, sc-48340, H-11, dilution 1:5) primary antibody was purchased from Santa Cruz Biotechnology Inc. (Santa Cruz, CA, USA). β-actin (mouse monoclonal; 3700S, 8H10D10, dilution 1:1000) primary antibody was obtained from Cell Signaling Technology (Danvers, Massachusetts, USA). Target proteins were detected by conjugation of specific primary antibodies with horseradish-conjugated secondary antibody followed by reaction with chemiluminescent substrate. Data were analyzed using the Compass Software version 2.6.5.0 by ProteinSimple.

**Kinase Assays.** The kinase inhibition assays were conducted by DiscoverX on a platform assay by Ambit BioSciences as described previously [16,17]. FKK6 was screened at a single concentration of 10 μM in duplicate. The *scan*MAX[SM] assay panel measures 468 kinases (https://www.discoverx.com/services/drug-discovery-development-services/kinase-profiling/kinomescan/scanmax). The assays measure a compound's ability to inhibit binding of a canonical ligand substrate, with 0% activity of control corresponding to full inhibition and 100% activity of control to no inhibition. This data is then used to calculate a selectivity score (S-score), which is a quantitative measure of compound selectivity based on the number of kinase hits (enzymes inhibited) divided by the total number of distinct kinases tested. Three different activity cut-offs were analyzed – S(35) or % activity of control < 35; S(10) or % activity of control < 10; and S(1) or % activity of control < 1. To illustrate effect or lack of effect, an image of the entire family of kinases tested were grouped by families and S(35). The TREE*spot* Kinase dendrogram image was generated using TREE*spot*™ Software Tool and reprinted with permission from KINOME*scan*®, a division of DiscoveRx Corporation, © DISCOVERX CORPORATION 2010. FKK5 at concentrations of 10 and 25⌠M were additionally screened against the following kinase assays (CDK2, CDK4, CDK5, MAPK3, MAPK1, MAPK6, MAPK4, MAPK7, MAPK15, GSK3A, GSK3B, CHUK, MAPK8, MAPK9, PRKACA, PRKACB, RPS6KB1) in two separate experiments each performed as a single read (KINOME*scan*[TM] Profiling Service).

The following kinase assays (HDAC, SIRT1, PRMT1, PARP1, p300, pCAF) were performed by Reaction Biology Corp. FKK6 was screened at a single concentration of 10 μM in duplicate in two separate experiments. The appropriate controls for the individual assays are published. The overall enzyme activity in the assay is reported as a % activity in DMSO control wells. Control $IC_{50}$ concentrations are shown in the Table. Briefly, for the Sirtuin (SIRT1) assay, human SIRT1 (GenBank

Accession No. NM012238) full-length N-terminal His-tag protein (MW 83.2 kDa) was purified after expression in *E. coli*. The reaction buffer was freshly prepared and constituted 50 mM Tris-HCl (pH 8), 137 mM NaCl, 2.7 mM KCl, 1 mM $MgCl_2$, and just before use, 1 mg/mL BSA and 1% DMSO was added. The substrate was a fluorogenic acetylated peptide of p53 residues 379–382 (RHKK (Ac)-AMC). In the multi-well format, the 2X enzyme was mixed with the reaction buffer followed by compound (in 100% DMSO) titration via Acoustic technology (Echo550; LabCyte Inc, Sunnyvale, CA) in the nanoliter range. The mixture was spun down and 2X substrate (including $NAD^+$) was added to initiate the reaction at 30°C at 2h under seal. The reactions were subsequently quenched using a developer with nicotinamide. Fluorescence kinetic measurements were conducted for 1.5 h in 15 min intervals using Envision (Ex 360nm/Em 460nm). For the Poly (ADP-ribose) Polymerases (PARP) assay, PARP protein (RBC Cat# PAR-21-346) 2.5 nM was mixed with reaction buffer (50 mM Tris-HCl (pH 8), 50 mM NaCl, 10 mM $MgCl_2$, 0.02% Brij35, 1 mM DTT, 1% DMSO, and 20 μg/mL activated DNA (Sigma Cat# D4522) and substrate solution consisting of 0.01 mg/mL Core Histones from chicken (RBC Cat # HMT-35-435). In the multi-well format, the enzyme mixture was mixed with the compounds (in 100% DMSO) titrated via Acoustic technology (Echo550; LabCyte Inc, Sunnyvale, CA) in the nanoliter range. The compound mixture was incubated for 20 min at room temperature following which $^{32}P$-$NAD^+$ (PerkinElmer Cat# BLU023X250UC) was added to the reaction mixture and incubated for 1 h at room temperature. The compound mixture was placed on filter paper and washed with 0.75% phosphoric acid and radioactivity counted. The histone methyltransferase assay protocol is previously published [18]. The Histone Acetyltransferases (HAT) assay, proceeds as described for the other assays, however, uses *E. coli* expressed human recombinant pCAF catalytic domain (aa 492–658) (GenBank accession number: NM_003884, MW 19.3 kDa) or p300 (aa 1284–1672) (GenBank accession number: NM_001429, MW 45.1 kDa). The reaction buffer is 50 mM Tris-HCl (pH 8), 50 mM NaCl, 0.1 mM EDTA, 1 mM DTT, 1 mM PMSF, and 1% DMSO. The substrate is $^{3}H$-CoA (acetyl coenzyme A) and the incubation time is for 1 h at 30°C. The radioactivity in mixture is detected on washed filter paper using a scintillation counter. The Histone Deacetylase (HDAC) fluorescent activity assay (HDAC1-3) is based on the using 2X purified human enzyme mixed with the reaction buffer (50 mM Tris-HCl pH 8, 137 mM NaCl, 2.7 mM KCl, and 1 mM $MgCl_2$, 1 mg/mL fresh BSA and 1% fresh DMSO) and then compounds (in 100% DMSO) are titrated in using Acoustic Technology. The 2X HDAC substrate is a fluorogenic acetylated peptide from p53 (residues 379 - 382 RHKK(Ac)AMC) which is mixed in with the enzyme-compound complex and incubated for 1-2 h at 30 °C. In the development step, a proprietary developer with trichostatin A (or TMP269) is added to stop the reaction and to generate fluorescent color which is read using an EnVision Multilabel Plate Reader (Perkin Elmer) (Ex 360 nm/ Em 460 nm). For all enzyme activity assays, the percentage of enzyme activity (relative to DMSO controls) and $IC_{50}$ values were calculated using GraphPad Prism 4 program (sigmoidal dose response equations).

**Isolation of cell nuclei.** Rat liver nuclear fraction was prepared from rat livers by the procedure established and optimized previously [19]. Nuclear fraction can be prepared either from fresh or deep-frozen liver tissues (-80 $_{o}$C), and all subsequent steps have to be carried out at 0–4 $_{o}$C. The liver tissue was at first washed in SM solution (0.32 M sucrose, 1 mM $MgCl_{2,}$ 0.1 mM PMSF and freshly prepared 1 mM DTT), consequently cut into smaller pieces and homogenized by the glass-teflon homogenizer (8–10 smooth strokes). Thus, prepared homogenate was centrifuged at 1 000 x *g* for 10 minutes. Crude pellet was again washed with SM solution and thus prepared pellet was mixed with 2.3 M sucrose solution, with 1 mM $MgCl_{2,}$ 0.1 mM PMSF and 1 mM freshly prepared DTT, and treated by isopycnic ultracentrifugation at 220 000 x *g* for 40 minutes. Isolated nuclei were washed twice with SMCT solution (0.32 M sucrose, 10 mM Tris-HCl, 1 mM $MgCl_2$, 1 mM $CaCl_2$, 0.1 mM PMSF and freshly prepared 1 mM DTT at pH 7.4) once in the presence of 0.25% Triton X-100 and once in its absence. Nuclear protein fraction containing RXRs was obtained from purified nuclear pellet with a high ionic strength buffer containing 0.3 M KCl, with the addition of 1 mM $MgCl_2$,10 mM Tris-HCl and freshly prepared 1 mM DTT at $0_oC$ for 60 minutes followed by ultracentrifugation at 135 000 x *g* for 60 minutes. Concentration of the prepared nuclear fraction of proteins was estimated by Lowry's method using bovine serum albumin as a standard.

**Labeled[11,12-$^3$H(N)] 9-*cis* retinoic acid (9*c*RA) binding competition assay (RXR ligand binding)** [19]**.** The assay on labeled 9*c*RA binding to nuclear proteins was performed in dark at $20_oC$, in a high ionic strength buffer (0.3 M KCl, 1 mM $MgCl_2$, 10 mM Tris-HCl buffer, and pH 7.0). Samples for total radioligand binding to its cognate nuclear receptors were incubated with 0.66 nM of [11,12-$^3$H(N)]9*c*RA (Canberra Packard, USA) and samples confirming displacement of bound radioligand from nuclear receptors were incubated either with 0.66 nM of [11,12-$^3$H(N)]9*c*RA together with 60-fold molar excess of 9*c*RA (39.94 nM). Samples evaluating displacement of bound radioligand from nuclear receptors by FKK compounds were incubated with 0.66 nM of [11,12-$^3$H(N)]9*c*RA together with 60-fold molar excess of respective FKK compounds (39.94 nM), in 3 µl ethanol solution per 0.5 mL sample for 120 minutes. After incubation, 0.5 mL of cold (0□4 $_o$C) charcoal-dextran suspension in a high ionic strength buffer containing 0.3 M KCl, 1 mM $MgCl_2$, 10 mM Tris-HCl buffer (pH 7.0) was added to each sample. After short vortexing, the suspension was centrifuged at 1 500 x g for 10 minutes. Then 0.5 mL of supernatant was transferred and mixed with 6 mL scintillation solution (INSTA-gel, Perkin Elmer, USA) and its radioactivity was quantified in the liquid scintillation counter Tri-Carb 2910 TR (Perkin Elmer, USA).

**hFXR TR-FRET.** The FXR ligand binding assay of FKK5, and FKK6 was performed by the LanthaScreen TR-FRET FXR Coactivator Assay Kit (PV4833; Invitrogen, USA) according to manufacturer's instructions, with GW4064 as a model FXR ligand. The assays were performed in a volume of 20 µl in 384-well black plates, with concentrations of tested compounds in the range from 1 nM to 25 µM. Dimethyl sulfoxide (DMSO; 1% v/v) and GW4064 (500 nM) were used as a negative and positive control, respectively. The reaction mixture was incubated at room temperature for 1 hour in a dark, and then fluorescent signals were measured at 495 nm and 520 nm, with the excitation filter 340 nm, on Infinite F200 microplate reader (Tecan Group Ltd, Switzerland). Finally, the TR-FRET ratio was calculated by dividing the emission signal of 520 nm by that at 495 nm. All experiments were done in tetraplicates and as two independent experiments. Final $EC_{50}$ values were obtained by processing the data with GraphPad Prism 6 using standard curve interpolation (sigmoidal, 4PL, variable slope).

**PPARγ reporter assay.** Transcriptional activity of PPARγ was studied in stably transfected reporter cell line PAZ-PPARγ [20]. Cells were seeded in 96-well plates, stabilized for 24 h, and then incubated with tested compound. Dimethyl sulfoxide (DMSO; 0.1% v/v) and 15-deoxy-δ12,14-prostaglandin J2 (15d-PGJ2; 40 µM) were used as a negative and positive control, respectively. After the treatments, cells were lysed and luciferase activity was measured on Tecan Infinite M200 Pro plate reader (Schoeller Instruments, Prague, Czech Republic). The data are expressed as fold induction ± SD of luciferase activity over the control cells. Differences were tested using one-way ANOVA with Dunnett's post hoc test, $p < 0.05$, was considered significant (*).

#### Appendix Figures

#### Appendix Figure S1

a.

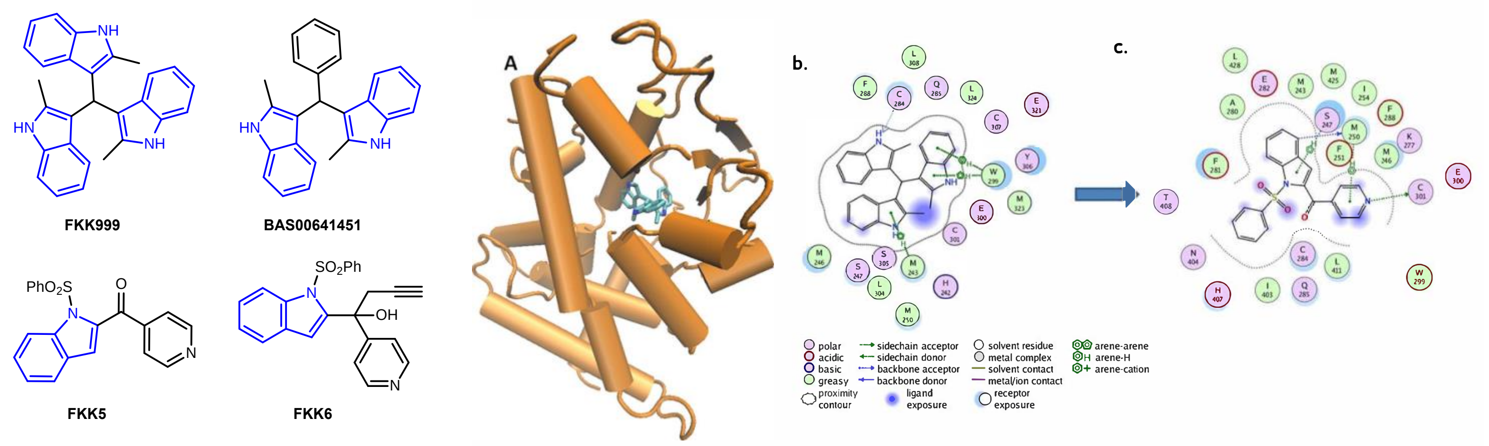

d.

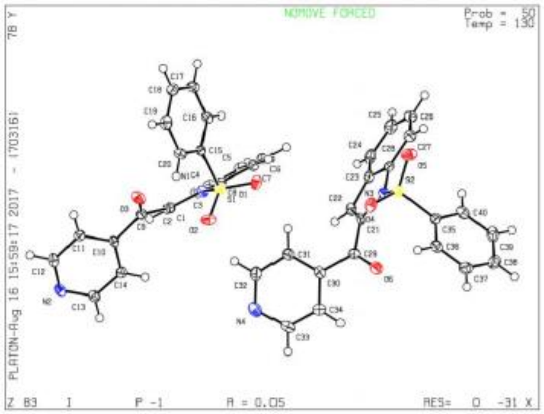

e.

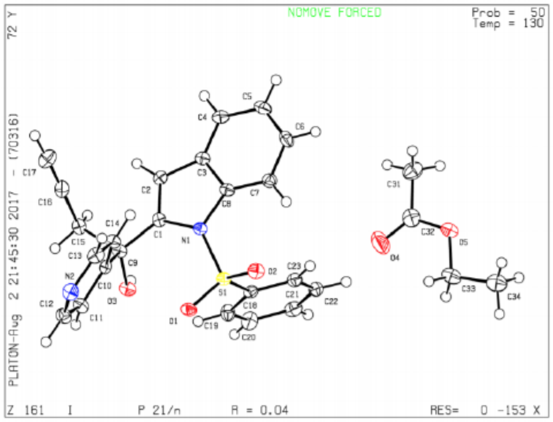

**Fig. S1: FKK compound structures and their docked complex with hPXR. a,** Chemical structures of FKK compounds and BAS00641451. Indole is colored blue. hPXR is shown as cartoon model and colored orange (A), **b**, FKK999 is shown as licorice sticks and colored atom type (C-cyan, N-blue, O-red). Hydrogen atoms are removed for clarity. Schematic representation of **b**, FKK999 and **c**, FKK5 in the binding pocket of hPXR (all residues within 4.5 Å). The legend below the figure describes the nature of interactions. **b**,**c**, were generated using the ligX module of MOE modeling package (ver 10.1). **d**, ORTEP drawing of FKK5. The crystal is non-merohedrally twinned, which causes the $R_{int}$ value to be slightly higher than usual. There are two molecules in the asymmetric unit. They are related by inversion symmetry, but there does not appear to be a crystallographic inversion center between them; attempts at using a smaller unit cell or a higher symmetry space group were not successful. **e**, ORTEP drawing of FKK6. The compound crystallized in a centrosymmetric space group, indicating that the crystal contains a racemic mixture. There is one molecule of ethyl acetate co-crystallized with FKK6. The OH group forms an intermolecular hydrogen bond with the pyridine group.

### Appendix Figure S2

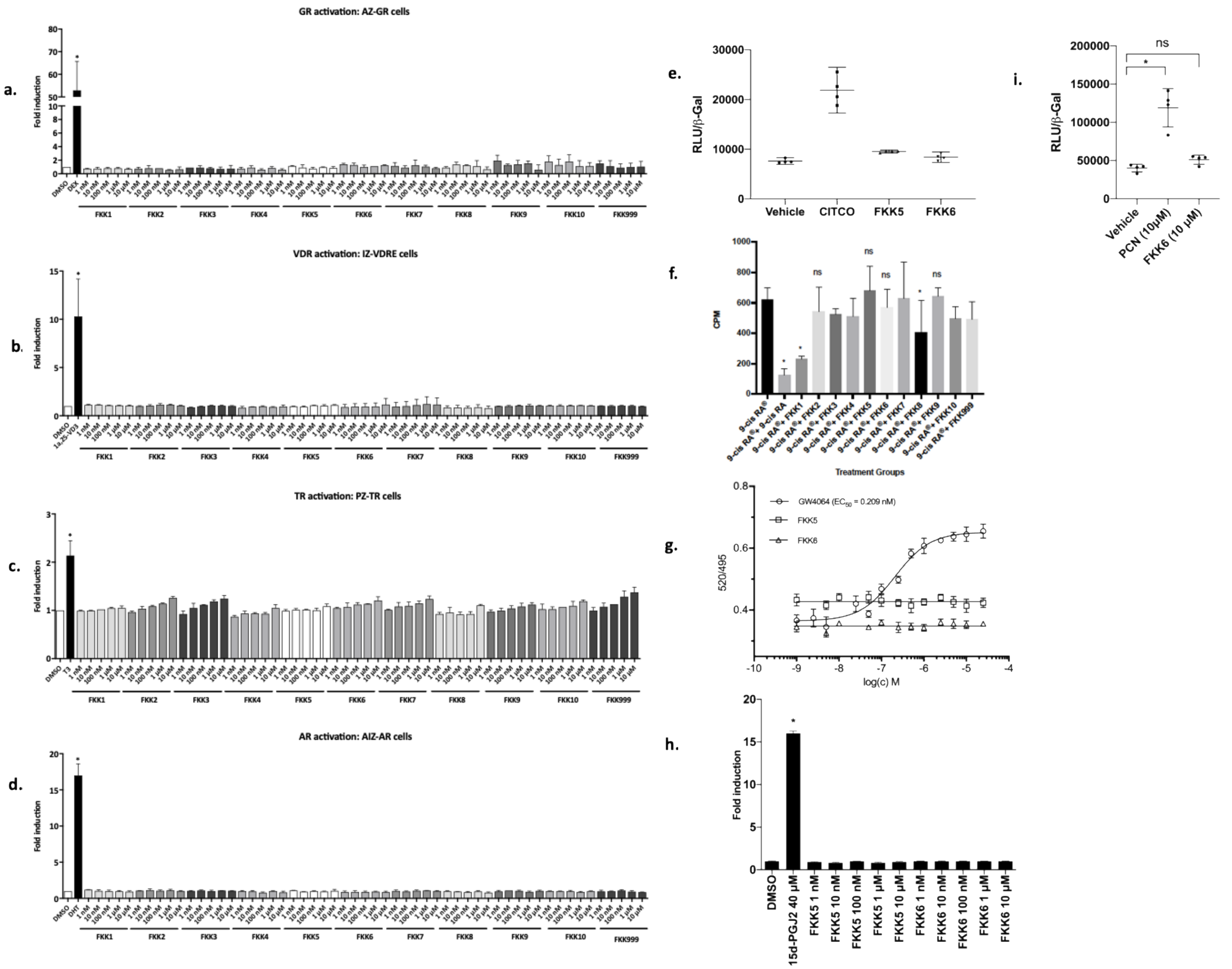

**Fig. S2: FKK5 and FKK6 do not activate steroid and related nuclear receptors.** Transcriptional activity of the respective nuclear receptors after incubations in the presence of their respective ligands. Transgenic AZ-GR, IZ-VDRE, PZ-TR, AIZ-AR and PAZ-PPARγ cells were incubated with tested compounds and the model ligands, i.e. dexamethasone (DEX; 100 nM), calcitriol (1α,25-VD3; 50 nM), 3,5,3'-triiodothyronine (T3; 10 nM) or dihydrotestosterone (DHT; 100 nM), 15-deoxy-δ12,14-prostaglandin J2 (15d-PGJ2; 40 µM), respectively. **a**, Glucocorticoid Receptor. **b**, Vitamin D Receptor. **c**, Thyroid Receptor. **d**, Androgen Receptor and **h**, Peroxisome Proliferator-Activated Receptor gamma (PPARγ). **a-d,** The histogram depicts mean ± SD from experiments performed in two consecutive passages of cells. **h,** histogram depicts mean ± SD from one of two independent experiments each performed in triplicate. * $p < 0.05$ , Two way ANOVA. **e**, human constitutive androstane receptors (hCAR) reporter activity in HEK293T cells. The RLU to beta-galactosidase ratio was calculated for CAR ligand CITCO, FKK5, FKK6 and vehicle control. This data represents one experiment performed in quadruplicate. Assays were performed at least two separate times, each in quadruplicate. **f**, displacement capacity of 9-*cis* retinoic acid and FKK compounds tested for their potential capability to play a role as RXRs ligands. [11,12-$^3$H(N)] 9*c*RA (0.66 nM) was applied in all groups. Compounds from left: 9-*cis* retinoic acid (39.94 nM), FKK1 – FKK 10 and FKK999 (39.94 nM) (**$p < 0.01$, *$p < 0.05$ vs total [11,12-$^3$H(N)] 9*c*RA binding , student t-test). **g**, FXR TR-FRET assay. TR-FRET ratio 520/495 nm are plotted against concentration of compound(s). Half-maximal inhibitory concentrations $IC_{50}$ were obtained from interpolated standard curves (sigmoidal, 4PL, variable slope); error bars show standard deviation of *n* = 2 independent experiments each with four technical replicates. **i**, mouse Pregnane X receptor (mPXR) reporter (MRP2) activity in HEK293T cells. The RLU to beta-galactosidase ratio was calculated for PXR ligand pregnane carbonitrile (PCN), FKK6 and vehicle control. This data represents one of two independent experiments, each performed in quadruplicate. The histogram depicts mean ± SD. * $p < 0.05$, One way ANOVA.

### Appendix Figure S3

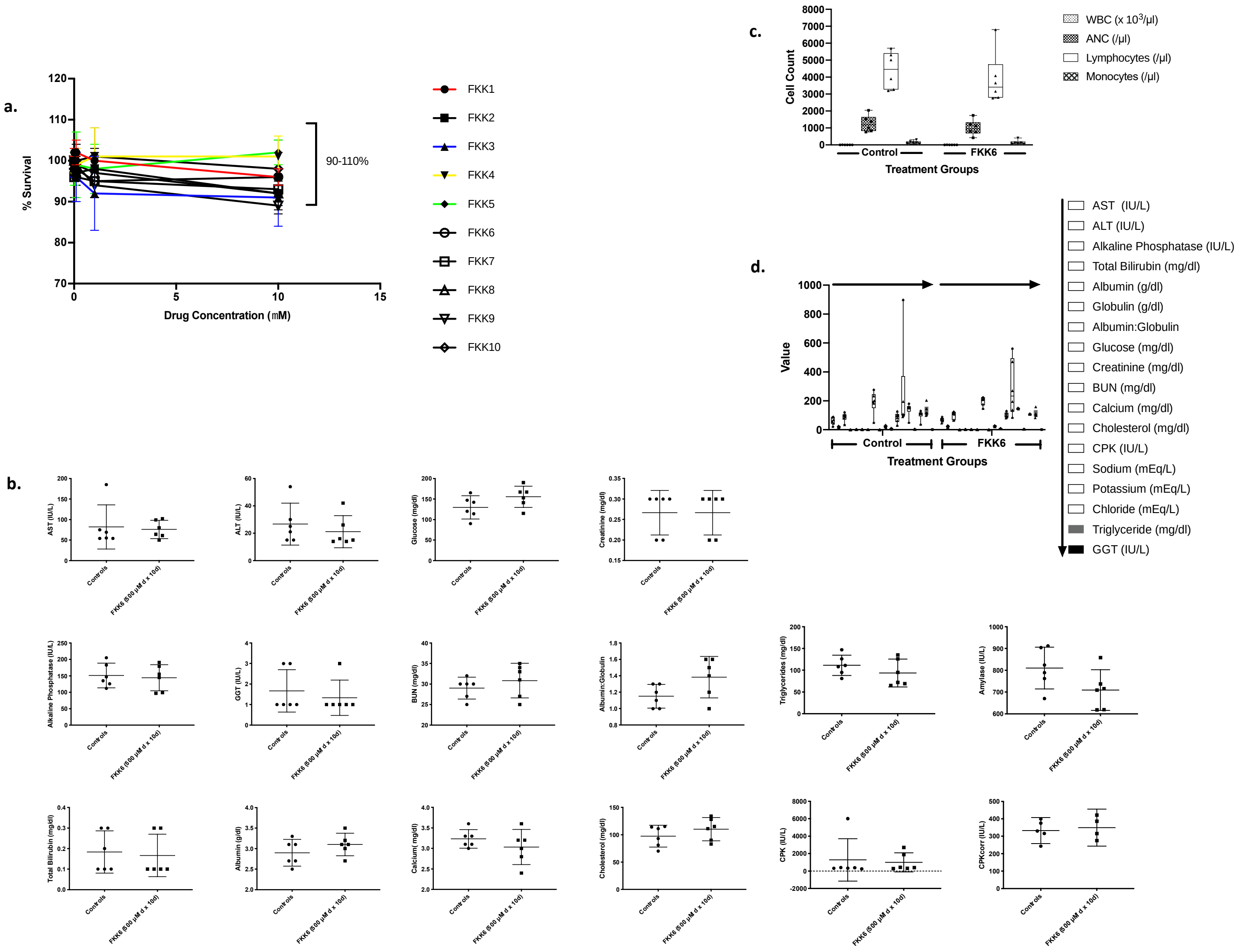

**Fig. S3: FKK6 is not cytotoxic to cells and mice. a,** Cell cytotoxicity (MTT) assays of FKK compounds in LS180 cells. Assays were repeated at least three independent times, each in quadruplicates. Data presented as mean (± SD). **b**, serum chemistry panel of mice exposed to FKK6 (n = 6) or Vehicle /Controls(*n* = 6). Studies were performed on day 10. **c**, complete blood count and **d**, serum chemistry panel of mice exposed to FKK6 (*n* = 6) or Vehicle /Controls (*n* = 6). Studies were performed on day 30. **b – d**, the data are plotted as mean (95%CI); not significant (*p* > 0.05) in any panel, student t-test. CPKcorr, corrected CPK values

#### Appendix Figure S4

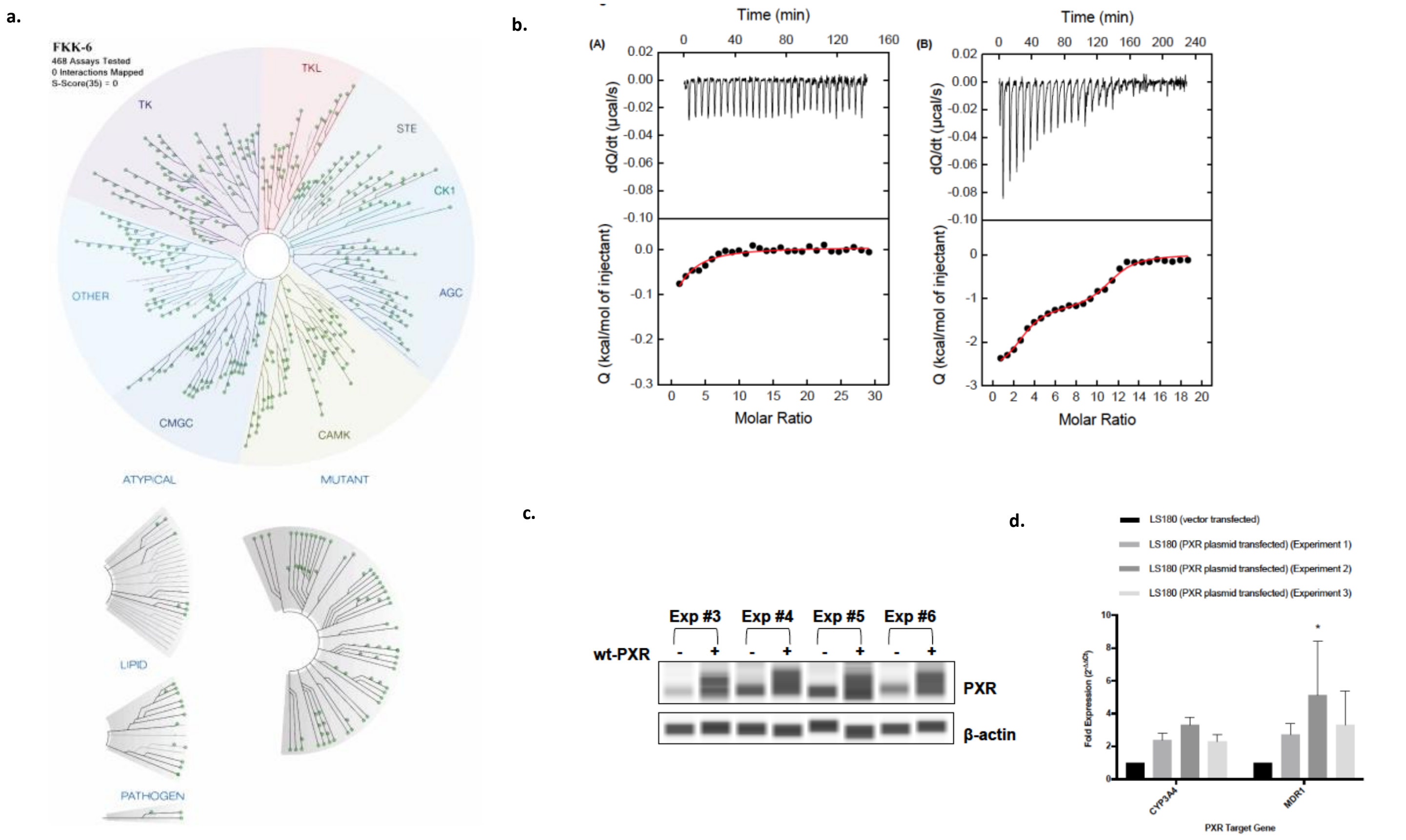

**Fig. S4: FKK6 does not inhibit a broad array of kinase enzymes. a,** TREE SpotTM Interaction Maps. The distribution of kinases and its activity in the presence of FKK6 (10 µM) in the screening panel is represented by green dots (kinase activity percent control > 35%) on a dendrogram representing the human kinome (kinome image is reproduced courtesy of TREE*Spot*™, DiscoverX proprietary data visualization tool, KINOME*scan*). **b**, Microcalorimetric titrations of PXR ligand binding domain (LBD) with (A) 3-indole propionic acid and (B) rifampicin. **c**, Confirmation of transfection PXR in LS180 cell line. The levels of PXR and β-actin proteins were determined by quantitative automated western-blot analyzer SallySue from the ProteinSimple software. wt-PXR, LS180 cells transiently transfected with wild-type human PXR plasmid; Exp (#number), number of the experiment. **d**, PXR expression in LS180 cells induces CYP3A4 and MDR1 mRNA expression. Expression data were normalized to the housekeeping gene GAPDH. Data are expressed as fold induction ± SD over the mock-transfected cells. Differences were tested using one way ANOVA with Dunnett's post hoc test, * $p < 0.01$.

#### Appendix Figure S5

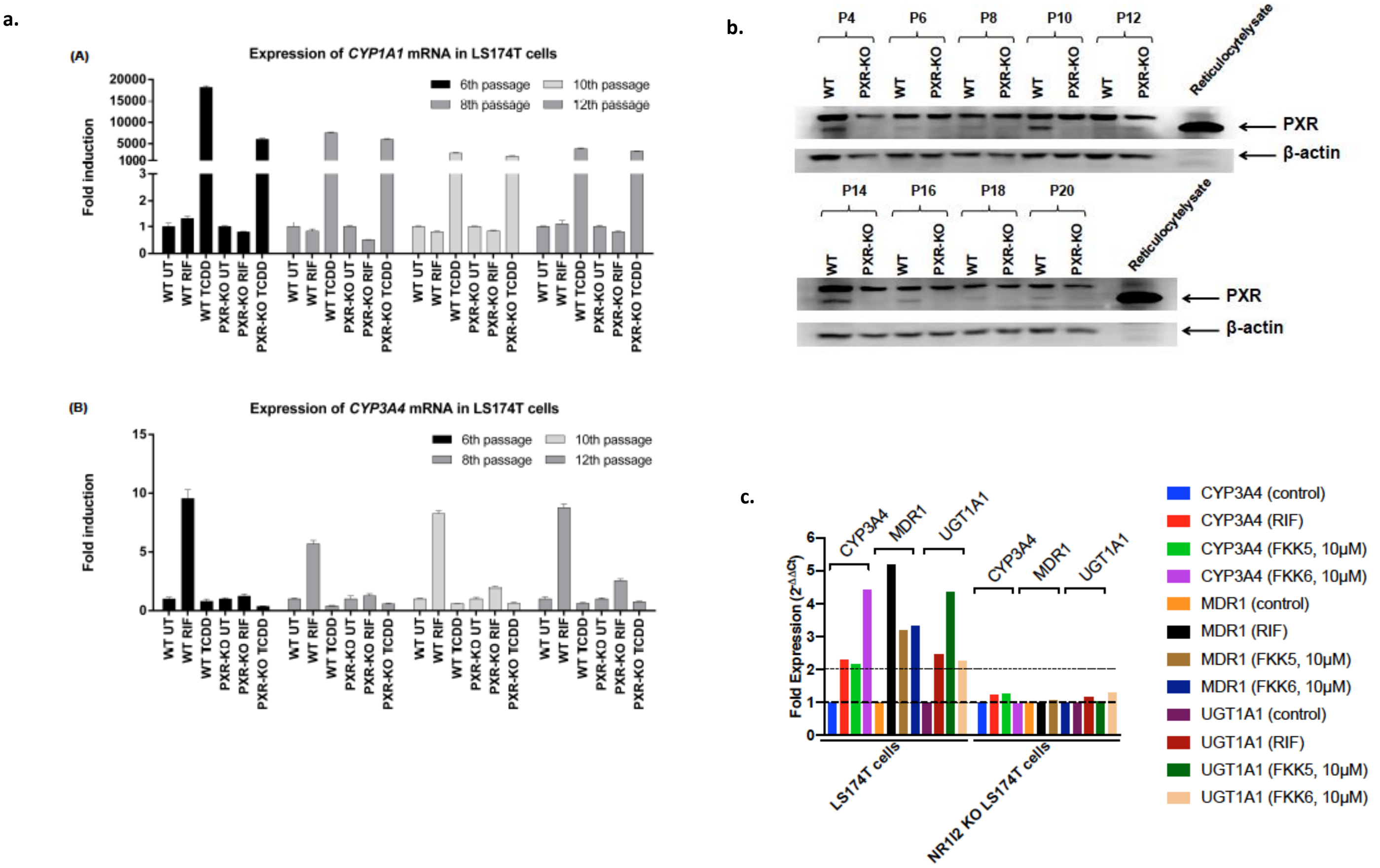

**Fig. S5: LS174T PXR knockout (KO) cells are not responsive to PXR ligands. a,** Expression of CYP1A1 and CYP3A4 mRNA in PXR-knock out and wildtype LS174T. Cells were treated with vehicle (UT; DMSO; 0.1% v/v), rifampicin (RIF; 10 μM), or 2,3,7,8-tetrachlordibenzodioxin (TCDD; 5 nM) for 24 hours. Experiment was performed in 4 independent cell passages (6th, 8th, 10th, and 12th). The mRNA expression of CYP1A1 and CYP3A4 mRNA was determined by RT-PCR and data were normalized to GAPDH mRNA level. (A) CYP1A1 mRNA. (B) CYP3A4 mRNA. Data are expressed as mean ± SD. **b**, immunoblot for PXR protein expression in PXR wild-type and knockout (KO) cells. Experiment was performed in nine independent cell passages (P). **c**, Fold mRNA expression of PXR target genes. Each tissue PCR was repeated in quadruplicate. The expression was normalized to internal control, GAPDH. The histograms show mean (95% CI) values and two way ANOVA with Sidak's post hoc test. The entire experiment was repeated n > 2 for reproducibility. The data shown is one representative experiment.

#### Appendix Figure S6

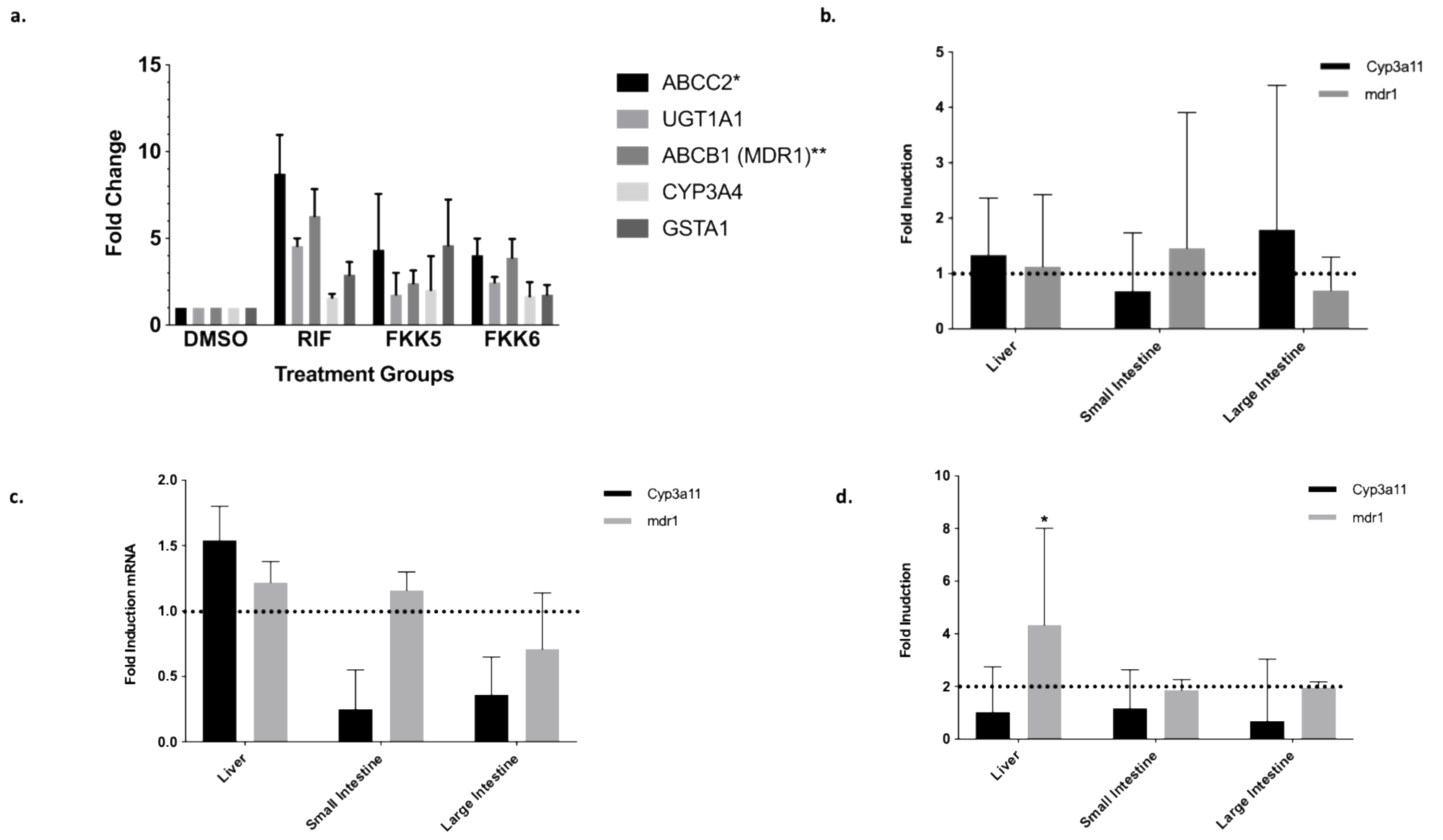

**Fig. S6: FKK6 induces PXR target gene expression in human duodenal enteroids and mice. a**, Fold change in mRNA expression of PXR target genes (ABCC2, UGT1A1, ABCB1, CYP3A4, GSTA1) in human duodenal enteroids (over 40 enteroids per sample). The graph shows mean (95% CI) fold change for each sample (n = 3). The data were normalized to H18S expression. *, ** $p < 0.05$ , two-way ANOVA followed with Dunnett's post hoc test. **b**, C57BL/6, **c**, *pxr* -/- and **d**, *hPXR mice* were administered 500 μM FKK6 or vehicle 10% DMSO by oral gavage every 12h for 5 consecutive doses (*n* = 3 mice/group). Mouse small, large intestine and liver samples were collected and total RNAs were isolated. RT-qPCR data show fold induction of respective gene mRNAs (TLR4, CYP3A4, MDR1). Each tissue PCR was repeated in quadruplicate. The expression was normalized to internal control, GAPDH. (**b,d**) The histograms show mean (95 %CI) values, **c**, mean (SD) and two way ANOVA followed with Tukey's post hoc test. The entire experiment was repeated *n* = 3 for reproducibility. The data shown is one representative experiment.

#### Extended View Results (Tables)

**X-Ray Crystallography**
Samples for X-ray crystallography were crystallized from ethyl acetate. Refer to Supplementary Fig. 1 for structures.

**Table EV1.** Crystal, intensity collection, and refinement data.

| | **FKK5** |
|---|---|
| lattice | Triclinic |
| formula | $C_{20}H_{14}N_2O_3S$ |
| formula weight | 362.39 |
| space group | *P-1* |
| $a$/Å | 9.4678(9) |
| $b$/Å | 10.7465(10) |
| $c$/Å | 17.3877(19) |
| $\alpha$/° | 88.938(7) |
| $\beta$/° | 80.003(7) |
| $\gamma$/° | 71.208(6) |
| $V$/Å$^3$ | 1648.1(3) |
| $Z$ | 4 |
| temperature (K) | 130(2) |
| radiation (λ, Å) | 0.71073 |
| ρ (calcd.) g cm$^{-3}$ | 1.461 |
| μ (Mo Kα), mm$^{-1}$ | 0.220 |
| θ max, deg. | 25.164 |
| no. of data collected | 137579 |
| no. of data | 5868 |
| no. of parameters | 470 |
| $R_1$ [$I > 2\sigma(I)$] | 0.0512 |
| $wR_2$ [$I > 2\sigma(I)$] | 0.1268 |
| $R_1$ [all data] | 0.0658 |
| $wR_2$ [all data] | 0.1333 |
| GOF | 1.082 |
| $R_{int}$ | 0.1420 |

**Table EV2.** Crystal, intensity collection, and refinement data.

| | **FKK6•EtOAc** |
|---|---|
| lattice | Monoclinic |
| formula | $C_{27}H_{26}N_2O_5S$ |
| formula weight | 490.56 |
| space group | $P2_1/n$ |
| $a$/Å | 7.7834(13) |
| $b$/Å | 19.087(3) |
| $c$/Å | 16.355(3) |
| α/° | 90 |
| β/° | 100.528(4) |
| γ/° | 90 |
| $V$/Å$^3$ | 2388.8(7) |
| $Z$ | 4 |
| temperature (K) | 130(2) |
| radiation (λ, Å) | 0.71073 |
| ρ (calcd.) g cm$^{-3}$ | 1.364 |
| μ (Mo Kα), mm$^{-1}$ | 0.178 |
| θ max, deg. | 27.601 |
| no. of data collected | 39284 |
| no. of data | 5140 |
| no. of parameters | 326 |
| $R_1$ [$I > 2\sigma(I)$] | 0.0426 |
| $wR_2$ [$I > 2\sigma(I)$] | 0.0816 |
| $R_1$ [all data] | 0.0814 |
| $wR_2$ [all data] | 0.0942 |
| GOF | 1.002 |
| $R_{int}$ | 0.0722 |

**Table EV3.**

| Organ | Vehicle n=6* | FKK6 n=6 |
|---|---|---|
| Heart | Minimal lymphoplasmacytic infiltrates (n=1) | – |
| Coronary Arteries/Aortic Root | – ** | – |
| Liver | Mild periportal inflammation (n=5), Hepatocyte necrosis (n=3) | Mild periportal inflammation (n=5) |
| Kidney | Degenerative tubules/vacuoles (rare) (n=6) | Degenerative tubules/vacuoles (rare) (n=6) |
| Lung | Mild leukocytes (n=3) | Mild leukocytes (n=2) |
| Stomach | – | – |
| Small Intestine | – | – |
| Large Intestine | – | – |
| Cecum | – | – |
| Colon | – | – |
| Spleen | Mild lymphoid hyperplasia (n=2) | Mild lymphoid hyperplasia (n=2) |
| Bone Marrow | – | – |

*One mouse was found deceased during day 10 cage inspection. All other mice examined were in good body condition with no minimal autolysis and no gross lesions. **Aortic root not visualized in (n=1) mouse.

**Table EV4.**

| Organ | Vehicle<br>n=6* | FKK6<br>n=6 |
|---|---|---|
| Heart | – | – |
| Coronary Arteries/Aortic Root | – ** | – |
| Liver | Mild perivascular lymphohistiolytic aggregates (n=3) | Mild perivascular lymphohistiocytic aggregates (n=4) |
| Kidney | Degenerative tubules/vacuoles (rare) (n=1) | – |
| Lung | Mild leukocytes (n=3) | Mild leukocytes (n=2) |
| Stomach | Focal neutrophilic aggregate (n=1) | – |
| Small Intestine | – | – |
| Large Intestine | – | – |
| Cecum | Moderate lymphoid hyperplasia (n=1) | – |
| Colon | – | – |
| Spleen | Mild lymphoid hyperplasia (n=2) | Mild lymphoid hyperplasia (n=2) |
| Bone Marrow | – | – |

#### Table EV5A.

#### S-Score (468 Kinase Screen) Results for FKK6

| Compound Name | Selectivity Score Type | Number of Hits | Number of Non-mutant Kinases | Screening Concentration (μM) | Selectivity Score (S-Score) |
|---|---|---|---|---|---|
| FKK6 | S(35) | 0 | 403 | 10 | 0 |
| FKK6 | S(10) | 0 | 403 | 10 | 0 |
| FKK6 | S(1) | 0 | 403 | 10 | 0 |

### Table EV5B.

#### Select Kinase Screen Results for FKK6

| Acetyltransferase | Substrate | FKK6 (data 1, Exp 1) | FKK6 (data 2, Exp 1) | FKK6 (data 1, Exp 2) | FKK6 (data 2, Exp 2) |
|---|---|---|---|---|---|
| p300 | Histone H3 | 100.32 | 97.73 | 98.72 | 95.38 |
| pCAF | Histone H3 | 100.32 | 101.68 | 104.61 | 104.47 |
| | | % Enzyme Activity to DMSO controls | | | |
| | | Positive controls used (IC50) C646, Anacardic acid (full data available upon request) | | | |

| Enzyme | FKK6 (data 1 | FKK6 (data 2, Exp 1) | FKK6 (data 1, Exp 2) | FKK6 (data 2, Exp 2) |
|---|---|---|---|---|
| PARP 1 | 99.18 | 98.25 | 91.38 | 91.03 |
| | | % Enzyme Activity to DMSO controls | | |
| | | Positive controls used (IC50) PJ34, PJ35 (full data available upon request) | | |

| Methyltransferase | Substrate | FKK6 (data 1, Exp 1) | FKK6 (data 2, Exp 1) | FKK6 (data 1, Exp 2) | FKK6 (data 2, Exp 2) |
|---|---|---|---|---|---|
| PRMT1 | Histone H4 | 86.43 | 88.77 | 78.47 | 81.3 |
| | | % Enzyme Activity to DMSO controls | | | |
| | | **Positive control is SAH, (S-(5'-Adenosyl)-L-homocysteine) (full data available upon request)** | | | |

| Histone Deacetylases | Substrate | FKK6 (data 1, Exp 1) | FKK6 (data 2, Exp 1) | FKK6 (data 1, Exp 2) | FKK6 (data 2, Exp 2) |
|---|---|---|---|---|---|
| HDAC1 | Peptide (Ac) | 105.36 | 101.46 | 89.76 | 86.7 |
| HDAC2 | Peptide (Ac) | 86.27 | 78.19 | 83.24 | 88.71 |
| HDAC3 | Peptide (Ac) | 122.38 | 116.9 | 114.05 | 107.82 |
| | | % Enzyme Activity to DMSO controls | | | |
| | | Positive controls used (IC50) Trichostatin A (full data available upon request) | | | |

| Enzyme | Substrate | FKK6 (data 1, Exp 1) | FKK6 (data 2, Exp 1) | FKK6 (data 1, Exp 2) | FKK6 (data 2, Exp 2) |
|---|---|---|---|---|---|
| SIRT1 | acetylated peptide | 100.18 | 100.03 | 99.84 | 94.47 |
| | | | % Enzyme Activity to DMSO controls | | |
| | | | Positive controls used (IC50) Suramin (full data available upon request) | | |

#### Table EV5C.

#### Select Kinase Screen Results for FKK5

**Top**

DiscoverX

**ALB002-01-p-00001 Study Results**

Table 1 - Matrix of Compound Screen for ALB002-01-p-00001

| Target | FKK5 | |
|---|---|---|
| Gene Symbol | %Ctrl @ 10000nM | %Ctrl @ 25000nM |
| CDK2 | 100 | 89 |
| CDK4 | 100 | 95 |
| CDK5 | 77 | 61 |
| ERK1 | 86 | 46 |
| ERK2 | 99 | 94 |
| ERK3 | 100 | 80 |
| ERK4 | 97 | 89 |
| ERK5 | 94 | 100 |
| ERK8 | 97 | 75 |
| GSK3A | 79 | 49 |
| GSK3B | 100 | 98 |
| IKK-alpha | 96 | 83 |
| JNK1 | 91 | 66 |
| JNK2 | 95 | 74 |
| PKAC-alpha | 100 | 85 |
| PKAC-beta | 100 | 92 |
| SGK1 | 79 | 71 |

%Ctrl Legend

| 0≤x<.1 | .1≤x<1 | 1≤x<10 | 10≤x<35 | x≥35 |
|---|---|---|---|---|

Confidential ALB002-01-p-00001 11/30/2018 5 of 6

**Bottom**

| Target | FKK5 | |
|---|---|---|
| Gene Symbol | %Ctrl @ 10000nM | %Ctrl @ 25000nM |
| CDK2 | 74 | 63 |
| CDK4 | 98 | 90 |
| CDK5 | 100 | 100 |
| ERK1 | 100 | 100 |
| ERK2 | 100 | 85 |
| ERK3 | 88 | 93 |
| ERK4 | 82 | 91 |
| ERK5 | 79 | 92 |
| ERK8 | 71 | 65 |
| GSK3A | 92 | 77 |
| GSK3B | 92 | 90 |
| IKK-alpha | 86 | 100 |
| JNK1 | 95 | 85 |
| JNK2 | 88 | 80 |
| PKAC-alpha | 100 | 95 |
| PKAC-beta | 57 | 48 |
| SGK1 | 100 | 89 |

%Ctrl Legend

| 0≤x<.1 | .1≤x<1 | 1≤x<10 | 10≤x<35 | x≥35 |
|---|---|---|---|---|

Confidential ALB003-01-p-00001 11/30/2018 5 of 6

#### REFERENCES

1 Abad-Zapatero, C. & Blasi, D. Ligand Efficiency Indices (LEIs): More than a Simple Efficiency Yardstick. *Molecular informatics* **30**, 122-132, doi:10.1002/minf.201000161 (2011).

2 Abad-Zapatero, C. *AtlasCBS: A Graphic Tool to Map the Content of Structure-Activity Databases*. (2016).

3 Sheldrick, G. M. A short history of SHELX. *Acta crystallographica. Section A, Foundations of crystallography* **64**, 112-122, doi:10.1107/s0108767307043930 (2008).

4 Sheldrick, G. M. SHELXT - Integrated space-group and crystal-structure determination. *Acta Crystallographica Section A* **71**, 3-8, doi:10.1107/S2053273314026370 (2015).

5 Sheldrick, G. M. TWINABS. *Version 2012/1. Georg-August-Universität Göttingen, Göttingen, Germany* (2012).

6 Chandran, A. & Vishveshwara, S. Exploration of the conformational landscape in pregnane X receptor reveals a new binding pocket. *Protein science : a publication of the Protein Society* **25**, 1989-2005, doi:10.1002/pro.3012 (2016).

7 Phillips, J. C. *et al.* Scalable molecular dynamics with NAMD. *J Comput Chem* **26**, 1781-1802, doi:10.1002/jcc.20289 (2005).

8 Darden, T. A. & Pedersen, L. G. Molecular modeling: an experimental tool. *Environ Health Perspect* **101**, 410-412 (1993).

9 Bussi, G. & Parrinello, M. Accurate sampling using Langevin dynamics. *Phys Rev E Stat Nonlin Soft Matter Phys* **75**, 056707, doi:10.1103/PhysRevE.75.056707 (2007).

10 Bussi, G., Donadio, D. & Parrinello, M. Canonical sampling through velocity rescaling. *J Chem Phys* **126**, 014101, doi:10.1063/1.2408420 (2007).

11 Hoover, W. G. Canonical dynamics: Equilibrium phase-space distributions. *Phys Rev A Gen Phys* **31**, 1695-1697 (1985).

12 Nosé, S. A molecular dynamics method for simulations in the canonical ensemble. *Molecular physics* **52**, 255-268 (1984).

13 Jones, G., Willett, P. & Glen, R. C. Molecular recognition of receptor sites using a genetic algorithm with a description of desolvation. *Journal of molecular biology* **245**, 43-53 (1995).

14 Jones, G., Willett, P., Glen, R. C., Leach, A. R. & Taylor, R. Development and validation of a genetic algorithm for flexible docking. *Journal of molecular biology* **267**, 727-748, doi:10.1006/jmbi.1996.0897 (1997).

15 Laskowski, R. A. & Swindells, M. B. LigPlot+: multiple ligand-protein interaction diagrams for drug discovery. *Journal of chemical information and modeling* **51**, 2778-2786, doi:10.1021/ci200227u (2011).

16 Fabian, M. A. *et al.* A small molecule-kinase interaction map for clinical kinase inhibitors. *Nature biotechnology* **23**, 329-336, doi:10.1038/nbt1068 (2005).

17 Karaman, M. W. *et al.* A quantitative analysis of kinase inhibitor selectivity. *Nature biotechnology* **26**, 127-132, doi:10.1038/nbt1358 (2008).

18 Horiuchi, K. Y. *et al.* Assay development for histone methyltransferases. *Assay Drug Dev Technol* **11**, 227-236, doi:10.1089/adt.2012.480 (2013).

19 Toporova, L., Macejova, D. & Brtko, J. Radioligand binding assay for accurate determination of nuclear retinoid X receptors: A case of triorganotin endocrine disrupting ligands. *Toxicology letters* **254**, 32-36, doi:10.1016/j.toxlet.2016.05.005 (2016).

20 Illes, P., Grycova, A., Krasulova, K. & Dvorak, Z. Effects of Flavored Nonalcoholic Beverages on Transcriptional Activities of Nuclear and Steroid Hormone Receptors: Proof of Concept for Novel Reporter Cell Line PAZ-PPARg. *J Agric Food Chem* **66**, 12066-12078, doi:10.1021/acs.jafc.8b05158 (2018).
